# CRISPR/dCas-mediated counter-silencing – Reprogramming dCas proteins into antagonists of xenogeneic silencers

**DOI:** 10.1101/2024.08.29.610263

**Authors:** Johanna Wiechert, Biel Badia Roigé, Doris Dohmen-Olma, Hindra, Xiafei Zhang, Roberto G. Stella, Marie A. Elliot, Julia Frunzke

## Abstract

Lsr2-like nucleoid-associated proteins function as xenogeneic silencers (XSs) inhibiting expression of horizontally acquired, AT-rich DNA in actinobacteria. Interference by transcription factors can lead to counter-silencing of XS target promoters, but relief of this repression typically requires promoter engineering. In this study, we developed a novel CRISPR/dCas-mediated counter-silencing (CRISPRcosi) approach by using nuclease-deficient dCas enzymes to counteract the Lsr2-like XS protein CgpS in *Corynebacterium glutamicum* or Lsr2 in *Streptomyces venezuelae*. Systematic *in vivo* reporter studies with dCas9 and dCas12a and various guide RNAs revealed effective counter-silencing of different CgpS target promoters in response to guide RNA/dCas DNA binding – independent of promoter sequence modifications. The most prominent CRISPRcosi effect was observed when targeting the CgpS nucleation site, an effect that was also seen in *S. venezuelae* when targeting a known Lsr2 nucleation site within the chloramphenicol biosynthesis gene cluster. Analyzing the system in *C. glutamicum* strains lacking the XS protein CgpS revealed varying strengths of counteracting CRISPR interference (CRISPRi) effects based on the target position and strand. Genome-wide transcriptome profiling in sgRNA/dCas9 co-expressing *C. glutamicum* wild-type strains revealed high counter-silencing specificity with minimal off-target effects. Thus, CRISPRcosi provides a promising strategy for the precise upregulation of XS target genes with significant potential for studying gene networks as well as for developing applications in biotechnology and synthetic biology.

**IMPORTANCE:** Lsr2-like nucleoid-associated proteins act as xenogeneic silencers (XSs), repressing the expression of horizontally acquired, AT-rich DNA in actinobacteria. The targets of Lsr2-like proteins are very diverse, including prophage elements, virulence gene clusters and biosynthetic gene clusters. Consequently, the targeted activation of XS target genes is of interest for fundamental research and biotechnological applications. Traditional methods for counter-silencing typically require promoter modifications. In this study, we developed a novel CRISPR/dCas-mediated counter-silencing (CRISPRcosi) approach, utilizing nuclease-deficient dCas enzymes to counteract repression by Lsr2-like proteins in *Corynebacterium glutamicum* and *Streptomyces venezuelae*. The strongest effect was observed when targeting the Lsr2 nucleation site. Genome-wide transcriptome profiling revealed high specificity with minimal off-target effects. Overall, CRISPRcosi emerges as a powerful tool for the precise activation of genes silenced by xenogeneic silencers, offering new opportunities for exploring gene networks and advancing biotechnological applications.

## INTRODUCTION

The acquisition of new genetic elements via horizontal gene transfer (HGT) represents a major driving force of prokaryotic evolution (1, 2). It can provide favorable phenotypic traits (1, 3); however, the unregulated expression of foreign genes can be energetically and physiologically costly for the recipient cell (2, 4–8). Consequently, bacteria evolved various immune systems to protect from invading DNA, including defense strategies against bacteriophages (9–13). Destructive, nuclease-based defense mechanisms, such as restriction modification and CRISPR/Cas systems catalyze the targeted degradation of foreign DNA (14–16).

In contrast to these destructive mechanisms, xenogeneic silencing fosters the acquisition of foreign DNA into the host chromosome by promoting tolerance (17, 18). The molecular basis of this trait are xenogeneic silencers (XSs), specialized nucleoid-associated proteins (NAPs). XS proteins are widely distributed and have convergently evolved across prokaryotic phylogenetic clades, emphasizing their evolutionary importance. They are currently classified into four families encompassing H-NS XS proteins of proteobacteria (19–21), Rok from bacilli species (22, 23), MvaT/U like XS proteins from the *Pseudomonodales* order (24), and Lsr2/Lsr2-like XS in *Actinobacteria* (25, 26). Although XS proteins differ in their DNA interaction mode, they commonly recognize and bind horizontally acquired DNA with higher AT content than their resident genomes (2, 18). By forming higher-order nucleoprotein complexes, XS proteins silence their target genes, e.g., by occlusion of the RNA polymerase (27, 28).

Lsr2 and Lsr2-like XS proteins play important roles in the biotechnologically and medically important phylum of *Actinobacteria*. In *Corynebacterium glutamicum*, the Lsr2-like XS protein CgpS plays an essential role as silencer of the cryptic prophage element CGP3 by preventing its entrance into the lytic cycle (4). In contrast, Lsr2 XS proteins in *Mycobacterium tuberculosis* and *Streptomyces* species are key players in silencing the expression of virulence gene (26, 29, 30) and specialized metabolic clusters (31), respectively. Consequently, these XS proteins can be considered as highly relevant targets for drug development (32) and for the identification of new bioactive compounds (31). These examples highlight the importance of understanding the rules underlying Lsr2-mediated xenogeneic silencing and of developing applicable counter-silencing mechanisms allowing for the precise and controlled reactivation of silenced gene clusters. In several systematic counter-silencing studies, transcription factor-(TF) binding within silenced promoter regions was demonstrated to interfere with the nucleoprotein complex, leading to promoter reactivation (33–36). In our previous study, we systematically inserted TF operator sequences at different positions into a set of CgpS target promoters in *C. glutamicum* and demonstrated highest TF-mediated counter-silencing efficiency close to the putative CgpS nucleation site in the DNA region of highest AT content (33). Overall, these TF-based counter-silencer approaches allow for the targeted reactivation of silenced genes. However, they all depend on native or artificially inserted TF operator sequences (33–36).

With a view to applications, this sequence dependency might represent a major constraint. With this motivation, we wanted to develop a counter-silencing system that allows for the targeted reactivation of silenced genes without the need to modify the silenced DNA sequence. In contrast to TF binding, small guide RNAs can direct (d)Cas enzymes precisely to DNA positions, thereby providing target specificity. We hypothesized that guide RNAs and dCas enzymes represent promising candidates for a **CRISPR**/dCas-mediated **co**unter-**si**lencing (CRISPRcosi) approach to counteract xenogeneic silencing independent of TF binding by using ‘dead‘ Cas proteins (dCas9: D10A and H840A (37, 38) and dCas12a: (D917A, E1006A) (39, 40)). Usually, these guide RNAs and dCas enzymes are the key components of CRISPR interference (CRISPRi) approaches, a well-established gene knockdown platform (38, 41). However, guide RNA positions outside of the core promoter region reveal a significant loss in repression efficiency in CRISPRi studies (38, 41). In the CRISPRcosi approach, optimal guide RNA/dCas target sites should feature low CRISPRi effects combined with strong interference effect, leading to promoter activation by CRISPRcosi.

In this study, we set out to systematically investigate the potential of CRISPRcosi as a modular approach to reactivate XS target genes by using the Lsr2-like XS protein CgpS as a model. To the best of our knowledge, this is the first regulatory system that combines a guide RNA and a dCas protein to enhance promoter activity without requiring an engineered dCas variant, as commonly used in CRISPR activation (CRISPRa) approaches (41, 42). In vivo reporter studies with a set of different CgpS target promoters and guide RNAs demonstrated that the binding position of the guide RNA/dCas complex determined the balance between counter-silencing and adverse CRISPRi effects. CgpS counter-silencing was most effective at the position of maximal CgpS binding (close to its putative nucleation site) but without strict strand specificity. This counter-silencing approach was also found to be applicable for Lsr2 in *Streptomyces venezuelae*, supporting the broad application across the Actinobacteria.

## RESULTS

### A new CRISPR/dCas application beyond CRISPR interference – Using dCas9 to reactivate targets of xenogeneic silencers

In this work, we present CRISPRcosi, a CRISPR/dCas-based counter-silencing system. In contrast to previously established transcription factor (TF)-based counter-silencing approaches, CRISPRcosi is independent of TF binding sites (Figure 1). Here, single guide RNAs (sgRNAs) were used to direct ‘dead’ Cas9 (dCas9) (37) to target promoters of the Lsr2-like protein CgpS. These silenced promoters typically feature a distinct drop in GC content and an inversely correlated CgpS binding peak covering several hundred nucleotides (4) (Figure 1A). Comparable to the TF-based approach, binding of the sgRNA/dCas9 complex close to the CgpS nucleation site might interfere with the silencer-DNA complex, leading to counter-silencing and transcription of the associated gene(s). The integrity of the silencer-DNA complex in response to such interference effects is usually not completely disrupted, meaning that the promoters are only partially reactivated (33). To monitor promoter activities, prophage promoters were fused to the reporter genes *venus* or *eyfp* (Figure 1B). The sgRNA-DNA complementarity, mediated by the sgRNA spacer sequence, in combination with an NGG protospacer adjacent motif (PAM) sequence, served as checkpoints to guide dCas9 to specific DNA positions and strands (43). sgRNAs were designed to bind the template or the coding strand of the DNA at different positions within the CgpS target promoters (Figure 1C). As proof-of-concept, microscopy studies with *C. glutamicum* cells carrying a reporter system consisting of the CgpS target promoter P_cg1974_ fused to the *eyfp* reporter gene demonstrated the potential of the CRISPRcosi approach. A strong yellow fluorescence signal only present in cells co-expressing *dcas9* and the specific sgRNA-CS3 confirmed the functionality of this activating approach and the strict dependency on both components (Figure 1D).

**Figure 1.**
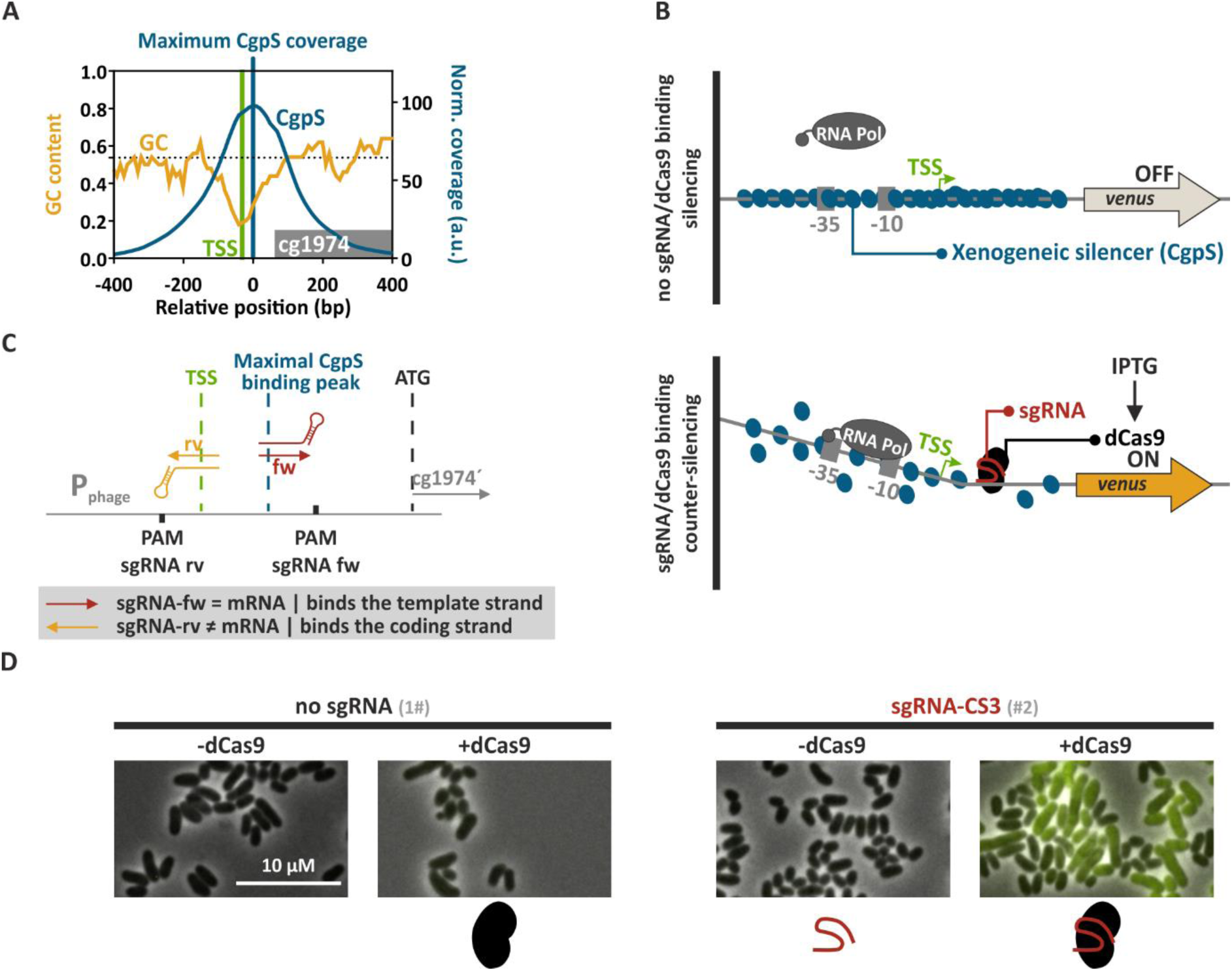
Principle of CRISPRcosi. (A) Typical inverse correlation of CgpS coverage (blue line) and GC profiles (orange line) of a representative CgpS target promoter (for gene cg1974 (grey box) from the CGP3 prophage region). Shown are the position of maximal CgpS coverage (blue line (4)), TSS (green) and the average *C. glutamicum* ATCC 13032 GC content (53.8%, dotted black line) (44). (B) Schematic overview of a CgpS target promoter and the principle of xenogeneic silencing and CRISPRcosi. (C). Design of single guide RNAs and corresponding adjacent NGG PAM sequences (black box). sgRNAs consisted of 20-nucleotide target specific spacer sequence and an optimized chimeric (d)Cas9 binding hairpin (45). (D) Representative fluorescence microscopy images of *C. glutamicum* wild-type cells with an integrated P_cg1974_-*eyfp* sequence (*C. glutamicum* ATCC 13032::P_cg1974_-*eyfp* (46)) that harboured either no sgRNA (#1) or the sgRNA-CS3 encoding sequence (#2), in combination with the *dcas9* gene. Cells were cultivated in the presence (+dCas9) or absence of 200 µM IPTG (-dCas9). The scale bar: 10 µm. For plasmid IDs (#number), refer to Table S2.

### Competing effects of CRISPRi and CRISPRcosi at CgpS target promoters

After demonstrating that the mechanism of CRISPRcosi allowed the targeted activation of CgpS silenced genes, we set out to systematically assess the opportunities and constraints of this approach. To prevent the induction of prophage genes, we used the prophage-free *C. glutamicum* Δphage::P*_cgpS_*-*cgpS* (33) derivative with a reintegrated copy of the *cgpS* gene as our platform strain. All plasmid IDs (#number) given in the figures and text are referenced in Table S2. The sgRNAs were designed to bind to different DNA strands and positions directly adjacent to NGG PAM sequences in the P_cg1974_ promoter sequence. While the homologous region on the coding strand of sgRNA-CS1 overlapped with the putative CgpS binding motif as well as with the dominant transcriptional start site (TSS) (33), sgRNA-CS3 bound the template strand in the region of maximal CgpS binding coverage seven nucleotides downstream of the CgpS nucleation site (Figure 2A). The backscatter-normalized specific Venus fluorescence signals (specific fluorescence=total Venus fluorescence/backscatter) measured after 20 h of cultivation reflected the promoter activities of strains with different sgRNA-encoding sequences and the presence and absence of the IPTG-inducible *dcas9* gene. Here, only the co-expression of *dcas9* and sgRNA-CS3, but not sgRNA-CS1, resulted in an IPTG-correlated stepwise increase in promoter activity, demonstrating success with CRISPRcosi in a dCas9-dose dependent manner (Figure 2B, Figure S1).

**Figure 2.**
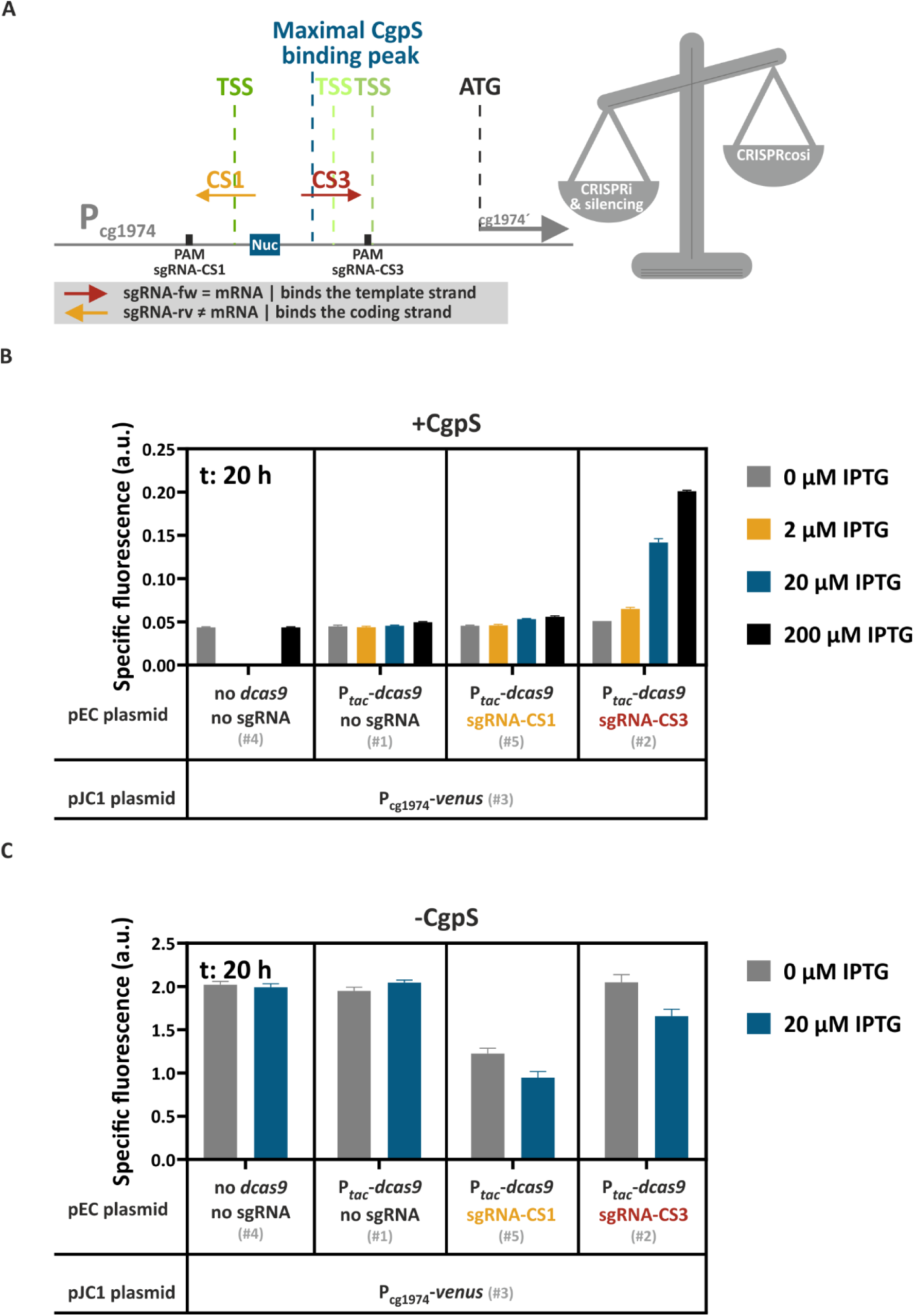
The balance between CRISPR interference and CRISPRcosi effects determined the activity of the CgpS target promoter. (A). Schematic overview of the two sgRNAs, sgRNA-CS1 and sgRNA-CS3 targeting P_cg1974_. Green vertical lines, TSSs (ranking of TSS enrichment scores: dark green>light green (33)); blue box, CgpS nucleation site (Nuc) (33). (B and C) Backscatter-normalized reporter outputs (t: 20 h; specific Venus fluorescence) of prophage-free, *cgpS* expressing *C. glutamicum* strains (Δphage::P*_cgpS_*-*cgpS*, (33), B) and derivatives lacking CgpS (Δphage, (**48**), C). All strains harboured the pJC1-P_cg1974_-*venus* plasmid (#3) and differed in their genetic repertoire of sgRNAs-encoding sequences and the presence/absence of the *dcas9* gene, depending on the co-transferred pEC-based plasmid (no sgRNA and no *dcas9* (#4), no sgRNA but *dcas9* (#1), sgRNA-CS1 and *dcas9* (#5), sgRNA-CS3 and *dcas9* (#2)). (n=3). Plasmid IDs (#number) refer to Table S2.

The combination of sgRNAs and dCas enzymes are commonly used to inhibit gene expression in CRISPR interference (CRISPRi) approaches (38). Depending on the position, binding of the sgRNA/dCas9 complex can sterically prevent transcriptional initiation or elongation leading to repression (38, 41, 47). The presented CRISPRcosi approach uses the same components, suggesting that binding of the sgRNA/dCas9 complex at specific positions might negatively affect P_cg1974_ promoter activity. Prophage promoters like P_cg1974_ are typically constitutively and strongly active in the absence of the xenogeneic silencer CgpS (33). To monitor putative CRISPRi effects, P_cg1974_-driven reporter outputs of prophage-free, CgpS-lacking *C. glutamicum* Δphage cells (48) were tested in the context of different sgRNA/dCas9 combinations. Even in the absence of IPTG, significantly reduced promoter activities were observed in cells expressing sgRNA-CS1, likely caused by leaky P*_tac_*-derived dcas9 expression. In contrast, sgRNA-CS3 showed only minor CRISPRi effects (Figure 2C). These findings aligned with results from CRISPRi studies, which demonstrated that CRISPRi performance is determined by the sgRNA/dCas9 target position, where effects ranged from neutral effects to highest effects when binding in close proximity of the TSS (38, 41, 49–52). sgRNA-CS3 binds the template DNA strand downstream of the dominant TSS, a position that is generally less sensitive to dCas9-mediated CRISPRi effects (38, 41). In conclusion, the balance between CRISPRi and counter-silencing effects determined the overall activity of the P_cg1974_ promoter in the presence of CgpS. With respect to CRISPRcosi, the counter-silencing effects had to be stronger than any CRISPRi effects, resulting in promoter activation.

### CRISPRi is dominant over CRISPRcosi effects

Having demonstrated that binding of the CgpS nucleation site-specific sgRNA-CS3/dCas9 complex counteracted CgpS silencing at the prophage promoter P_cg1974_, we next set out to determine the relative effect of CRISPRi compared with CRISPRcosi. Therefore, we applied a double plasmid system (pEC- and pJC1-derivates) to compare P_cg1974_-*venus* reporter outputs of cells differing in their set of sgRNAs and *dcas9* availability. Promoter activities revealed that two copies of sgRNA-CS3 did not result in a synergistic effect, suggesting that the amount of sgRNA-CS3 derived from a single copy did not limit counter-silencing efficiencies (Figure 3A). Second, the activating effect of the sgRNA-CS3/dCas9 complex was completely abolished in the presence of sgRNA-CS1, as shown by the very low P_cg1974_-driven reporter signal (Figure 3A). These results indicate that the sgRNA-CS1/dCas9-mediated CRISPRi effect was dominant over the sgRNA-CS3/dCas9-dependent CRISPRcosi effect. The PAM-dependency of CRISPRcosi was confirmed by the lack of responsiveness of the modified P_cg1974 ΔPAM_ promoter, which lacked the sgRNA-CS3-specific PAM sequence (Figure 3).

**Figure 3.**
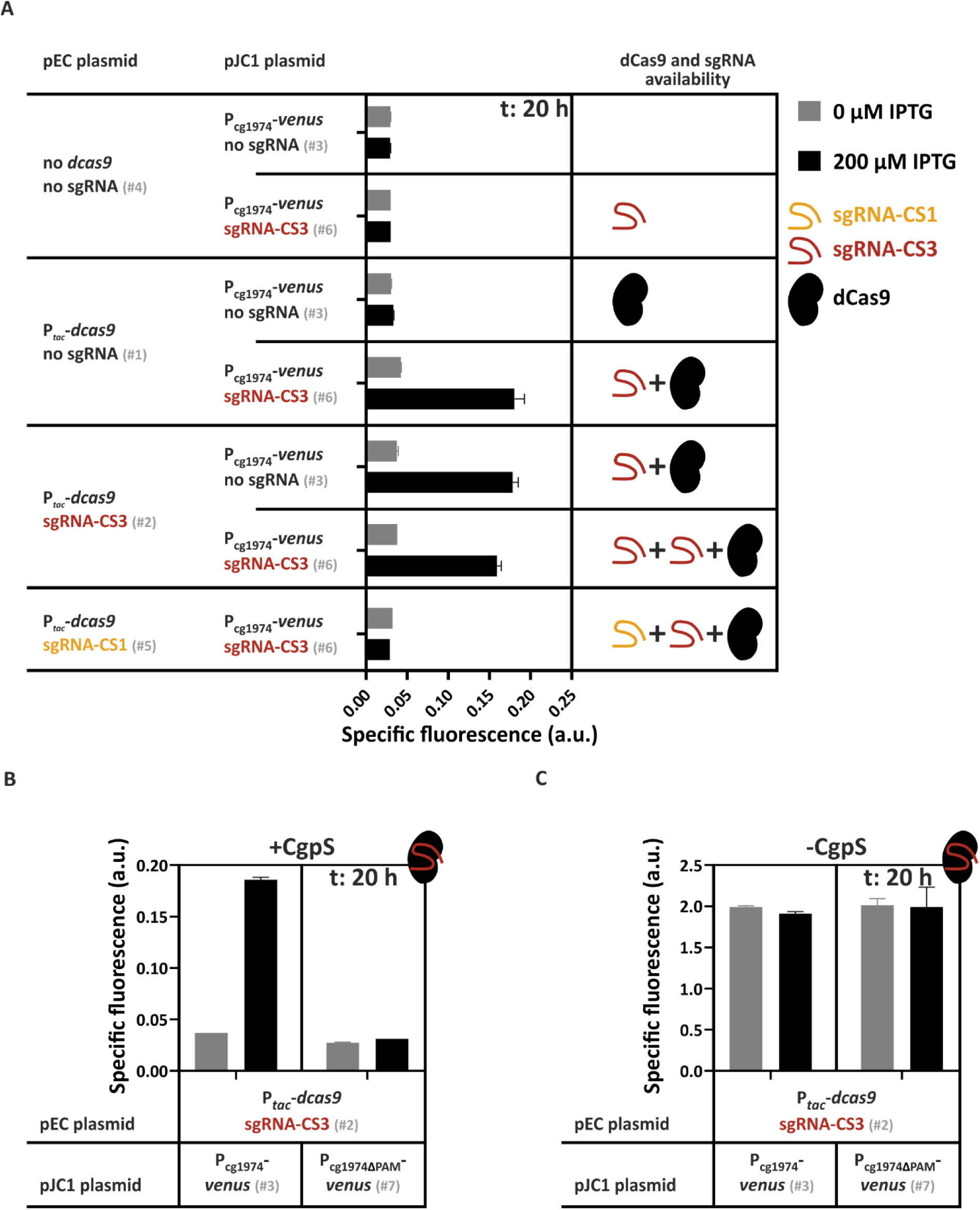
CRISPRcosi can be counteracted by CRISPRi effects and is PAM-dependent. (A) Backscatter-normalized reporter outputs (t: 20 h; specific Venus fluorescence) driven by the CgpS target promoter P_cg1974_ in different strains. (A) Prophage-free, *cgpS* expressing *C. glutamicum* strains (Δphage::P*_cgpS_*-*cgpS* (33)) containing the P_cg1974_-*venus* reporter system, but differing in their *dcas9* and sgRNA repertoire. (B, C) Prophage-free, *cgpS* expressing *C. glutamicum* strains (Δphage::P*_cgpS_*-*cgpS*, (33), B) and cells lacking the *cgpS* gene (Δphage, (48), C) were pEC plasmid-based equipped with the *dcas9* gene and the sgRNA-CS3 encoding sequence (#2). Additionally, cells harboured the pJC1 plasmid containing the native P_cg1974_ promoter (#3) or the P_cg1974ΔPAM_ derivative with a deleted sgRNA-CS3 PAM sequence (#7) in front of the *venus* reporter gene. (n=3). Plasmid IDs (#number) refer to Table S2.

### The efficiency of CRISPRcosi depends on the binding position of the sgRNA/dCas9 complex

The above-mentioned results indicated that the CRISPRcosi mechanism was most effective when the sgRNA/dCas9 complex did not interfere with transcription but bound to DNA regions that are effective in counteracting CgpS silencing (*e.g.,* CgpS nucleation site). To improve our understanding of the rules underlying this mechanism, we systematically screened sgRNAs that guide dCas9 to different positions of the CgpS target promoters P_cg1974_ and P_cg2014_ (Figure 4). We found both promoters were activated in the presence of sgRNAs directing dCas9 to the DNA sequences with the lowest GC content. In addition to sgRNA-CS3, sgRNA-33 counteracted CgpS silencing at the prophage promoter P_cg1974_ (Figure 4C). Interestingly, both sgRNAs were designed to bind the DNA template strand with a shift of only one nucleotide and overlapped with the annotated position of maximal CgpS binding (4). No other sgRNA induced prophage promoter activity, even when partly overlapping in binding position or bound in proximity.

**Figure 4.**
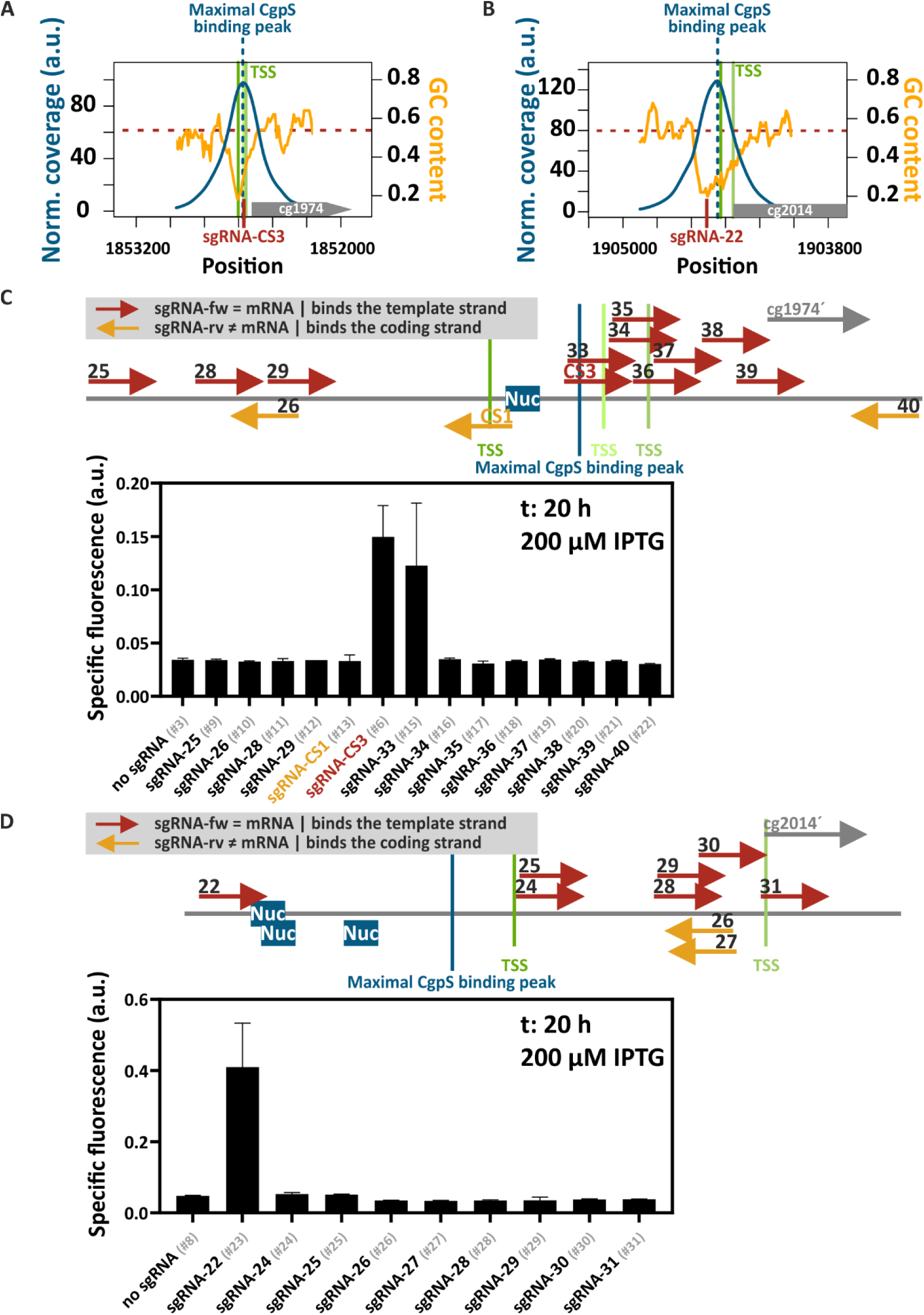
Targeting the CgpS nucleation site results in efficient CRISPRcosi. (A, B) Inverse correlation of CgpS coverage (blue line) and GC profiles (orange line) of the CgpS target promoters P_cg1974_ (A) and P_cg2014_ (B). Green lines, TSSs (ranking of TSS enrichment scores: dark green > light green (33)). (C, D) Target position and DNA strand of the different sgRNAs designed for dCas9 guiding to the CgpS target promoter P_cg1974_ (C) and P_cg2014_ (D) (red and orange arrows). Blue boxes (Nuc), CgpS nucleation sites (33). Bar plots represent backscatter-normalized reporter outputs (t: 20 h; specific Venus fluorescence) of prophage-free, *cgpS* expressing *C. glutamicum* strains (Δphage::P*_cgpS_*-*cgpS*, (33)) harbouring the #1 plasmid needed for IPTG-inducible *dcas9* expression. The second pJC1-based plasmids contained the P_cg1974_-*venus* (C) or the P_cg2014_-*venus* reporter construct (D) and either no sgRNA encoding sequence or one of the different sgRNA encoding sequences. (n=3). Plasmid IDs (#number) are referenced in Table S2.

In the case of P_cg2014,_ only sgRNA-22 led to CRISPRcosi (Figure 4D). sgRNA-22 was designed to bind the template DNA strand in the area of lowest GC content at the postulated CgpS nucleation sites (33) upstream of the position of maximal CgpS coverage (4) (Figure 4B,D). Dose-response studies using increments of IPTG to induce *dcas9* expression demonstrated a stepwise increase in P_cg2014_ promoter activity (Figure S1B). In summary, the screening results clearly demonstrated that CRISPRcosi is context dependent. We hypothesize that multiple sgRNA binding position-related factors hampered activation of the CgpS target promoters, including counteracting CRISPRi effects, the cooperative binding behavior of the silencer-DNA complex and the low frequency of the guanine-rich, NGG (d)Cas9 PAM sequence within AT-rich promoters. These constraints highlighted the need to increase the flexibility of this approach, for example by implementing alternative dCas enzymes like dCas12a that recognize AT-rich PAM sequences.

### The AT-rich PAM sequence facilitates dCas12a-dependent CRISPRcosi at AT-rich promoters

dCas12a (dFnCpf1;) (39, 40) represents a promising candidate to improve guide RNA selection within promoter regions, which are generally more AT-rich than non-promoter regions. In contrast to the downstream located, guanine-rich NGG (d)Cas9 PAM sequence, (d)Cas12a recognizes a (T)TTN PAM sequence upstream of the target (53–55) (Figure 5). We systematically assessed the effect of dCas12a-specific guide RNAs (crRNAs) in the context of CRISPR/dCas12a-mediated CgpS counter-silencing. Comparison of nine crRNAs across a broad range of binding positions in P_cg1974_ revealed two effective crRNAs, both designed to bind the coding strand at the annotated TSSs (33) (Figure S2A). While crRNA-9 had lower promoter activation capacities, crRNA-7-mediated guiding of dCas12a to the dominant TSS resulted in strong counter-silencing effects (Figure 5, Figure S2A). The other seven crRNAs were unable to counteract CgpS silencing, highlighting that only specific promoter regions were susceptible to CRISPR/dCas12a-mediated counter-silencing (Figure S2A), as observed for dCas9.

**Figure 5.**
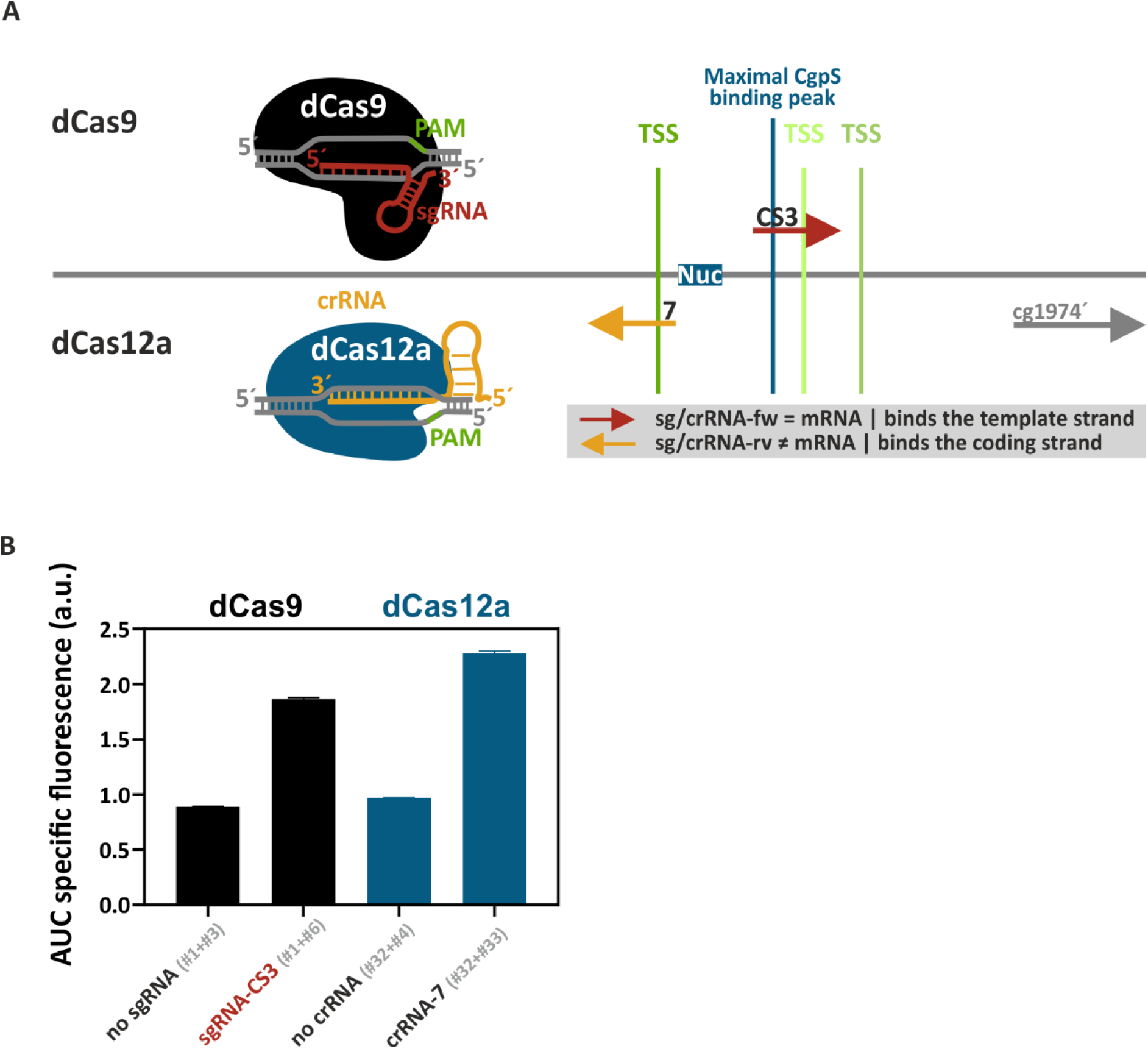
dCas9 and dCas12a both mediate efficient CRISPRcosi. (A) Arrows in the graphical representation show the target position and DNA strand of the different dCas9-guiding sgRNAs (top) and dCas12a-guiding crRNAs (bottom). Blue box (Nuc), CgpS nucleation site (33). (B) The bar plots represent the integrals over 24 h of cultivation (area under curve, AUC) of the backscatter-normalized reporter outputs (specific Venus fluorescence, shown in Figure S3A). Prophage-free, *cgpS* expressing *C. glutamicum* strains (Δphage::P*_cgpS_*-*cgpS*,(33)) harbouring the P_cg1974_-*venus* sequence and either the *dcas9* gene combined with the sgRNA-CS3-encoding sequence (#1+#6) or the *dcas12a* gene in concert with the crRNA-7 encoding sequence (#32+#33) were compared. Constructs lacking any guide RNAs encoding sequences (dCas9: #1+#3; dCas12a: #32+#4) served as controls. (n=3). Plasmid IDs (#number) refer to Table S2.

By using the most effective P_cg1974_ specific guide RNA candidates for each system (sgRNA-CS3 and crRNA-7), we directly compared the dCas9- and dCas12a-mediated CRISPRcosi utility (Figure 5, Figure S3A). Here, the crRNA-7/dCas12a approach showed slightly higher activation in comparison to the dCas9/sgRNA-CS3 system (Figure 5). In addition, the lower toxicity of dCas12a (54, 56) improved plasmid stability and allowed for the constitutive expression of the dCas enzyme in *C. glutamicum*, yielding strong and homogeneous counter-silencing effects. In contrast, *dcas9*-expressing cells often exhibited heterogeneous signal distribution, which was likely a consequence of the noisy, IPTG-inducible LacI-P*_tac_* expression system used to initiate *dcas9* transcription (Figure S3B).

### Guide RNA/dCas complex binding counteracted CgpS silencing at different prophage promoters

To evaluate the applicability of the CRISPRcosi mechanism as a tool to modulate gene expression of prophage genes, we tested our system with a set of 11 CgpS targets and both dCas enzymes. We analyzed promoter activities in prophage-free *C. glutamicum* cells (Δphage::P*_cgpS_*-*cgpS*, (33)) co-expressing *cgpS*, *dcas9* or *dcas12a*, and either a suitable guide RNA or no sg/crRNA (Figure S4). Ratios greater than one, for backscatter-normalized Venus reporter outputs of cells expressing a guide RNA and strains without any sg/crRNA, demonstrated counter-silencing. Values below one were indicative of CRISPRi effects (Figure 6). In summary, our dCas9 approach successfully yielded significantly increased (by >2fold) activity for the prophage promoters P_cg1974_, P_cg2014_ and P_cg2032_. With a 25-fold increase in promoter activity, cg2014_crRNA-2 was the top performing candidate for CRISPRcosi efficiencies (Figure 5, Figure S2).

**Figure 6.**
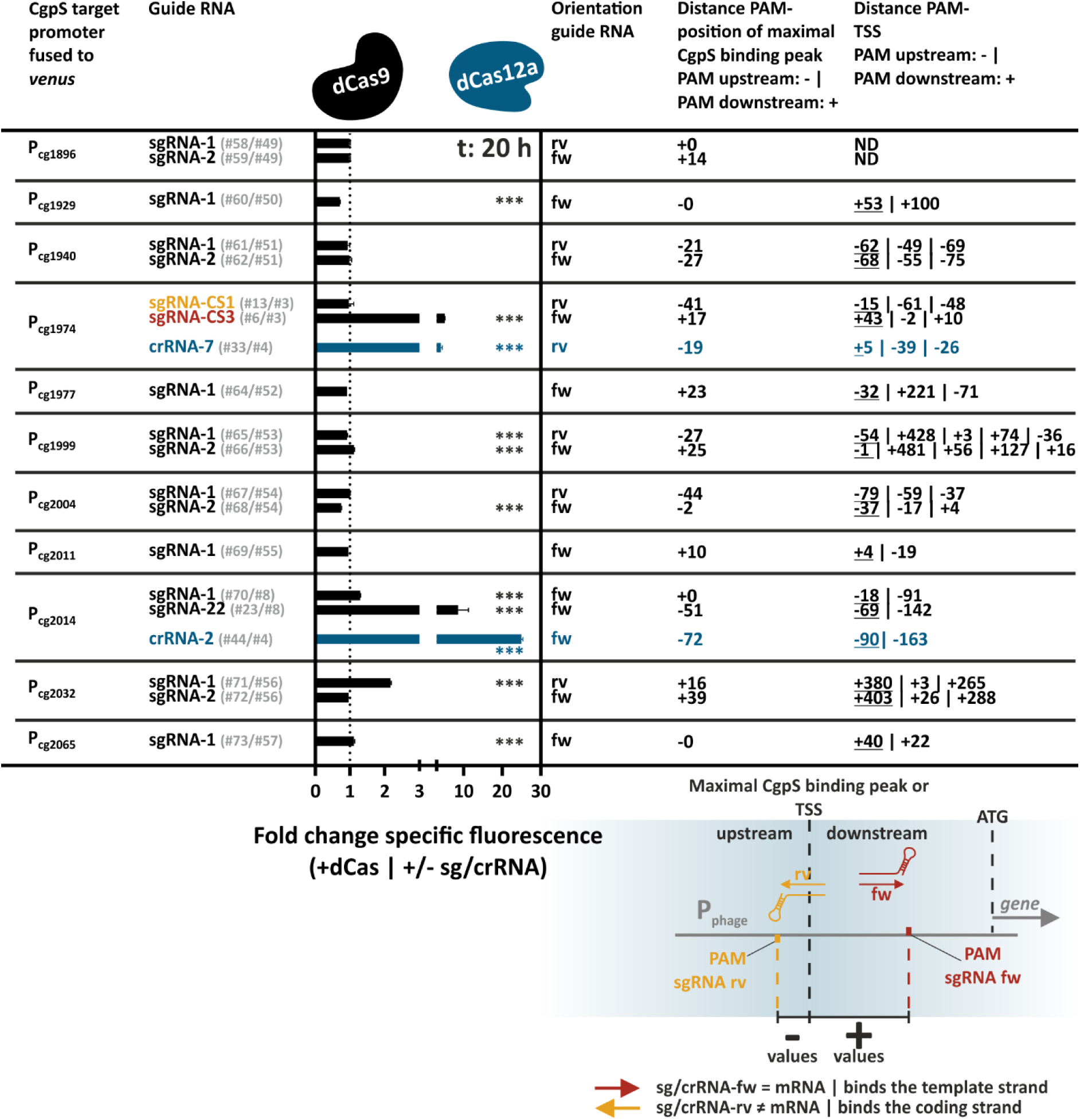
CRISPRcosi counteracts CgpS-mediated silencing at different prophage promoters. Backscatter-normalized fold changes of reporter outputs (t: 20 h; specific Venus fluorescence) of cells with and without guide RNA encoding sequences using dCas9 (black bars) or Cas12a (blue bars). Prophage-free, *cgpS* and *dcas9* or *dcas12a* co-expressing *C. glutamicum* cells (Δphage::P*_cgpS_*-*cgpS*, (33)) served as platform strain. *dcas9* expression was mediated by plasmid #1. sgRNA-free strains were equipped with pJC1-plasmids containing the sequences of the different prophage promoters fused to the reporter gene *venus*. P_J23319_-mediated expression of specific sgRNAs was achieved by inserting the guide RNA encoding sequences into the pJC1-plasmids (see Table S2). The orientation of the guide RNAs (fw: forward, sg/crRNA binds the template strand; rv: reverse, sg/crRNA binds the coding strand) and the distances between the positions of maximal CgpS binding/TSSs and the different GG or TTT PAM sequences in nucleotides are given. The value for the TSS with the highest enrichment score is underlined and all others are ordered according to this ranking (TSS from (33)). The scheme exemplarily represents the relative position of dCas9-guiding sgRNAs with respect to the TSS/maximal CgpS binding peak. (n=3). Significantly up- and downregulated values are marked by asterisks (unpaired t-test, ***two-tailed *p*-value <0.005). Time-resolved representation of the backscatter-normalized specific fluorescence values of all strains are given in Figure S4. Plasmid IDs (#number) can be found in Table S2.

Comparable to the sgRNA-22/dCas9 targeting cg2014, crRNA-2 facilitated counter-silencing by directing dCas12a to the template DNA strand at the putative CgpS nucleation site upstream of the P_cg2014_ core promoter region (Figure 6, Figure S2B). Unexpectedly, crRNA-3 targeted the opposite coding strand and failed to induce counter-silencing of P_cg2014_, although its binding region perfectly matched that of cg2014_crRNA-2 (Figure S2B). This demonstrated that the target DNA strand was relevant for dCas12a-based CRISPRcosi efficiency. Previous work has suggested that dCas9 binding to the coding strand leads to stronger CRISPRi effects (38, 41), while dCas12a-based gene repression is more efficient when targeting the template strand (39). Therefore, our results suggested that CRISPRcosi might be favored when CRISPRi effects were disfavored. This hypothesis was also supported by the point that all three counter-silencing inducing sgRNAs guide dCas9 to positions outside the core promoter (Figure 6), thereby avoiding the region generally considered to be most sensitive for CRISPRi (38, 41). In conclusion, we observed a trend that binding outside of the core promoter region and/or to the less efficient DNA strand with respect to CRISPRi, increased the likelihood of achieving CRISPRcosi. The implementation of additional dCas enzymes expanded the positional flexibility and facilitated the avoidance of counteracting CRISPRi effects.

### CRISPRcosi produces almost no off-target effects

To assess potential off-target effects of the CRISPRcosi approach, genome-wide transcriptome analysis of our CRISPRcosi approach was performed with *C. glutamicum* wild-type cells co-expressing *dcas9* and either P_cg1974_-specific sgRNA-CS3 or sgRNA-CS1 (Figure 7, Figure S5, Table S5). While sgRNA-CS3 facilitated CRISPRcosi, sgRNA-CS1 served as negative control in this context. (Figure 2). In comparison to the sgRNA-CS1 reference group, 14 genes were significantly upregulated (FDR-*p*-values<0.05; log_2_ fold change>1) in the sgRNA-CS3 samples (Figure 7, Figure S5), while one gene was downregulated (FDR-*p*-values<0.05; log_2_ fold change<1) (Figure 7A,B, Figure S5A). By using the Cas-OFFinder online tool (http://www.rgenome.net/cas-offinder/) (57), no sgRNA-CS3 or -CS1 off-target was associated with these genes (Table S6).

**Figure 7.**
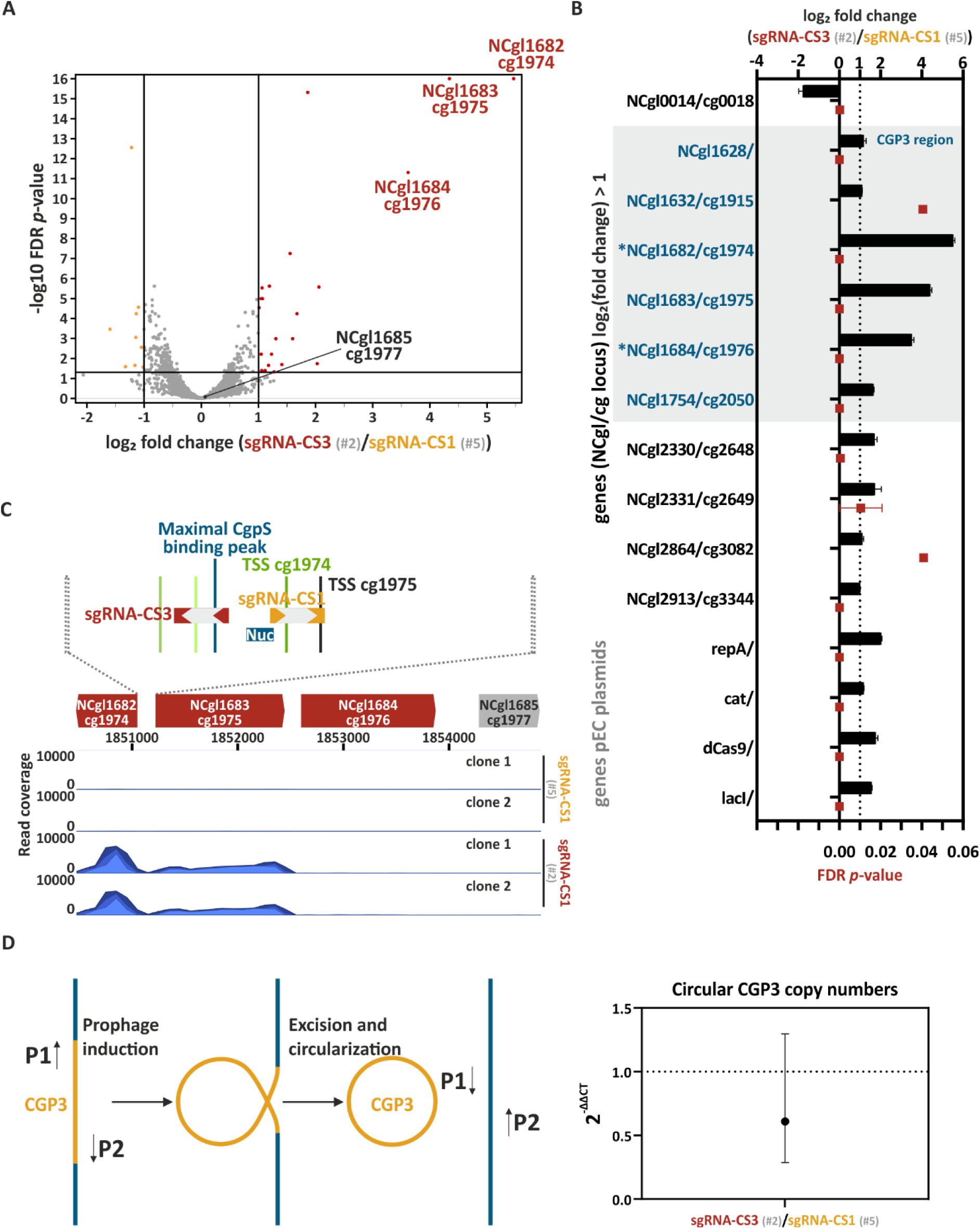
Investigation of potential off-target effects of CRISPRcosi. Genome-wide analysis of gene expression (A-C) and potential induction/excision of the CGP3 prophage (D) of *C. glutamicum* wild-type cells co-expressing *dcas9* and sgRNA-CS3 (#2) in comparison with control cells co-expressing *dcas9* and sgRNA-CS1 (#5). The given cg and NCgl loci numbers both refer to the strain *C. glutamicum* ATCC 13032 (cg: prophage-free strain, NCBI reference sequence NC_022040.1; NCgl: wild-type strain, NCBI reference sequence NC_003450.3). (A) Red and orange dots in the volcano plot represent significantly up- and downregulated genes (FDR-*p*-values<0.05 and |log_2_ fold change|>1), respectively, in cells expressing sgRNA-CS3 (clone 2) in comparison to cells expressing sgRNA-CS-1 (clone 1+2) (clone 1 is shown in Figure S5A). Non-significant hits are given as grey dots (FDR-*p*-values >0.05 and/or |log_2_ fold change| <1), significance thresholds are represented by black lines. (B) The bar plot represents all genes that were significantly up- and downregulated in sgRNA-CS3 cells compared to sgRNA-CS1 samples (FDR-*p*-values <0.05 and |log_2_ fold change (sgRNA-CS3/sgRNA-CS1)|>1). Black bars and red squares represent the means of log_2_ fold change and FDR values, respectively, while the error bars show the ranges for clone 1 and 2. * marked genes represented known CgpS targets (4). (C) RNA-seq read coverage profiles plotted over the CGP3 prophage genes cg1974-77 (NCgl1682-85) of clones expressing sgRNA-CS1 and sgRNA-CS3, see supplementary Figure S5 and Table S5. (D) CGP3 induction leads to excision and circularization of the prophage DNA element, which can be detected by qPCR with primers B040 (P1) and B041 (P2). Dot plot represents the 2^-ΔΔCT^ values based on geometric C_T_ means and error bars the range of relative circular CGP3 levels measured in biological triplicates and technical duplicates by qPCR. Plasmid IDs (#number) are summarized in Table S2.

We found the differentially expressed genes contained only two known CgpS targets (4): cg1974 (NCgl1682) – the targeted gene - and cg1976 (NCgl1684) (highlighted by * in Figure 7B). Together with cg1975 (NCgl1683), these three neighboring loci were by far the most upregulated genes in cells expressing sgRNA-CS3 (Figure 7A, B Figure S5). The highest upregulation of more than 44-fold (log_2_ fold change 5.5) was observed for the targeted prophage gene cg1974 (NCgl1682). The divergently oriented gene cg1975 (NCgl1683) increased expression by more than 20-fold and represented the second-most upregulated gene (FDR-*p*-values <0.05) (Figure 7A-C, Figure S5A). This increased P_cg1975_-derived transcription was likely a direct effect resulting from dCas9-mediated interference with the CgpS nucleoprotein complex in the divergently oriented P_cg1974_/P_cg1975_ promoter region.

Previous analyses from Pfeifer and colleagues revealed that overexpression of a truncated CgpS variant has a dominant-negative effect on the silencing mechanism, and results in 194 significantly differentially expressed genes (58). Expression of a truncated CgpS variant also led to prophage excision and circularization of the CGP3 element (4). In contrast to this relative broad response, our CRISPR/dCas9-based counter-silencing approach seemed to act more specifically, resulting in only 15 significantly affected genes (Figure 7A, Figure S5A) and no increased copy number of the circular CGP3 (Figure 7D). In conclusion, our results demonstrated that CgpS target genes located within the CGP3 prophage element could be specifically activated by counter-silencing.

### Increased neighboring transcriptional activities can counteract CgpS-mediated xenogeneic silencing

The third most prominent significantly upregulated candidate in the transcriptome analysis was the known CgpS target cg1976 which is located downstream of cg1975 (Figure 7A-C, Figure S5A). Recently, it was demonstrated that increased transcription rates in the neighborhood of silenced genes might counteract the xenogeneic silencing mechanism (59–61). Based on these findings, we hypothesized that an increased transcription rate of gene cg1975 favored the expression of the downstream located cg1976 gene. To investigate the effect of high transcription rates upstream of a CgpS silenced promoter, we inserted short constitutive promoters upstream of the P_cg1974_-*venus* reporter sequence in tandem or divergent orientation (Figure 8). Interestingly, all three tandemly oriented promoters efficiently counteracted CgpS silencing at P_cg1974_. According to their promoter strength in *E. coli* (62), the highest Venus reporter outputs were obtained with the tandemly oriented P_J23119_ promoter, followed by P_J23150_ (both from the Anderson Promoter Collection (https://parts.igem.org/Promoters/Catalog/Constitutive)) and P*_Lac_* (62), demonstrating transcription rate-dependent effects on the silenced prophage promoter. Surprisingly, even divergently oriented promoters affected the P_cg1974_ promoter activity, although the effects were reduced compared to the tandem constructs (Figure 8). The observed bidirectional promoter activity of P_J23119_ highlighted that divergent promoter activities should be considered when working with these promoter systems, including putative effects on DNA regions targeted by xenogeneic silencers.

**Figure 8.**
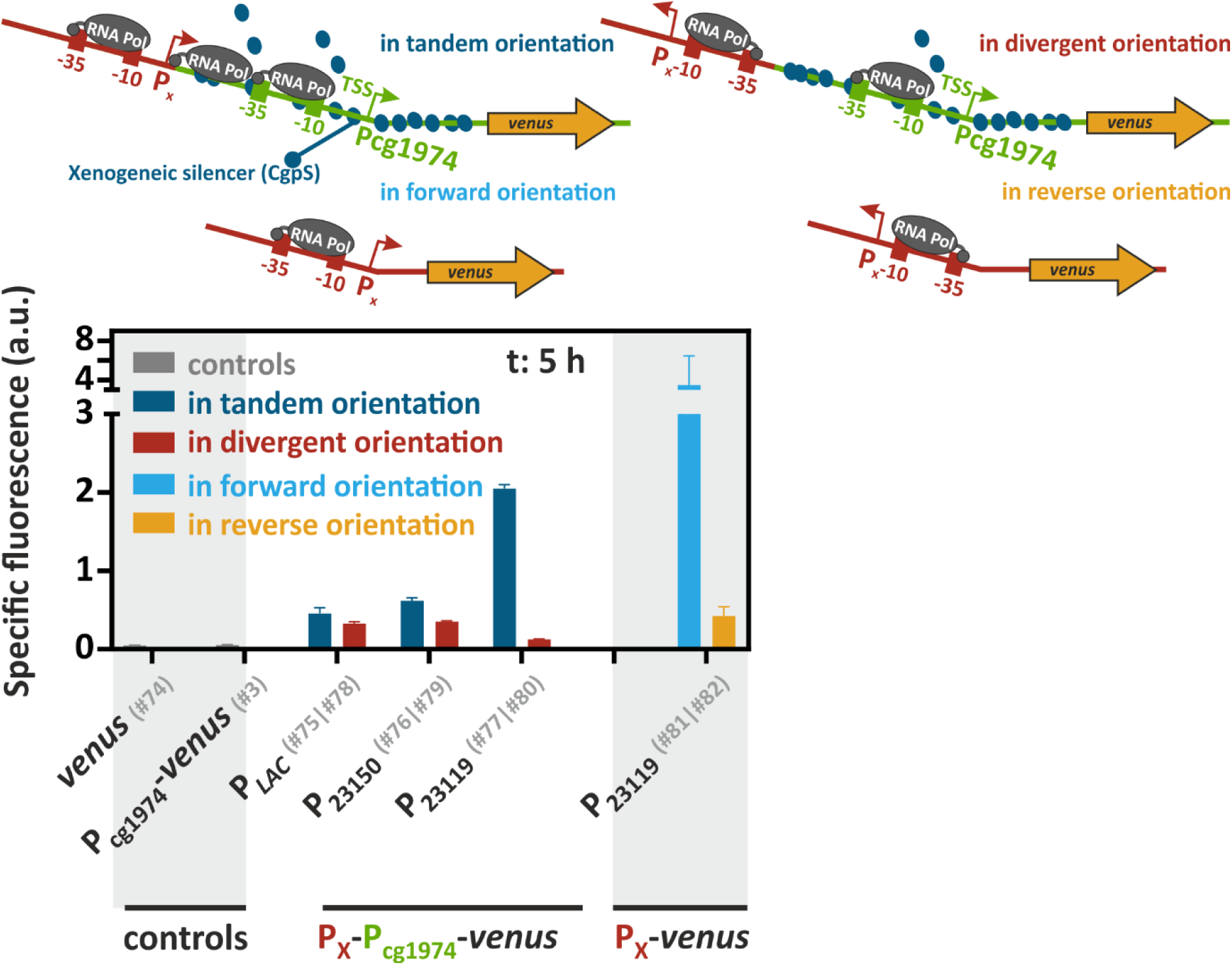
Enhanced transcriptional activity upstream of the CgpS target promoter P_cg1974_ counteracted xenogeneic silencing. Schematic overview represents the *venus* reporter constructs with upstream inserted arrangements of constitutive promoters P*_LAC_*, P_J23150_ or P_J23119_ (P_X_, red) in tandem and in divergent orientation with the CgpS target promoter P_cg1974_ (green). Bar plots show backscatter-normalized reporter outputs (t: 5 h; specific Venus fluorescence) of prophage-free, *cgpS* expressing *C. glutamicum* strains (Δphage::P*_cgpS_*-*cgpS*, (33)) (n=3). Cells designated as controls harboured only the *venus* (#74) or the P_cg1974_-*venus* sequence (#3). Tandem promoter orientations were investigated with plasmids #75 (P*_LAC_*), #76 (P_J23150_) and #77 (P_J23119_), while divergent promoter orientations were analyzed with plasmids #78 (P*_LAC_*), #79 (P_J23150_) and #80 (P_J23119_). Direct fusions of P_J23119_ with the *venus* gene independent of P_cg1974_ were constructed in forward (#81) or reverse promoter orientation (#82). Plasmid IDs (#number) are summarized in Table S2.

In conclusion, we showed that high transcription rates upstream of a prophage promoter could lead to counter-silencing of CgpS. These findings are in agreement with the previous observations for H-NS and Lsr2 (59–61). Apparently, interference with the formation of the XS nucleoprotein complex can be achieved either via direct competition of DNA-binding proteins at the XS nucleation site or by actively transcribing RNA polymerase initiated from upstream promoters.

### Application of CRISPRcosi to increase antibiotic production by *Streptomyces*

Having demonstrated CRISPRcosi effects on CgpS silencing in *C. glutamicum*, we sought to test whether these observations could be extended to other bacteria, for example to the pharmacologically interesting streptomycetes. In *S. venezuelae*, Lsr2 (CgpS-equivalent) plays an important role in repressing a wide array of specialized metabolic clusters that direct natural product synthesis (31), including the chloramphenicol biosynthetic cluster (61). We compared the effects of guiding dCas9 to a key promoter region (CRISPRi), with the effects of targeting the Lsr2 nucleation site within the downstream coding sequence (CRISPRcosi) (Figure 9A). As expected, chloramphenicol production levels dropped when the promoter region was targeted by dCas9 (Figure 9B, C), as had been expected from the CRISPRi observations in *C. glutamicum*. In contrast, targeting of the Lsr2 nucleation site within the downstream coding sequence led to a significant increase (approximately three times) in chloramphenicol production, relative to what had been observed for the no sgRNA wild-type control (Figure 9B, C). It has to be noted that also addition of thiostrepton alone induced chloramphenicol production of a *S. venezuelae* wild-type strain carrying a *dcas9*-free plasmid (that integrated into the same chromosomal locus) in a dose-dependent manner (Figure S6D). However, under the applied conditions (50 µg/ml thiostrepton) we did observe higher chloramphenicol production in the presence of the guide RNA targeting the Lsr2 nucleation site.

**Figure 9.**
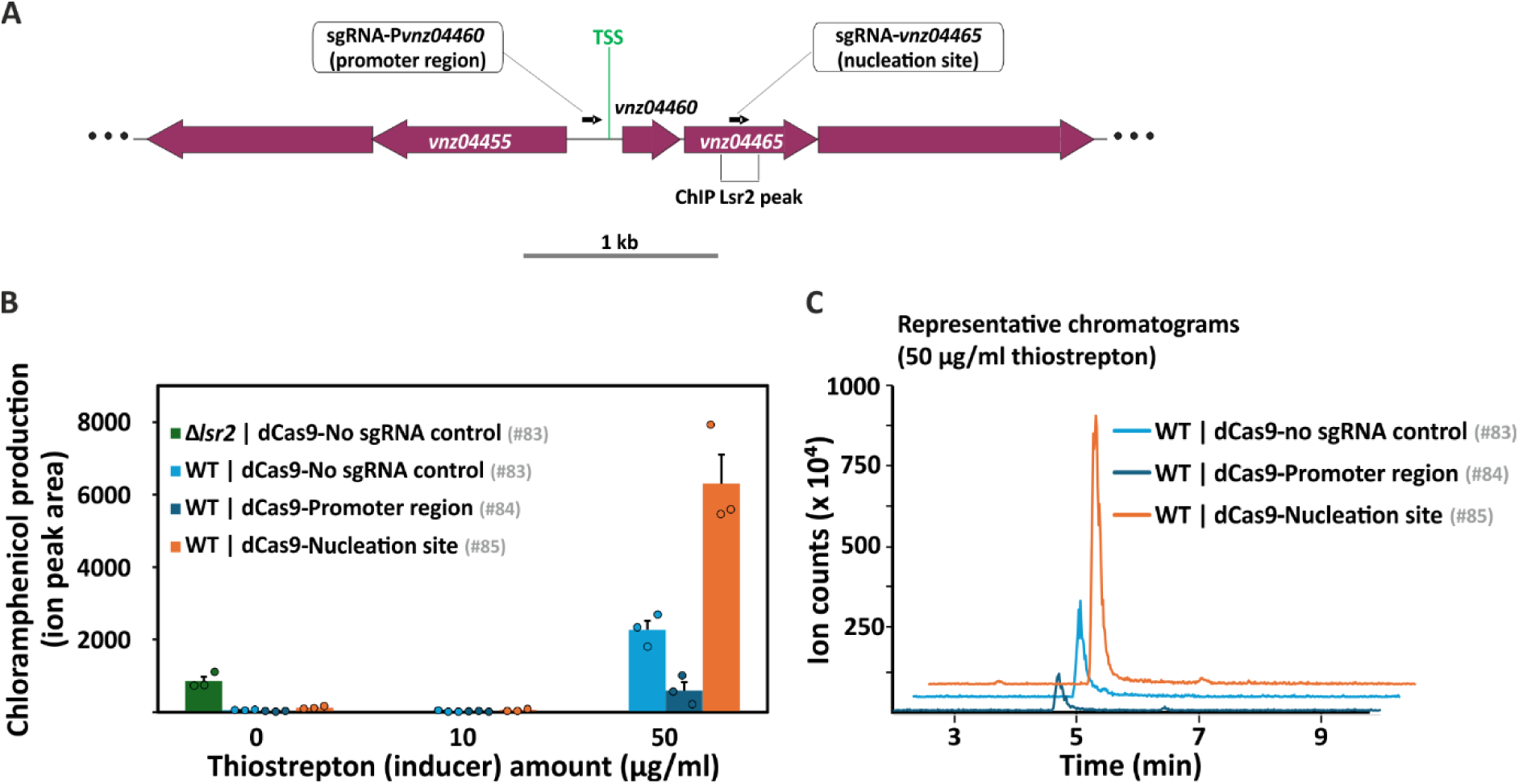
Impact of CRISPR counter silencing and interference in antibiotic (chloramphenicol) production by *Streptomyces venezuelae*. (A) Schematic of the chloramphenicol biosynthetic genes targeted by Lsr2 (CgpS equivalent in the streptomycetes), including the promoter region and transcription start site (TSS) of *vnz04460*, the Lsr2 nucleation site within *vnz04465* (site of maximal peak in Lsr2 chromatin-immunoprecipitation experiments), and the relative location of forward-oriented sgRNAs. CRISPR counter silencing: sgRNA-*vnz04465* designed to bind the template strand within an Lsr2 nucleation site. CRISPR interference: sgRNA-*vnz04460* designed to bind the promoter region. (B) Bar plots represent chloramphenicol production by *S. venezuelae* wild-type carrying empty dCas9 plasmid control (no sgRNA in light blue, #83), sgRNA-*vnz04460* (promoter-targeting in dark blue, #84), sgRNA-*vnz04465* (nucleation-targeting in orange, #85), or a *lsr2* mutant carrying the empty plasmid as a positive uninduced control (green, #83). Expression of *dcas9* was induced using thiostrepton (10 and 50 µg/mL). Error bars indicate standard error of three biological replicates and colored circles the values of each replicate of MYM liquid cultures grown for 3 days. Ion chromatograms from LC-MS measurements are given in (C) (50 µg/ml thiostrepton) as well as Figure S6B (0 µg/ml thiostrepton) and C (10 µg/ml thiostrepton). (C) Representative ion chromatograms of chloramphenicol measurements ([M-H]^-^ at *m/z* 321.00) from the extracts of the dCas9 constructs-carrying strains under induction by 50 µg/mL thiostrepton. Chloramphenicol was eluted at 4.75 min under the gradient used in the mobile phase, shown by major peaks close to the 5 min mark in the figure. Plasmid IDs (#number) refer to Table S2.

In an effort to demonstrate that the disruption of Lsr2 repression by CRISPRcosi was responsible for the increased chloramphenicol production, we attempted the same experiment in an *lsr2* mutant background, using both promoter- and Lsr2 nucleation site-targeting sgRNAs. The growth of these strains was, however, profoundly compromised following thiostrepton addition, and consequently, it was not possible to monitor chloramphenicol production or compare with wild-type.

Altogether, these findings suggested dCas9-mediated Lsr2 counter-silencing within the chloramphenicol biosynthesis gene cluster of *S. venezuelae*, thereby demonstrating the broad applicability of this CRISPRcosi approach across Actinobacteria.

## DISCUSSION

While xenogeneic silencers are important and often even essential for their host’s fitness (4, 19, 20, 63, 64), their high silencing efficiency often limit the activation of target genes and gene clusters. Previous genome-wide counter-silencing approaches have employed truncated or competing XS variants to broadly disrupt the silencing mechanism (4, 65). In contrast, native and synthetic TF-based counter-silencing systems enable the targeted reactivation of silenced genes (33–36). However, TF binding is contingent upon the presence of specific operator sequences, often necessitating modifications to the promoters of interest.

In this work, we present CRISPRcosi, a CRISPR/dCas-mediated counter-silencing approach that specifically activates prophage promoters silenced by the Lsr2-like XS protein CgpS from *C. glutamicum,* and that relieves Lsr2 repression of the chloramphenicol biosynthetic cluster in *S. venezuelae*. We systematically probed the potential for this approach using CgpS as a model. Our results provide a promising system for the precise and tunable upregulation of silenced genes that is independent of promoter engineering. We demonstrated that guide RNAs can direct both dCas9 and dCas12a to silenced promoter regions, leading to interference with the CgpS nucleoprotein complex and counter-silencing. Here, the most effective binding positions were located close to the XS nucleation site. We note that 80% of the five effective counter-silencing guide RNAs bound outside of the core promoter region. This trend was in line with previous CRISPRi studies demonstrating that dCas9 binding at the core promoter leads to repression by inhibiting RNA polymerase binding (38, 41). Our CRISPRcosi experiments did not conclusively determine a preferred DNA strand for counter-silencing, but did reveal that directing different dCas enzymes to nearly identical positions on the coding strand of the P_cg1974_ core promoter yielded contrasting outcomes. Guide RNAs sgRNA-CS1 and crRNA-7 shared 17 of 20 nucleotides; however, only dCas12a facilitated counter-silencing, while dCas9 led to CRISPRi-mediated repression. This aligns with prior studies indicating that dCas9 targeting the coding strand effectively blocks transcription elongation (38, 41), while dCas12a-mediated CRISPRi is more efficient on the template strand (39). These divergent preferences are attributed to the opposite orientations of the 5’-dCas12a hairpin and the 3’-dCas9 hairpin in the guide RNAs (38, 39).

Our systematic screen, which included various CgpS target promoters, guide RNAs and two dCas enzymes, revealed that some XS target promoter regions were not susceptible to CRISPRcosi. We hypothesize that multiple factors hampered activation of the CgpS target promoters, including counteracting CRISPRi effects, a high level of cooperativity of the silencer-DNA complex, and limited availability of suitable PAM sequences as well as interference with further transcription factors or nucleoid-associated proteins. Here, we showed that using different dCas enzymes with distinct PAM sequences could expand the applicability of our approach. In contrast to TF-based counter-silencing approaches that suggested a certain degree of flexibility in promoter architecture (33, 34), the systematic screen of different dCas9 target positions at P_cg1974_ demonstrated that the flexibility of the CRISPRcosi system is limited. Here, a shift of only 12 nucleotides between different sgRNAs (no. 33 and 34) completely abolished the counter-silencing effect. The complex interplay between the XS mechanism and the CRISPRi effects precludes the use of guide RNA arrays and will likely require the screening of individual PAM sites within regions of interest in future applications.

As an alternative counter-silencing approach, we and others have also highlighted the strong impact of upstream promoter activities on XS target promoters. It was demonstrated that increased transcription rates in the neighborhood of silenced genes counteract the xenogeneic silencing mechanism either by inducing positive supercoiling leading to complex destabilization or by remodeling the nucleoprotein complex by the invading transcribing RNA-polymerase (59–61). This principle has been previously demonstrated for the regulation of the chloramphenicol cluster in *S. venezuelae*. Here, the cluster-specific transcription factor CmlR acts as a counter-silencer of Lsr2 by recruiting RNA polymerase. After transcription initiation, RNA polymerase effectively removes bound Lsr2 from the target DNA, highlighting the physiological relevance (61). The results of our study suggest that CRISPRcosi can achieve similar effects. As Lsr2-like proteins are among the most abundant regulators encoded by phages infecting actinobacteria (66), promoter clearance facilitated by specific transcription factors likely represents an important mechanism with additional relevance to the coordination of (pro-)phage gene expression.

The targets of Lsr2-like proteins in actinobacteria are manifold and are important for fundamental research (e.g., illuminating the dark matter of phage genomes), as well as for medical and biotechnological applications. This study establishes CRISPRcosi as a novel approach enabling the targeted activation of genes that are silenced by XS proteins. Combining guide RNA libraries with dCas enzymes may offer an efficient strategy to activate silenced genes. Furthermore, CRISPRcosi has the potential to enhance our understanding of CRISPRi screenings, as identifying upregulated genes could provide insights into chromosome structure mediated by NAPs.

## MATERIAL AND METHODS

### Microbial strains and cultivation conditions

All bacterial strains used in this study are listed in supplementary information Table S1. *Escherichia coli* DH5α cells were used for plasmid amplification and storage. *C. glutamicum* ATCC 13032 (67) and *S. venezuelae* NRRL B-65442 (68) as well as their derivates served as platform strains for the characterization of the CRISPRcosi systems. Detailed information about cultivation conditions is given in Text S1.

### Recombinant DNA work

All used plasmids are listed in Table S2. Each plasmid was assigned a unique identifier (#letter/number) that remains consistent throughout the study to facilitate referencing. Standard molecular methods were performed according to the manufactureŕs instructions or as previously described (69, 70) (71). Plasmid sequencing and synthesis of DNA oligonucleotides were performed by Eurofins Genomics (Ebersberg, Germany), Plasmidsaurus using Oxford Nanopore Technology, or the Farncombe Sequencing Facility (McMaster University) and Integrated DNA Technologies (Table S3). Plasmid transfer into *C. glutamicum* was carried out by electroporation (72). Detailed information about the design of all plasmids constructed in this study is available in the supplementary material (Table S2). The stepwise constructions of *S. venezuelae* plasmids is described in Text S1 in the supplementary material (“Recombinant DNA work”). Briefly, all of them contained a thiostrepton-inducible promoter in front of the *dcas9* gene, the *tipA* gene (whose product is required for thiostrepton induction), a translationally-acting theophylline riboswitch (73) upstream of *tipA,* the phiC31 *attP-int* locus (74), and, in case of pMC346 (#84) and pMC347 (#85), a specific spacer-encoding sequence upstream of the guide RNA scaffold. Sequence-validated plasmids were passaged through the non-methylating *E. coli* strain ET12567 (75) and introduced into *S. venezuelae* via conjugation (76).

### Design and reporter-based characterization of the CRISPRcosi system

The CRISPRcosi system was systematically characterized by performing reporter-based promoter activity studies with *C. glutamicum* Δphage::P*_cgpS_*-*cgpS* (33) cells producing the XS protein CgpS or with *C. glutamicum* Δphage (77) control cells lacking CgpS (Table S1). The key components of this approach were a prophage promoter that is silenced by CgpS, a dCas protein, and a specific guide RNA that targeted the dCas protein to a specific DNA position within the silenced promoter region. Therefore, cells harbored different pJC1 (pCG1 origin of replication)- and/or pEC-XC99E-derivates (pGA1 origin of replication) containing the respective DNA sequences (Table S2).

dCas encoding genes under control of IPTG inducible (*lacI*-P*_tac_*-*dcas9*) or constitutive expression systems (P*_lacM_*-*dcas12a*) were inserted into the respective plasmids. To express guide RNAs (sgRNA, crRNA), the constitutive promoters P*_tacΔ_* (78) or P_J23119_ (from Anderson Promoter Collection https://parts.igem.org/Promoters/Catalog/Constitutive) were fused to the corresponding guide RNA-encoding sequences and inserted into either the pEC-XC99E- or pJC1-based plasmid. A detailed description of the guide RNA design, which followed recently published guidelines (45, 79), is provided in Text S1 in the supplementary material (“Guide RNA design”). All sgRNA and crRNA spacer sequences are listed in Table S4. To monitor promoter activities via a reporter system (yellow fluorescence), prophage promoters known to be silenced by CgpS were fused to the reporter gene *venus* via a linker containing a stop codon and an optimal *C. glutamicum* ribosomal binding site (80).

We systematically analyzed the CRISPRcosi system by performing fluorescence microscopy or by monitoring growth and Venus fluorescence of respective *C. glutamicum* strains during cultivation in the BioLector I® microtiter system (Beckman Coulter, Brea, US-CA) (81). A detailed description of both reporter-based methods is provided in Text S1 in sections “Fluorescence microscopy” and “Microtiter cultivation to monitor growth and fluorescence in reporter assays”. Briefly, cells were cultivated in microtiter plates in CGXII minimal medium (82) supplemented with 111 mM glucose, respective antibiotics (25 µg/ml kanamycin and/or 10 µg/ml chloramphenicol) and up to 200 µM IPTG for P*_tac_*-derived *dcas9* expression as required.

### Transcriptome analysis

In addition to the plasmid-based approaches to monitor promoter activities with reporter genes, we set out to evaluate the effectiveness and specificity of CRISPRcosi on silenced genes in their native context. Therefore, RNA-seq analysis was performed with *C. glutamicum* wild-type cells co-expressing *dcas9* and sgRNA-CS3 (#2) or sgRNA-CS1 (#5). A detailed description is given in Text S1 of the supplemental material in section “Transcriptome analysis”. Briefly, pre-treatment and sequencing of total RNA from mid-exponential phase cells (7 h) was performed by Genewiz (Leipzig, Germany) using the Illumina NovaSeq system. The *C. glutamicum* ATCC 13032 genome (NCBI reference sequence NC_003450.3) combined with the respective sequences of plasmids pEC-XC99E-*lacI*-P*_tac_*-*dcas9*--P*_tacΔ_*-sgRNA-CS3 (#2) or pEC-XC99E-*lacI*-P*_tac_*-*dcas9*--P*_tacΔ_*-sgRNA-CS1 (#5) served as reference. Please note that ‘cǵ and ‘NCgĺ loci numbers both refer to the strain *C. glutamicum* ATCC 13032 (cg: prophage-free strain, NCBI reference sequence NC_022040.1; NCgl: wild-type, NCBI reference sequence NC_003450.3)). The software CLC genomics workbench v.24 (Qiagen, Hilden, Germany) was used for data analysis and for the identification of significantly differentially expressed genes (FDR-*p*-values<0.05 and |log_2_ fold change values|>1) in clones expressing P_cg1974__sgRNA-CS3 (#2) compared to the P_cg1974__sgRNA-CS1 reference cells (#5) (full data set can be found in Table S5).

To identify putative off-targets of sgRNA-CS1 and sgRNA-CS3, the Cas-OFFinder online tool (http://www.rgenome.net/cas-offinder/) from Bae and colleagues (57) was used (see Text S1 for further information). A full overview of all genome-wide off-targets is given in Table S6.

### Quantitative PCR (qPCR)

To identify counter-silencing-associated effects on prophage induction, quantitative PCR (qPCR) was performed to measure the relative amount of circular CGP3 prophage DNA in cells co-expressing *dcas9* and sgRNA-CS3 (#2) or –CS1 (#5). A detailed summary of the qPCR analysis can be found in the supplemental material (Text S1, “Quantitative PCR (qPCR)”). Briefly, oligonucleotides B040/B041 were used to detect the circularized CGP3 prophage DNA element, while oligonucleotides B042/B043 amplified a fragment from the *ddh* (NCgl2528) reference gene (Table S3C). For data evaluation, the C_T_ value-based 2^−ΔΔCT^ method from Livak and Schmittgen (83) was applied. Here, 2^-ΔΔCT^ values and the ranges of circular CGP3 levels were calculated based on the geometric C_T_ means of biological triplicates and technical duplicates.

### Analysis of Streptomyces metabolite extracts

To assess the potential of CRISPRcosi in organisms beyond *C. glutamicum,* we tested our system in *S. venezuelae*, focusing on the ability to activate the chloramphenicol biosynthesis gene cluster, which is normally repressed by the xenogeneic silencer Lsr2 (61). Detailed information about the cultivation conditions, metabolite extraction and analytical steps to quantify the chloramphenicol levels are available in section “Analysis of *Streptomyces* metabolite extracts” of Text S1 in the supplemental material.

### Statistical Analyses

The software GraphPad prism 8.00 (GraphPad Software, La Jolla, US-CA) was used to calculate areas under curves, and significance levels (two-tailed, unpaired t tests) of backscatter-normalized reporter outputs (specific fluorescence) were based on biological triplicates (n=3). The software CLC genomics workbench v.24 (Qiagen, Hilden, Germany) was used for statistical analysis of the RNA-seq data (see *Transcriptome analysis* section). Statistical analysis of the qPCR data is described in the *Quantitative PCR (qPCR)* section in Text S1.

### Data availability

The RNA-seq data were deposited in the European Nucleotide Archive (ENA) at EMBL-EBI (https://www.ebi.ac.uk/ena/browser/home) under accession number PRJEB78733. Plasmids constructed in this work will be shared on request to the corresponding author.

## ACKNOWLEDGEMENTS

We give thanks to our talented students Anna Mosmann, Jan Schorling, Sophia Lorke and Benjamin Chapple for their help with plasmid constructions. We thank Lukas Osswald for the plasmid pJYS3-*dcas12a* (#AB), Dr. Tilmann Weber for plasmid pCRISPR-dCas9 (#AT), and Dr. Susana Matamouros for plasmids pSM22 (#A) and pSM24 (#B).

