## Supplementary Material, Figure S1-6, Tables S1-S4 for "CRISPR/dCas-mediated counter-silencing – Reprogramming dCas proteins into antagonists of xenogeneic silencers"

### TEXT SUPPLEMENTARY MATERIAL

Text S1. Supplementary information on methods used in this study. (Separate file)

### TABLES SUPPLEMENTARY MATERIAL

Table S1. Strains used in this study.

Table S2. Plasmids used in this study.

Table S3A. Oligonucleotides used for plasmid constructions (see Table S2).

Table S3B. Oligonucleotides used for sequencing and colony PCR (see Table S2).

Table S3C. Oligonucleotides used for qPCR.

Table S4. Overview of guide RNA spacer sequences.

Table S5. RNA-seq - Differential gene expression (cg1974_sgRNA-CS3 clone 1 (A) and 2 (B)). (Excel file)

Table S6. Off-targets of cg1974_sgRNA-CS1 (A) and –CS3 (B). (Excel file)

### FIGURES SUPPLEMENTARY MATERIAL

Figure S1. Correlation of CRISPRcosi efficiencies at prophage promoter $P_{cg1974}$ (A) and $P_{cg2014}$ (B) and *dcas9* expression levels.

Figure S2. The crRNA binding positions determine the dCas12a-mediated CRISPRcosi efficiency.

Figure S3. Supplementary information on Figure 5: dCas9 and dCas12a both mediate efficient CRISPRcosi.

Figure S4. Supplementary information on Figure 6: CRISPRcosi counteracts CgpS-mediated silencing at different prophage promoters.

Figure S5. Supplementary data on Figure 7: Investigation of potential off-target effects of CRISPRcosi.

Figure S6. Supplementary data on Figure 9: Impact of CRISPR counter silencing and interference in antibiotic (chloramphenicol) production by *Streptomyces venezuelae*.

**Table S1. Strains used in this study.**

The published strain names are given in brackets after the reference in case of deviations.

| Strain | Relevant characteristics | Reference |
|---|---|---|
| *Escherichia coli* DH5α | $F^-$ Φ80*lacZ*ΔM15 Δ(*lacZYA-argF*) U169 *recA1 endA1 hsdR17*($r_k^-$, $m_k^+$) *phoA supE44 thi-1 gyrA96 relA1* $\lambda^-$, strain used for cloning procedures | Invitrogen |
| *E. coli* ET12567 (pUZ8002) | $F^-$ *dam13*::Tn9 *dcm6 hsdM hsdR recF143 zjj201*::Tn10 *galK2 galT22 ara14 lacY1 xyl5 leuB6 thi1 tonA31 rpsL136 hisG4 tsx78 mtli glnV44*, pUZ8002 (helper plasmid)<br><br>non-methylating *E. coli* strain used for conjugation of *Streptomyces* | (1) |
| *Corynebacterium glutamicum* ATCC 13032 | Biotin-auxotrophic *C. glutamicum* wild-type | (2) |
| *C. glutamicum* ATCC 13032::$P_{cg1974}$-*eyfp* | Derivative of ATCC 13032 containing the prophage reporter $P_{cg1974}$-*eyfp* integrated into the intergenic region of cg1121-cg1122 | (3) (ATCC 13032::$P_{lys}$-*eyfp*) |
| Δphage | Derivative of ATCC 13032 with deletion of prophages CGP1 (cg1507-cg1524), CGP2 (cg1746-cg1752), and CGP3 (cg1890-cg2071) | (4) (MB001) |
| Δphage::$P_{cgpS}$-*cgpS* | Derivative of Δphage (MB001) with re-integrated *cgpS* together with its native promoter $P_{cgpS}$ in the intergenic region of cg1199-cg1201 | (5) |
| *Streptomyces venezuelae* NRRL B-65442 | *S. venezuelae* wild-type | (6) |
| E327A | *S. venezuelae* Δ*lsr2* | (7) |

| | | |
|---|---|---|
| E339 | *S. venezuelae* wild-type carrying the plasmid pIJ6902, in which the apramycin resistance gene has been replaced with a hygromycin resistance gene | This work |

**Table S2. Plasmids used in this study.**

All constructed plasmids were based on the pJC1, pSM22 or the pEC-XC99E vector systems providing a kanamycin (*kan*$^R$), carbenicillin (*amp*$^R$) or chloramphenicol resistance (*cm*$^R$), respectively. pJC1- (pCG1 replication origin) and pEC-XC99E- (pGA1 replication origin) based plasmids were transferred into *C. glutamicum* cells. If needed, pJC1- and pEC-XC99E-derivates were co-transferred as a double-plasmid system. pSM22-based plasmids were designed as intermediate constructs to facilitate cloning of pJC1 and pEC-XC99E derivatives. All plasmids used for *S. venezuelae* were based on the pCRISPR-dCas9 plasmid which was a gift from Tilmann Weber (Addgene plasmid #125687; http://n2t.net/addgene:125687 RRID:Addgene_125687) (8). We assigned an ID (indicated by #) to all plasmids and categorized them into two different groups. Group 1, labelled with letters, contains plasmids used from other sources (#a-#d) and from this work (#A-#AV) that were not used in experiments but were used as helper vectors for plasmid construction. Group 2 contains plasmids from this work and other sources (in case of differences, the published plasmid names are given in brackets after the reference) that were used to characterize the principle of CRISPRcosi in experiments. These plasmids were numbered (#1-#85) according to their first mention in the results section. Oligonucleotide pairs used in PCRs for insert amplification (Table S3A) are given as numbers in the column "Construction of plasmids used in this work". DNA templates are listed in brackets behind the oligonucleotides. Chromosomal *C. glutamicum* ATCC 13032 DNA was isolated as noted previously (9) and served as PCR template for insert amplification if indicated. Plasmid backbones including the restriction enzymes used for linearization are given (*). Oligonucleotides used for colony PCR (C) and Sequencing (S) are listed (Table S3B). Information about promoter regions bound by CgpS were based on previous work from Pfeifer et al. (10).

| Plasmid ID | Plasmid | Construction of plasmids used in this work | Relevant characteristics | Oligonucleotides used for colony PCR (C) and sequencing (S) | Source or reference |
|---|---|---|---|---|---|
| #a | pEC-XC99E | | *cm*$^R$, *E. coli*/*C. glutamicum* shuttle vector (*cat*$_I$, *lacI*$^q$, P$_{trc}$, *rrnB* (T1 and T2), *oriV*$_{Ec}$, *per*, *repA* (pGA1 *ori*$_{Cg}$) | | (11) |
| #b | pPBEx2 | | *kan*$^R$, *E. coli*/*C. glutamicum* shuttle vector (P$_{tacI}$; *lacI*$^q$; *ori*$_{Cg}$ from pBL1, *ori*$_{Ec}$ ColE1 from pUC18) | | (12) |
| #c | #44251 (addgene) | | *amp*$^R$, *E. coli* vector (*ori*$_{Ec}$ ColE1 from pUC19, contains the P$_{J23119\ (SpeI)}$-guide RNA (bacteria) sequence) | | (13) |
| #d | pCLTON1 | | *kan*$^R$, *E. coli*/*C. glutamicum* shuttle vector (*B. subtilis* | | (14) |

| | | | | | |
|---|---|---|---|---|---|
| | | | derived $P_{gap}$-*tetR*__$P_{tet}$ expression system, $oriV_{Cg}$, $oriV_{Ec}$) | | |
| #A | pSM22 | Three consecutive site-directed mutagenesis (SDM) rounds (1st: OSM35/OSM36; 2nd: OSM37/OSM38; 3rd: OSM47/OSM48 on plasmid #c | *amp*$^{R}$, #44251 (#c) derivative containing the $P_{J23119\ (SpeI)}$-promoter followed by an optimized sgRNA hairpin sequence, based on (13) | | Gift from Susana Matamouros |
| #B | pSM24 | Gibson assembly: OSM13/OSM14 into (#d)*PstI, followed by SDM with OSM39/OSM40 | *kan*$^{R}$, pCLTON1$^{TS}$ (RepA$^{P47S}$) (#d) derivative containing the *dcas9* gene | | Gift from Susana Matamouros |
| #C | pEC-XC99E-*dcas9*-sgRNA_hairpin-$T_{Sp}$ | Gibson assembly: R69/R70 (#A) and R67/R68 (#B) into #a *NdeI *SalI | *cm*$^{R}$, pEC-XC99E (#a) derivative containing a promoter-less *dcas9* gene and, in divergent orientation and separated by a linker sequence containing the XmaI and SpeI sites, the sequence encoding a structurally optimized sgRNA hairpin followed by a *Streptococcus* | C: R71/R72<br>S: R71, R72, R73, R74, R75, R76, R77, R78 | This work |

| | | | | | |
|---|---|---|---|---|---|
| | | | *pyogenes*-derived terminator (13) (the term sgRNA hairpin includes the hairpin and the terminator in the following) | | |
| #D | pEC-XC99E-*lacI*-$P_{tac}$-*dcas9*--$P_{tac\Delta}$-SpeI_site-sgRNA_hairpin-$T_{Sp}$ | Gibson assembly R91/R92 (#b) and R93/R94 (#b) into #C *SpeI *XmaI | *cm*$^R$, pEC-XC99E-*dcas9*-sgRNA_hairpin-$T_{Sp}$ (#C) derivative with an inserted *lacI*$^q$-$P_{tac}$ sequence upstream of *dcas9* and a truncated, constitutive $P_{tac\Delta}$ promoter upstream of the sgRNA hairpin sequence, separated by a SpeI site | C: R95/R96<br>S: R169, R171 | This work |
| #E | pSM22-$P_{J32119}$-$P_{cg1974}$_sgRNA-CS3 | Gibson assembly 864/865 (#A *SpeI), circularized | *amp*$^R$, pSM22 derivative with an inserted $P_{cg1974}$_sgRNA-CS3 spacer encoding sequence at the SpeI site between the $P_{J23119}$ promoter and the sgRNA hairpin | S: 890 | This work |
| #F | pSM22-$P_{J32119}$-$P_{cg1974}$_sgRNA-25 | Gibson assembly 1711/1712 (#A *SpeI), circularized | *amp*$^R$, pSM22 derivative with an inserted $P_{cg1974}$_sgRNA-25 spacer encoding sequence at the SpeI site between the $P_{J23119}$ promoter | S: 890 | This work |

| | | | | | |
|---|---|---|---|---|---|
| | | | and the sgRNA hairpin | | |
| #G | pSM22-$P_{J32119}$-$P_{cg1974}$_sgRNA-26 | Gibson assembly 1713/1714 (#A *SpeI), circularized | *amp*$^R$, pSM22 derivative with an inserted $P_{cg1974}$_sgRNA-26 spacer encoding sequence at the SpeI site between the $P_{J23119}$ promoter and the sgRNA hairpin | S: 890 | This work |
| #H | pSM22-$P_{J32119}$-$P_{cg1974}$_sgRNA-28 | Gibson assembly 1715/1716 (#A *SpeI), circularized | *amp*$^R$, pSM22 derivative with an inserted $P_{cg1974}$_sgRNA-28 spacer encoding sequence at the SpeI site between the $P_{J23119}$ promoter and the sgRNA hairpin | S: 890 | This work |
| #I | pSM22-$P_{J32119}$-$P_{cg1974}$_sgRNA-29 | Gibson assembly 1717/1718 (#A *SpeI), circularized | *amp*$^R$, pSM22 derivative with an inserted $P_{cg1974}$_sgRNA-29 spacer encoding sequence at the SpeI site between the $P_{J23119}$ promoter and the sgRNA hairpin | S: 890 | This work |
| #J | pSM22-$P_{J32119}$-$P_{cg1974}$_sgRNA-CS1 | Gibson assembly 1719/1720 (#A *SpeI), circularized | *amp*$^R$, pSM22 derivative with an inserted $P_{cg1974}$_sgRNA-CS1 spacer encoding sequence at the | S: 890 | This work |

| | | | | | |
|---|---|---|---|---|---|
| | | | SpeI site between the $P_{J23119}$ promoter and the sgRNA hairpin | | |
| #K | pSM22-$P_{J32119}$-$P_{cg1974}$_sgRNA-33 | Gibson assembly 1721/1722 (#A *SpeI), circularized | *amp$^R$*, pSM22 derivative with an inserted $P_{cg1974}$_sgRNA-33 spacer encoding sequence at the SpeI site between the $P_{J23119}$ promoter and the sgRNA hairpin | S: 890 | This work |
| #L | pSM22-$P_{J32119}$-$P_{cg1974}$_sgRNA-34 | Gibson assembly 1723/1724 (#A *SpeI), circularized | *amp$^R$*, pSM22 derivative with an inserted $P_{cg1974}$_sgRNA-34 spacer encoding sequence at the SpeI site between the $P_{J23119}$ promoter and the sgRNA hairpin | S: 890 | This work |
| #M | pSM22-$P_{J32119}$-$P_{cg1974}$_sgRNA-35 | Gibson assembly 1725/1726 (#A *SpeI), circularized | *amp$^R$*, pSM22 derivative with an inserted $P_{cg1974}$_sgRNA-35 spacer encoding sequence at the SpeI site between the $P_{J23119}$ promoter and the sgRNA hairpin | S: 890 | This work |
| #N | pSM22-$P_{J32119}$-$P_{cg1974}$_sgRNA-36 | Gibson assembly 1727/1728 (#A | *amp$^R$*, pSM22 derivative with an inserted $P_{cg1974}$_sgRNA-36 | S: 890 | This work |

| | | | | | |
|---|---|---|---|---|---|
| | | *SpeI), circularized | spacer encoding sequence at the SpeI site between the $P_{J23119}$ promoter and the sgRNA hairpin | | |
| #O | pSM22-$P_{J32119}$-$P_{cg1974}$_sgRNA-37 | Gibson assembly 1729/1730 (#A *SpeI), circularized | *amp*$^R$, pSM22 derivative with an inserted $P_{cg1974}$_sgRNA-37 spacer encoding sequence at the SpeI site between the $P_{J23119}$ promoter and the sgRNA hairpin | S: 890 | This work |
| #P | pSM22-$P_{J32119}$-$P_{cg1974}$_sgRNA-38 | Gibson assembly 1731/1732 (#A *SpeI), circularized | *amp*$^R$, pSM22 derivative with an inserted $P_{cg1974}$_sgRNA-38 spacer encoding sequence at the SpeI site between the $P_{J23119}$ promoter and the sgRNA hairpin | S: 890 | This work |
| #Q | pSM22-$P_{J32119}$-$P_{cg1974}$_sgRNA-39 | Gibson assembly 1733/1734 (#A *SpeI), circularized | *amp*$^R$, pSM22 derivative with an inserted $P_{cg1974}$_sgRNA-39 spacer encoding sequence at the SpeI site between the $P_{J23119}$ promoter and the sgRNA hairpin | S: 890 | This work |

| | | | | | |
|---|---|---|---|---|---|
| #R | pSM22-$P_{J32119}$-$P_{cg1974}$_sgRNA-40 | Gibson assembly 1735/1736 (#A *SpeI), circularized | *amp*$^{R}$, pSM22 derivative with an inserted $P_{cg1974}$_sgRNA-40 spacer encoding sequence at the SpeI site between the $P_{J23119}$ promoter and the sgRNA hairpin | S: 890 | This work |
| #S | pSM22-$P_{J32119}$-$P_{cg2014}$_sgRNA-22 | Gibson assembly 1659/1660 (#A *SpeI), circularized | *amp*$^{R}$, pSM22 derivative with an inserted $P_{cg2014}$_sgRNA-22 spacer encoding sequence at the SpeI site between the $P_{J23119}$ promoter and the sgRNA hairpin | S: 890 | This work |
| #T | pSM22-$P_{J32119}$-$P_{cg2014}$_sgRNA-24 | Gibson assembly 1661/1662 (#A *SpeI), circularized | *amp*$^{R}$, pSM22 derivative with an inserted $P_{cg2014}$_sgRNA-24 spacer encoding sequence at the SpeI site between the $P_{J23119}$ promoter and the sgRNA hairpin | S: 890 | This work |
| #U | pSM22-$P_{J32119}$-$P_{cg2014}$_sgRNA-25 | Gibson assembly 1663/1664 (#A *SpeI), circularized | *amp*$^{R}$, pSM22 derivative with an inserted $P_{cg2014}$_sgRNA-25 spacer encoding sequence at the SpeI site between the $P_{J23119}$ promoter | S: 890 | This work |

| | | | | | |
|---|---|---|---|---|---|
| | | | and the sgRNA hairpin | | |
| #V | pSM22-$P_{J32119}$-$P_{cg2014}$_sgRNA-26 | Gibson assembly 1665/1666 (#A *SpeI), circularized | *amp*$^{R}$, pSM22 derivative with an inserted $P_{cg2014}$_sgRNA-26 spacer encoding sequence at the SpeI site between the $P_{J23119}$ promoter and the sgRNA hairpin | S: 890 | This work |
| #W | pSM22-$P_{J32119}$-$P_{cg2014}$_sgRNA-27 | Gibson assembly 1667/1668 (#A *SpeI), circularized | *amp*$^{R}$, pSM22 derivative with an inserted $P_{cg2014}$_sgRNA-27 spacer encoding sequence at the SpeI site between the $P_{J23119}$ promoter and the sgRNA hairpin | S: 890 | This work |
| #X | pSM22-$P_{J32119}$-$P_{cg2014}$_sgRNA-28 | Gibson assembly 1669/1670 (#A *SpeI), circularized | *amp*$^{R}$, pSM22 derivative with an inserted $P_{cg2014}$_sgRNA-28 spacer encoding sequence at the SpeI site between the $P_{J23119}$ promoter and the sgRNA hairpin | S: 890 | This work |
| #Y | pSM22-$P_{J32119}$-$P_{cg2014}$_sgRNA-29 | Gibson assembly 1671/1672 (#A *SpeI), circularized | *amp*$^{R}$, pSM22 derivative with an inserted $P_{cg2014}$_sgRNA-29 spacer encoding sequence at the | S: 890 | This work |

| | | | | | |
|---|---|---|---|---|---|
| | | | SpeI site between the $P_{J23119}$ promoter and the sgRNA hairpin | | |
| #Z | pSM22-$P_{J32119}$-$P_{cg2014}$_sgRNA-30 | Gibson assembly 1673/1674 (#A *SpeI), circularized | *amp*$^R$, pSM22 derivative with an inserted $P_{cg2014}$_sgRNA-30 spacer encoding sequence at the SpeI site between the $P_{J23119}$ promoter and the sgRNA hairpin | S: 890 | This work |
| #AA | pSM22-$P_{J32119}$-$P_{cg2014}$_sgRNA-31 | Gibson assembly 1675/1676 (#A *SpeI), circularized | *amp*$^R$, pSM22 derivative with an inserted $P_{cg2014}$_sgRNA-31 spacer encoding sequence at the SpeI site between the $P_{J23119}$ promoter and the sgRNA hairpin | S: 890 | This work |
| #AB | pJYS3-*dcas12a* | | *kan*$^R$, pJYS3 derivative with a *dcas12a* (*cpf1* from *Francisella novicida* carrying mutations D917A and E1006A) sequence under control of the constitutive $P_{lacM}$ promoter | | Gift from Lukas Osswald |
| #AC | pSM22-$P_{J23119}$-crRNA_SpeI-site | Gibson assembly 1780/1781 (#A | *amp*$^R$, pSM22 derivative containing the $P_{J23119}$ promoter | C: 890/891<br>S: 890 | This work |

| | | | | | |
|---|---|---|---|---|---|
| | | *SpeI *BamHI), circularized | and the sequence encoding the crRNA hairpin followed by a SpeI site to facilitate spacer insertion (helper plasmid for crRNA amplification) | | |
| #AD | pSM22-$P_{J23119}$-$P_{cg1896}$_sgRNA-1 | Gibson assembly 852/853 (#A *SpeI), circularized | *amp*$^R$, pSM22 derivative with an inserted $P_{cg1896}$_sgRNA-1 spacer encoding sequence at the SpeI site between the $P_{J23119}$ promoter and the sgRNA hairpin | S: 890 | This work |
| #AE | pSM22-$P_{J23119}$-$P_{cg1896}$_sgRNA-2 | Gibson assembly 854/855 (#A *SpeI), circularized | *amp*$^R$, pSM22 derivative with an inserted $P_{cg1896}$_sgRNA-2 spacer encoding sequence at the SpeI site between the $P_{J23119}$ promoter and the sgRNA hairpin | S: 890 | This work |
| #AF | pSM22-$P_{J23119}$-$P_{cg1929}$_sgRNA-1 | Gibson assembly 856/857 (#A *SpeI), circularized | *amp*$^R$, pSM22 derivative with an inserted $P_{cg1929}$_sgRNA-1 spacer encoding sequence at the SpeI site between the $P_{J23119}$ promoter and the sgRNA hairpin | S: 890 | This work |

| | | | | | |
|---|---|---|---|---|---|
| #AG | pSM22-$P_{J23119}$-$P_{cg1940}$_sgRNA-1 | Gibson assembly 858/859 (#A *SpeI), circularized | *amp*$^R$, pSM22 derivative with an inserted $P_{cg1940}$_sgRNA-1 spacer encoding sequence at the SpeI site between the $P_{J23119}$ promoter and the sgRNA hairpin | S: 890 | This work |
| #AH | pSM22-$P_{J23119}$-$P_{cg1940}$_sgRNA-2 | Gibson assembly 860/861 (#A *SpeI), circularized | *amp*$^R$, pSM22 derivative with an inserted $P_{cg1940}$_sgRNA-2 spacer encoding sequence at the SpeI site between the $P_{J23119}$ promoter and the sgRNA hairpin | S: 890 | This work |
| #AJ | pSM22-$P_{J23119}$-$P_{cg1977}$_sgRNA-1 | Gibson assembly 866/867 (#A *SpeI), circularized | *amp*$^R$, pSM22 derivative with an inserted $P_{cg1977}$_sgRNA-1 spacer encoding sequence at the SpeI site between the $P_{J23119}$ promoter and the sgRNA hairpin | S: 890 | This work |
| #AK | pSM22-$P_{J23119}$-$P_{cg1999}$_sgRNA-1 | Gibson assembly 868/869 (#A *SpeI), circularized | *amp*$^R$, pSM22 derivative with an inserted $P_{cg1999}$_sgRNA-1 spacer encoding sequence at the SpeI site between the $P_{J23119}$ promoter | S: 890 | This work |

| | | | | | |
|---|---|---|---|---|---|
| | | | and the sgRNA hairpin | | |
| #AL | pSM22-$P_{J23119}$-$P_{cg1999}$_sgRNA-2 | Gibson assembly 870/871 (#A *SpeI), circularized | *amp^R^*, pSM22 derivative with an inserted $P_{cg1999}$_sgRNA-2 spacer encoding sequence at the SpeI site between the $P_{J23119}$ promoter and the sgRNA hairpin | S: 890 | This work |
| #AM | pSM22-$P_{J23119}$-$P_{cg2004}$_sgRNA-1 | Gibson assembly 872/873 (#A *SpeI), circularized | *amp^R^*, pSM22 derivative with an inserted $P_{cg2004}$_sgRNA-1 spacer encoding sequence at the SpeI site between the $P_{J23119}$ promoter and the sgRNA hairpin | S: 890 | This work |
| #AN | pSM22-$P_{J23119}$-$P_{cg2004}$_sgRNA-2 | Gibson assembly 874/875 (#A *SpeI), circularized | *amp^R^*, pSM22 derivative with an inserted $P_{cg2004}$_sgRNA-2 spacer encoding sequence at the SpeI site between the $P_{J23119}$ promoter and the sgRNA hairpin | S: 890 | This work |
| #AO | pSM22-$P_{J23119}$-$P_{cg2011}$_sgRNA-1 | Gibson assembly 876/877 (#A *SpeI), circularized | *amp^R^*, pSM22 derivative with an inserted $P_{cg2011}$_sgRNA-1 spacer encoding sequence at the | S: 890 | This work |

| | | | | | |
|---|---|---|---|---|---|
| | | | SpeI site between the $P_{J23119}$ promoter and the sgRNA hairpin | | |
| #AP | pSM22-$P_{J23119}$-$P_{cg2014}$_sgRNA-1 | Gibson assembly 878/879 (#A *SpeI), circularized | *amp*$^R$, pSM22 derivative with an inserted $P_{cg2014}$_sgRNA-1 spacer encoding sequence at the SpeI site between the $P_{J23119}$ promoter and the sgRNA hairpin | S: 890 | This work |
| #AQ | pSM22-$P_{J23119}$-$P_{cg2032}$_sgRNA-1 | Gibson assembly 880/881 (#A *SpeI), circularized | *amp*$^R$, pSM22 derivative with an inserted $P_{cg2032}$_sgRNA-1 spacer encoding sequence at the SpeI site between the $P_{J23119}$ promoter and the sgRNA hairpin | S: 890 | This work |
| #AR | pSM22-$P_{J23119}$-$P_{cg2032}$_sgRNA-2 | Gibson assembly 882/883 (#A *SpeI), circularized | *amp*$^R$, pSM22 derivative with an inserted $P_{cg2032}$_sgRNA-2 spacer encoding sequence at the SpeI site between the $P_{J23119}$ promoter and the sgRNA hairpin | S: 890 | This work |
| #AS | pSM22-$P_{J23119}$-$P_{cg2065}$_sgRNA-1 | Gibson assembly 884/885 (#A | *amp*$^R$, pSM22 derivative with an inserted $P_{cg2065}$_sgRNA-1 | S: 890 | This work |

| | | | | | |
|---|---|---|---|---|---|
| | | *SpeI), circularized | spacer encoding sequence at the SpeI site between the $P_{J23119}$ promoter and the sgRNA hairpin | | |
| #AT | #125687 (Addgene)<br>pCRISPR-dCas9 | | *apr*$^R$, *E. coli* vector containing *Streptomyces* codon optimized spdCas9, sgRNA cassette | | (15) |
| #AU | pMC386 | | *thio*$^R$, *E. coli* vector containing theophylline riboswitch controlled *tipA* | | Gift from Meghan Pepler |
| #AV | pSET152 | | *apr*$^R$, *E. coli*-*Streptomyces* shuttle vector containing a phiC31 *attP-int* cassette | | (16) |
| #1 | pEC-XC99E-*lacI*-$P_{tac}$-*dcas9*_no-sgRNA | Gibson assembly: R87/R88 (#b) into #C *SpeI *XmaI | *cm*$^R$, pEC-XC99E-*dcas9*-sgRNA_hairpin-$T_{Sp}$ (#C) derivative with an inserted *lacI*-$P_{tac}$ sequence upstream of the *dcas9* gene | C: R95/R96<br>S: R171, R169 | This work |
| #2 | pEC-XC99E-*lacI*-$P_{tac}$-*dcas9*--$P_{tac\Delta}$-sgRNA-CS3 | Gibson assembly: annealed R176/R177 dsDNA into #D *Bsp1407I *SpeI | *cm*$^R$, pEC-XC99E-*lacI*-$P_{tac}$-*dcas9*--$P_{tac\Delta}$-SpeI_site-sgRNA_hairpin-$T_{Sp}$ (#D) derivative with an inserted $P_{cg1974}$_sgRNA-CS3 spacer encoding | S: R171 | This work |

| | | | | | |
|---|---|---|---|---|---|
| | | | sequence at the SpeI site | | |
| #3 | pJC1-$P_{cg1974}$-*venus* | | *kan*$^R$, pJC1-based plasmid carrying the CgpS bound area of the promoter of cg1974 (444 bp) and the first 30 bp of the coding sequence fused to the reporter gene *venus* via a linker containing a stop codon and an artificial RBS | | (5) (pJC1-$P_{lys}$-*venus*) |
| #4 | pEC-XC99E_empty | Gibson assembly annealed R65/R66 dsDNA into #a *NdeI *SalI | *cm*$^R$, pEC-XC99E (#a) derivative lacking the *lacI*$^q$ gene and the $P_{trc}$ promoter | C: R71/R72<br>S: R71 | This work |
| #5 | pEC-XC99E-*lacI*-$P_{tac}$-*dcas9*--$P_{tac\Delta}$-sgRNA-CS1 | Gibson assembly annealed R172/R173 dsDNA into #D *Bsp1407I *SpeI | *cm*$^R$, pEC-XC99E-*lacI*-$P_{tac}$-*dcas9*--$P_{tac\Delta}$-SpeI_site-sgRNA_hairpin-$T_{Sp}$ (#D) derivative with an inserted $P_{cg1974}$_sgRNA-CS1 spacer encoding sequence at the SpeI site | S: R171 | This work |
| #6 | pJC1-$P_{cg1974}$-*venus*_T_$P_{J23119}$-sgRNA-CS3 | Gibson assembly 1341/1342 (#E) into #3 *SalI | *kan*$^R$, pJC1-$P_{cg1974}$-*venus* (#3) derivative with an inserted $P_{cg1974}$_sgRNA-CS3 sequence under | C: 766/R13<br>S: R13 | This work |

| | | | | | |
|---|---|---|---|---|---|
| | | | control of the $P_{J23119}$ promoter and followed by the *rrnB* terminator behind the *venus*-*rrnB* (T) sequence | | |
| #7 | pJC1-$P_{cg1974\Delta PAM}$-*venus* | Gibson assembly 117/B024 (#3) and B026/116 (#3) into #74 *BamHI *SpeI | *kan*$^R$, pJC1-$P_{cg1974}$-*venus* (#3) derivative with a deleted sgRNA-CS3 specific PAM sequence in $P_{cg1974}$ (change of AGG to AAC 39-37 bp upstream of gene start) | C: R12/R13<br>S: R12, R13 | This work |
| #8 | pJC1-$P_{cg2014}$-*venus* | | *kan*$^R$, pJC1-based plasmid carrying the CgpS bound area of the promoter of cg2014 (545 bp) and the first 30 bp of the coding sequence fused to the reporter gene *venus* via a linker containing a stop codon and an artificial RBS | | (5) |
| #9 | pJC1-$P_{cg1974}$-*venus*_T_$P_{J23119}$-sgRNA-25 | Gibson assembly 1341/1342 (#F) into #3 *SalI | *kan*$^R$, pJC1-$P_{cg1974}$-*venus* (#3) derivative with an inserted $P_{cg1974}$_sgRNA-25 sequence under control of the $P_{J23119}$ promoter and followed by the | C: 554/R13<br>S: 554, R13 | This work |

| | | | | | |
|---|---|---|---|---|---|
| | | | *rrnB* terminator behind the *venus*-*rrnB* (T) sequence | | |
| #10 | pJC1-$P_{cg1974}$-*venus*_T_$P_{J23119}$-sgRNA-26 | Gibson assembly 1341/1342 (#G) into #3 *SalI | *kan$^R$*, pJC1-$P_{cg1974}$-*venus* (#3) derivative with an inserted $P_{cg1974}$_sgRNA-26 sequence under control of the $P_{J23119}$ promoter and followed by the *rrnB* terminator behind the *venus*-*rrnB* (T) sequence | C: 554/R13<br>S: 554, R13 | This work |
| #11 | pJC1-$P_{cg1974}$-*venus*_T_$P_{J23119}$-sgRNA-28 | Gibson assembly 1341/1342 (#H) into #3 *SalI | *kan$^R$*, pJC1-$P_{cg1974}$-*venus* (#3) derivative with an inserted $P_{cg1974}$_sgRNA-28 sequence under control of the $P_{J23119}$ promoter and followed by the *rrnB* terminator behind the *venus*-*rrnB* (T) sequence | C: 554/R13<br>S: 554, R13 | This work |
| #12 | pJC1-$P_{cg1974}$-*venus*_T_$P_{J23119}$-sgRNA-29 | Gibson assembly 1341/1342 (#I) into #3 *SalI | *kan$^R$*, pJC1-$P_{cg1974}$-*venus* (#3) derivative with an inserted $P_{cg1974}$_sgRNA-29 sequence under control of the $P_{J23119}$ promoter and followed by the *rrnB* terminator | C: 554/R13<br>S: 554, R13 | This work |

| | | | | | |
|---|---|---|---|---|---|
| | | | behind the *venus*-*rrnB* (T) sequence | | |
| #13 | pJC1-$P_{cg1974}$-*venus*_T_$P_{J23119}$-sgRNA-CS1 | Gibson assembly 1341/1342 (#J) into #3 *SalI | *kan^R*, pJC1-$P_{cg1974}$-*venus* (#3) derivative with an inserted $P_{cg1974}$_sgRNA-CS1 sequence under control of the $P_{J23119}$ promoter and followed by the *rrnB* terminator behind the *venus*-*rrnB* (T) sequence | C: 554/R13<br>S: 554, R13 | This work |
| #15 | pJC1-$P_{cg1974}$-*venus*_T_$P_{J23119}$-sgRNA-33 | Gibson assembly 1341/1342 (#K) into #3 *SalI | *kan^R*, pJC1-$P_{cg1974}$-*venus* (#3) derivative with an inserted $P_{cg1974}$_sgRNA-33 sequence under control of the $P_{J23119}$ promoter and followed by the *rrnB* terminator behind the *venus*-*rrnB* (T) sequence | C: 554/R13<br>S: 554, R13 | This work |
| #16 | pJC1-$P_{cg1974}$-*venus*_T_$P_{J23119}$-sgRNA-34 | Gibson assembly 1341/1342 (#L) into #3 *SalI | *kan^R*, pJC1-$P_{cg1974}$-*venus* (#3) derivative with an inserted $P_{cg1974}$_sgRNA-34 sequence under control of the $P_{J23119}$ promoter and followed by the *rrnB* terminator behind the *venus*-*rrnB* (T) sequence | C: 554/R13<br>S: 554, R13 | This work |

| | | | | | |
|---|---|---|---|---|---|
| #17 | pJC1-$P_{cg1974}$-*venus*_T_$P_{J23119}$-sgRNA-35 | Gibson assembly 1341/1342 (#M) into #3 *SalI | *kan*$^R$, pJC1-$P_{cg1974}$-*venus* (#3) derivative with an inserted $P_{cg1974}$_sgRNA-35 sequence under control of the $P_{J23119}$ promoter and followed by the *rrnB* terminator behind the *venus*-*rrnB* (T) sequence | C: 554/R13<br>S: 554, R13 | This work |
| #18 | pJC1-$P_{cg1974}$-*venus*_T_$P_{J23119}$-sgRNA-36 | Gibson assembly 1341/1342 (#N) into #3 *SalI | *kan*$^R$, pJC1-$P_{cg1974}$-*venus* (#3) derivative with an inserted $P_{cg1974}$_sgRNA-36 sequence under control of the $P_{J23119}$ promoter and followed by the *rrnB* terminator behind the *venus*-*rrnB* (T) sequence | C: 554/R13<br>S: 554, R13 | This work |
| #19 | pJC1-$P_{cg1974}$-*venus*_T_$P_{J23119}$-sgRNA-37 | Gibson assembly 1341/1342 (#O) into #3 *SalI | *kan*$^R$, pJC1-$P_{cg1974}$-*venus* (#3) derivative with an inserted $P_{cg1974}$_sgRNA-37 sequence under control of the $P_{J23119}$ promoter and followed by the *rrnB* terminator behind the *venus*-*rrnB* (T) sequence | C: 554/R13<br>S: 554, R13 | This work |

| | | | | | |
|---|---|---|---|---|---|
| #20 | pJC1-$P_{cg1974}$-*venus*_T_$P_{J23119}$-sgRNA-38 | Gibson assembly 1341/1342 (#P) into #3 *SalI | *kan*$^R$, pJC1-$P_{cg1974}$-*venus* (#3) derivative with an inserted $P_{cg1974}$_sgRNA-38 sequence under control of the $P_{J23119}$ promoter and followed by the *rrnB* terminator behind the *venus*-*rrnB* (T) sequence | C: 554/R13<br>S: 554, R13 | This work |
| #21 | pJC1-$P_{cg1974}$-*venus*_T_$P_{J23119}$-sgRNA-39 | Gibson assembly 1341/1342 (#Q) into #3 *SalI | *kan*$^R$, pJC1-$P_{cg1974}$-*venus* (#3) derivative with an inserted $P_{cg1974}$_sgRNA-39 sequence under control of the $P_{J23119}$ promoter and followed by the *rrnB* terminator behind the *venus*-*rrnB* (T) sequence | C: 554/R13<br>S: 554, R13 | This work |
| #22 | pJC1-$P_{cg1974}$-*venus*_T_$P_{J23119}$-sgRNA-40 | Gibson assembly 1341/1342 (#R) into #3 *SalI | *kan*$^R$, pJC1-$P_{cg1974}$-*venus* (#3) derivative with an inserted $P_{cg1974}$_sgRNA-40 sequence under control of the $P_{J23119}$ promoter and followed by the *rrnB* terminator behind the *venus*-*rrnB* (T) sequence | C: 554/R13<br>S: 554, R13 | This work |

| | | | | | |
|---|---|---|---|---|---|
| #23 | pJC1-$P_{cg2014}$-*venus*_T_$P_{J23119}$-sgRNA-22 | Gibson assembly 1341/1342 (#S) into #8 *SalI | *kan*$^R$, pJC1-$P_{cg2014}$-*venus* (#8) derivative with an inserted $P_{cg2014}$_sgRNA-22 sequence under control of the $P_{J23119}$ promoter and followed by the *rrnB* terminator behind the *venus*-*rrnB* (T) sequence | C: 554/R13<br>S: 554, R13 | This work |
| #24 | pJC1-$P_{cg2014}$-*venus*_T_$P_{J23119}$-sgRNA-24 | Gibson assembly 1341/1342 (#T) into #8 *SalI | *kan*$^R$, pJC1-$P_{cg2014}$-*venus* (#8) derivative with an inserted $P_{cg2014}$_sgRNA-24 sequence under control of the $P_{J23119}$ promoter and followed by the *rrnB* terminator behind the *venus*-*rrnB* (T) sequence | C: 554/R13<br>S: 554, R13 | This work |
| #25 | pJC1-$P_{cg2014}$-*venus*_T_$P_{J23119}$-sgRNA-25 | Gibson assembly 1341/1342 (#U) into #8 *SalI | *kan*$^R$, pJC1-$P_{cg2014}$-*venus* (#8) derivative with an inserted $P_{cg2014}$_sgRNA-25 sequence under control of the $P_{J23119}$ promoter and followed by the *rrnB* terminator behind the *venus*-*rrnB* (T) sequence | C: 554/R13<br>S: 554, R13 | This work |

| | | | | | |
|---|---|---|---|---|---|
| #26 | pJC1-$P_{cg2014}$-*venus*_T_$P_{J23119}$-sgRNA-26 | Gibson assembly 1341/1342 (#V) into #8 *SalI | *kan^R^*, pJC1-$P_{cg2014}$-*venus* (#8) derivative with an inserted $P_{cg2014}$_sgRNA-26 sequence under control of the $P_{J23119}$ promoter and followed by the *rrnB* terminator behind the *venus-rrnB* (T) sequence | C: 554/R13<br>S: 554, R13 | This work |
| #27 | pJC1-$P_{cg2014}$-*venus*_T_$P_{J23119}$-sgRNA-27 | Gibson assembly 1341/1342 (#W) into #8 *SalI | *kan^R^*, pJC1-$P_{cg2014}$-*venus* (#8) derivative with an inserted $P_{cg2014}$_sgRNA-27 sequence under control of the $P_{J23119}$ promoter and followed by the *rrnB* terminator behind the *venus-rrnB* (T) sequence | C: 554/R13<br>S: 554, R13 | This work |
| #28 | pJC1-$P_{cg2014}$-*venus*_T_$P_{J23119}$-sgRNA-28 | Gibson assembly 1341/1342 (#X) into #8 *SalI | *kan^R^*, pJC1-$P_{cg2014}$-*venus* (#8) derivative with an inserted $P_{cg2014}$_sgRNA-28 sequence under control of the $P_{J23119}$ promoter and followed by the *rrnB* terminator behind the *venus-rrnB* (T) sequence | C: 554/R13<br>S: 554, R13 | This work |

| | | | | | |
|---|---|---|---|---|---|
| #29 | pJC1-$P_{cg2014}$-*venus*_T_$P_{J23119}$-sgRNA-29 | Gibson assembly 1341/1342 (#Y) into #8 *SalI | *kan*$^R$, pJC1-$P_{cg2014}$-*venus* (#8) derivative with an inserted $P_{cg2014}$_sgRNA-29 sequence under control of the $P_{J23119}$ promoter and followed by the *rrnB* terminator behind the *venus*-*rrnB* (T) sequence | C: 554/R13<br>S: 554, R13 | This work |
| #30 | pJC1-$P_{cg2014}$-*venus*_T_$P_{J23119}$-sgRNA-30 | Gibson assembly 1341/1342 (#Z) into #8 *SalI | *kan*$^R$, pJC1-$P_{cg2014}$-*venus* (#8) derivative with an inserted $P_{cg2014}$_sgRNA-30 sequence under control of the $P_{J23119}$ promoter and followed by the *rrnB* terminator behind the *venus*-*rrnB* (T) sequence | C: 554/R13<br>S: 554, R13 | This work |
| #31 | pJC1-$P_{cg2014}$-*venus*_T_$P_{J23119}$-sgRNA-31 | Gibson assembly 1341/1342 (#AA) into #8 *SalI | *kan*$^R$, pJC1-$P_{cg2014}$-*venus* (#8) derivative with an inserted $P_{cg2014}$_sgRNA-31 sequence under control of the $P_{J23119}$ promoter and followed by the *rrnB* terminator behind the *venus*-*rrnB* (T) sequence | C: 554/R13<br>S: 554, R13 | This work |

| | | | | | |
|---|---|---|---|---|---|
| #32 | pJC1-$P_{cg1974}$-*venus*_T_$P_{lacM}$-*dcas12a* | Gibson assembly 1778/1779 (#AB) into #3 *SalI | *$kan^R$*, pJC1-$P_{cg1974}$-*venus* (#3) derivative with an inserted *dcas12a* (*cpf1* from *Francisella novicida* carrying mutations D917A and E1006A) sequence under control of the constitutive $P_{lacM}$ promoter and followed by the *rrnB* terminator behind the *venus*-*rrnB* (T) sequence | C: 1788/1792<br>S: 1787, 1788, 1789, 1790, 1791, 1792 | This work |
| #33 | pEC-XC99E_$P_{J23119}$-$P_{cg1974}$_crRNA-7 | Gibson assembly 1782/1800 (#AC *SpeI) into 1873/1874 (#4 *SdaI) | *$cm^R$*, pEC-XC99E_empty (#4) derivative with an inserted $P_{cg1974}$_crRNA-7 sequence under control of the $P_{J23119}$ promoter in front of the *rrnB* terminator | C: 1782/1442<br>S: 1441 | This work |
| #34 | pEC-XC99E_$P_{J23119}$-$P_{cg1974}$_crRNA-1 | Gibson assembly 1782/1783 (#AC *SpeI) into 1873/1874 (#4 *SdaI) | *$cm^R$*, pEC-XC99E_empty (#4) derivative with an inserted $P_{cg1974}$_crRNA-1 sequence under control of the $P_{J23119}$ promoter in front of the *rrnB* terminator | C: 1782/1442<br>S: 1441 | This work |
| #35 | pEC-XC99E_$P_{J23119}$-$P_{cg1974}$_crRNA-3 | Gibson assembly 1782/1796 (#AC *SpeI) into | *$cm^R$*, pEC-XC99E_empty (#4) derivative with an inserted $P_{cg1974}$_crRNA-3 | C: 1782/1442<br>S: 1441 | This work |

| | | | | | |
|---|---|---|---|---|---|
| | | 1873/1874 (#4 *SdaI) | sequence under control of the $P_{J23119}$ promoter in front of the *rrnB* terminator | | |
| #36 | pEC-XC99E_$P_{J23119}$-$P_{cg1974}$_crRNA-4 | Gibson assembly 1782/1797 (#AC *SpeI) into 1873/1874 (#4 *SdaI) | $cm^R$, pEC-XC99E_empty (#4) derivative with an inserted $P_{cg1974}$_crRNA-4 sequence under control of the $P_{J23119}$ promoter in front of the *rrnB* terminator | C: 1782/1442<br>S: 1441 | This work |
| #37 | pEC-XC99E_$P_{J23119}$-$P_{cg1974}$_crRNA-5 | Gibson assembly 1782/1798 (#AC *SpeI) into 1873/1874 (#4 *SdaI) | $cm^R$, pEC-XC99E_empty (#4) derivative with an inserted $P_{cg1974}$_crRNA-5 sequence under control of the $P_{J23119}$ promoter in front of the *rrnB* terminator | C: 1782/1442<br>S: 1441 | This work |
| #38 | pEC-XC99E_$P_{J23119}$-$P_{cg1974}$_crRNA-6 | Gibson assembly 1782/1799 (#AC *SpeI) into 1873/1874 (#4 *SdaI) | $cm^R$, pEC-XC99E_empty (#4) derivative with an inserted $P_{cg1974}$_crRNA-6 sequence under control of the $P_{J23119}$ promoter in front of the *rrnB* terminator | C: 1782/1442<br>S: 1441 | This work |
| #39 | pEC-XC99E_$P_{J23119}$-$P_{cg1974}$_crRNA-8 | Gibson assembly 1782/1801 (#AC *SpeI) into 1873/1874 (#4 *SdaI) | $cm^R$, pEC-XC99E_empty (#4) derivative with an inserted $P_{cg1974}$_crRNA-8 sequence under control of the $P_{J23119}$ | C: 1782/1442<br>S: 1441 | This work |

| | | | | | |
|---|---|---|---|---|---|
| | | | promoter in front of the *rrnB* terminator | | |
| #40 | pEC-XC99E_$P_{J23119}$-$P_{cg1974}$_crRNA-9 | Gibson assembly 1782/1802 (#AC *SpeI) into 1873/1874 (#4 *SdaI) | $cm^R$, pEC-XC99E_empty (#4) derivative with an inserted $P_{cg1974}$_crRNA-9 sequence under control of the $P_{J23119}$ promoter in front of the *rrnB* terminator | C: 1782/1442<br>S: 1441 | This work |
| #41 | pEC-XC99E_$P_{J23119}$-$P_{cg1974}$_crRNA-10 | Gibson assembly 1782/1803 (#AC *SpeI) into 1873/1874 (#4 *SdaI) | $cm^R$, pEC-XC99E_empty (#4) derivative with an inserted $P_{cg1974}$_crRNA-10 sequence under control of the $P_{J23119}$ promoter in front of the *rrnB* terminator | C: 1782/1442<br>S: 1441 | This work |
| #42 | pJC1-$P_{cg2014}$-*venus*_T_$P_{lacM}$-*dcas12a* | Gibson assembly 1778/1779 (#AB) into #8 *SalI | $kan^R$, pJC1-$P_{cg2014}$-*venus* (#8) derivative with an inserted *dcas12a* (*cpf1* from *Francisella novicida* carrying mutations D917A and E1006A) sequence under control of the constitutive $P_{lacM}$ promoter and followed by the *rrnB* terminator behind the *venus*-*rrnB* (T) sequence | C: 1788/1792<br>S: 1787, 1788, 1789, 1790, 1791, 1792 | This work |

| | | | | | |
|---|---|---|---|---|---|
| #43 | pEC-XC99E_$P_{J23119}$-$P_{cg2014}$_crRNA-1 | Gibson assembly 1782/1875 (#AC *SpeI) into 1873/1874 (#4 *SdaI) | *$cm^R$*, pEC-XC99E_empty (#4) derivative with an inserted $P_{cg2014}$_crRNA-1 sequence under control of the $P_{J23119}$ promoter in front of the *rrnB* terminator | C: 1782/1442<br>S: 1441 | This work |
| #44 | pEC-XC99E_$P_{J23119}$-$P_{cg2014}$_crRNA-2 | Gibson assembly 1782/1876 (#AC *SpeI) into 1873/1874 (#4 *SdaI) | *$cm^R$*, pEC-XC99E_empty (#4) derivative with an inserted $P_{cg2014}$_crRNA-2 sequence under control of the $P_{J23119}$ promoter in front of the *rrnB* terminator | C: 1782/1442<br>S: 1441 | This work |
| #45 | pEC-XC99E_$P_{J23119}$-$P_{cg2014}$_crRNA-3 | Gibson assembly 1782/1877 (#AC *SpeI) into 1873/1874 (#4 *SdaI) | *$cm^R$*, pEC-XC99E_empty (#4) derivative with an inserted $P_{cg2014}$_crRNA-3 sequence under control of the $P_{J23119}$ promoter in front of the *rrnB* terminator | C: 1782/1442<br>S: 1441 | This work |
| #46 | pEC-XC99E_$P_{J23119}$-$P_{cg2014}$_crRNA-4 | Gibson assembly 1782/1878 (#AC *SpeI) into 1873/1874 (#4 *SdaI) | *$cm^R$*, pEC-XC99E_empty (#4) derivative with an inserted $P_{cg2014}$_crRNA-4 sequence under control of the $P_{J23119}$ promoter in front of the *rrnB* terminator | C: 1782/1442<br>S: 1441 | This work |

| | | | | | |
|---|---|---|---|---|---|
| #47 | pEC-XC99E_P$_{J23119}$-P$_{cg2014}$_crRNA-6 | Gibson assembly 1782/1880 (#AC *SpeI) into 1873/1874 (#4 *SdaI) | *cm*$^R$, pEC-XC99E_empty (#4) derivative with an inserted P$_{cg2014}$_crRNA-6 sequence under control of the P$_{J23119}$ promoter in front of the *rrnB* terminator | C: 1782/1442<br>S: 1441 | This work |
| #48 | pEC-XC99E_P$_{J23119}$-P$_{cg2014}$_crRNA-7 | Gibson assembly 1782/1881 (#AC *SpeI) into 1873/1874 (#4 *SdaI) | *cm*$^R$, pEC-XC99E_empty (#4) derivative with an inserted P$_{cg2014}$_crRNA-7 sequence under control of the P$_{J23119}$ promoter in front of the *rrnB* terminator | C: 1782/1442<br>S: 1441 | This work |
| #49 | pJC1-P$_{cg1896}$-*venus* | Gibson assembly 789/722 (genomic *C. glutamicum* DNA) and 115/116 (#74) into #74 *BamHI *SpeI | *kan*$^R$, pJC1-*venus* (#74) derivative carrying the CgpS bound area of the promoter of cg1896 (185 bp) and the first 30 bp of the coding sequence fused to the reporter gene *venus* via a linker containing a stop codon and an artificial RBS | C: R12/R13<br>S: R12, R13 | This work |
| #50 | pJC1-P$_{cg1929}$-*venus* | Gibson assembly 792/723 (genomic *C. glutamicum* DNA) and 115/116 (#74) | *kan*$^R$, pJC1-*venus* (#74) derivative carrying the CgpS bound area of the promoter of cg1929 (441 bp) and the first 30 bp of the | C: R12/R13<br>S: R12, R13 | This work |

| | | | | |
|---|---|---|---|---|
| | | into #74 *BamHI *SpeI | coding sequence fused to the reporter gene *venus* via a linker containing a stop codon and an artificial RBS | |
| #51 | pJC1-$P_{cg1940}$-*venus* | | *kan*$^{R}$, pJC1-based plasmid carrying the CgpS bound area of the promoter of cg1940 (563 bp) and the first 30 bp of the coding sequence fused to the reporter gene *venus* via a linker containing a stop codon and an artificial RBS | (5) |
| #52 | pJC1-$P_{cg1977}$-*venus* | | *kan*$^{R}$, pJC1-based plasmid carrying the CgpS bound area of the promoter of cg1977 (653 bp) and the first 30 bp of the coding sequence fused to the reporter gene *venus* via a linker containing a stop codon and an artificial RBS | (5) |
| #53 | pJC1-$P_{cg1999}$-*venus* | | *kan*$^{R}$, pJC1-based plasmid carrying the CgpS bound area of the | (5) |

| | | | | | |
|---|---|---|---|---|---|
| | | | promoter of cg1999 (448 bp) and the first 30 bp of the coding sequence fused to the reporter gene *venus* via a linker containing a stop codon and an artificial RBS | | |
| #54 | pJC1-$P_{cg2004}$-*venus* | Gibson assembly 790/724 (genomic *C. glutamicum* DNA) and 115/116 (#74) into #74 *BamHI *SpeI | *kan*$^R$, pJC1-*venus* (#74) derivative carrying the CgpS bound area of the promoter of cg2004 (473 bp) and the first 30 bp of the coding sequence fused to the reporter gene *venus* via a linker containing a stop codon and an artificial RBS | C: R12/R13<br>S: R12, R13 | This work |
| #55 | pJC1-$P_{cg2011}$-*venus* | Gibson assembly 791/725 (genomic *C. glutamicum* DNA) and 115/116 (#74) into #74 *BamHI *SpeI | *kan*$^R$, pJC1-*venus* (#74) derivative carrying the CgpS bound area of the promoter of cg2011 (363 bp) and the first 30 bp of the coding sequence fused to the reporter gene *venus* via a linker containing a stop codon and an artificial RBS | C: R12/R13<br>S: R12, R13 | This work |

| | | | | | |
|---|---|---|---|---|---|
| #56 | pJC1-$P_{cg2032}$-*venus* | | *kan*$^R$, pJC1-based plasmid carrying the CgpS bound area of the promoter of cg2032 (490 bp) and the first 30 bp of the coding sequence fused to the reporter gene *venus* via a linker containing a stop codon and an artificial RBS | | (5) |
| #57 | pJC1-$P_{cg2065}$-*venus* | Gibson assembly 793/726 (genomic *C. glutamicum* DNA) and 115/116 (#74) into #74 *BamHI *SpeI | *kan*$^R$, pJC1-*venus* (#74) derivative carrying the CgpS bound area of the promoter of cg2065 (480 bp) and the first 30 bp of the coding sequence fused to the reporter gene *venus* via a linker containing a stop codon and an artificial RBS | C: R12/R13<br>S: R12, R13 | This work |
| #58 | pJC1-$P_{cg1896}$-*venus*_T_$P_{J23119}$-sgRNA-1 | Gibson assembly 1341/1342 (#AD) into #49 *SalI | *kan*$^R$, pJC1-$P_{cg1896}$-*venus* (#49) derivative with an inserted $P_{cg1896}$_sgRNA-1 sequence under control of the $P_{J23119}$ promoter and followed by the *rrnB* terminator | C: 554/R13<br>S: 554, R13 | This work |

| | | | | | |
|---|---|---|---|---|---|
| | | | behind the *venus*-*rrnB* (T) sequence | | |
| #59 | pJC1-$P_{cg1896}$-*venus*_T_$P_{J23119}$-sgRNA-2 | Gibson assembly 1341/1342 (#AE) into #49 *SalI | *kan*$^R$, pJC1-$P_{cg1896}$-*venus* (#49) derivative with an inserted $P_{cg1896}$_sgRNA-2 sequence under control of the $P_{J23119}$ promoter and followed by the *rrnB* terminator behind the *venus*-*rrnB* (T) sequence | C: 554/R13<br>S: 554, R13 | This work |
| #60 | pJC1-$P_{cg1929}$-*venus*_T_$P_{J23119}$-sgRNA-1 | Gibson assembly 1341/1342 (#AF) into #50 *SalI | *kan*$^R$, pJC1-$P_{cg1929}$-*venus* (#50) derivative with an inserted $P_{cg1929}$_sgRNA-1 sequence under control of the $P_{J23119}$ promoter and followed by the *rrnB* terminator behind the *venus*-*rrnB* (T) sequence | C: 554/R13<br>S: 554, R13 | This work |
| #61 | pJC1-$P_{cg1940}$-*venus*_T_$P_{J23119}$-sgRNA-1 | Gibson assembly 1341/1342 (#AG) into #51 *SalI | *kan*$^R$, pJC1-$P_{cg1940}$-*venus* (#51) derivative with an inserted $P_{cg1940}$_sgRNA-1 sequence under control of the $P_{J23119}$ promoter and followed by the *rrnB* terminator behind the *venus*-*rrnB* (T) sequence | C: 554/R13<br>S: 554, R13 | This work |

| | | | | | |
|---|---|---|---|---|---|
| #62 | pJC1-$P_{cg1940}$-*venus*_T_$P_{J23119}$-sgRNA-2 | Gibson assembly 1341/1342 (#AH) into #51 *SalI | *$kan^R$*, pJC1-$P_{cg1940}$-*venus* (#51) derivative with an inserted $P_{cg1940}$_sgRNA-2 sequence under control of the $P_{J23119}$ promoter and followed by the *rrnB* terminator behind the *venus*-*rrnB* (T) sequence | C: 554/R13<br>S: 554, R13 | This work |
| #64 | pJC1-$P_{cg1977}$-*venus*_T_$P_{J23119}$-sgRNA-1 | Gibson assembly 1341/1342 (#AJ) into #52 *SalI | *$kan^R$*, pJC1-$P_{cg1977}$-*venus* (#52) derivative with an inserted $P_{cg1977}$_sgRNA-1 sequence under control of the $P_{J23119}$ promoter and followed by the *rrnB* terminator behind the *venus*-*rrnB* (T) sequence | C: 554/R13<br>S: 554, R13 | This work |
| #65 | pJC1-$P_{cg1999}$-*venus*_T_$P_{J23119}$-sgRNA-1 | Gibson assembly 1341/1342 (#AK) into #53 *SalI | *$kan^R$*, pJC1-$P_{cg1999}$-*venus* (#53) derivative with an inserted $P_{cg1999}$_sgRNA-1 sequence under control of the $P_{J23119}$ promoter and followed by the *rrnB* terminator behind the *venus*-*rrnB* (T) sequence | C: 554/R13<br>S: 554, R13 | This work |

| | | | | | |
|---|---|---|---|---|---|
| #66 | pJC1-$P_{cg1999}$-*venus*_T_$P_{J23119}$-sgRNA-2 | Gibson assembly 1341/1342 (#AL) into #53 *SalI | *kan*$^R$, pJC1-$P_{cg1999}$-*venus* (#53) derivative with an inserted $P_{cg1999}$_sgRNA-2 sequence under control of the $P_{J23119}$ promoter and followed by the *rrnB* terminator behind the *venus*-*rrnB* (T) sequence | C: 554/R13<br>S: 554, R13 | This work |
| #67 | pJC1-$P_{cg2004}$-*venus*_T_$P_{J23119}$-sgRNA-1 | Gibson assembly 1341/1342 (#AM) into #54 *SalI | *kan*$^R$, pJC1-$P_{cg2004}$-*venus* (#54) derivative with an inserted $P_{cg2004}$_sgRNA-1 sequence under control of the $P_{J23119}$ promoter and followed by the *rrnB* terminator behind the *venus*-*rrnB* (T) sequence | C: 554/R13<br>S: 554, R13 | This work |
| #68 | pJC1-$P_{cg2004}$-*venus*_T_$P_{J23119}$-sgRNA-2 | Gibson assembly 1341/1342 (#AN) into #54 *SalI | *kan*$^R$, pJC1-$P_{cg2004}$-*venus* (#54) derivative with an inserted $P_{cg2004}$_sgRNA-2 sequence under control of the $P_{J23119}$ promoter and followed by the *rrnB* terminator behind the *venus*-*rrnB* (T) sequence | C: 554/R13<br>S: 554, R13 | This work |

| | | | | | |
|---|---|---|---|---|---|
| #69 | pJC1-$P_{cg2011}$-*venus*_T_$P_{J23119}$-sgRNA-1 | Gibson assembly 1341/1342 (#AO) into #55 *SalI | *kan^R*, pJC1-$P_{cg2011}$-*venus* (#55) derivative with an inserted $P_{cg2011}$_sgRNA-1 sequence under control of the $P_{J23119}$ promoter and followed by the *rrnB* terminator behind the *venus*-*rrnB* (T) sequence | C: 554/R13<br>S: 554, R13 | This work |
| #70 | pJC1-$P_{cg2014}$-*venus*_T_$P_{J23119}$-sgRNA-1 | Gibson assembly 1341/1342 (#AP) into #8 *SalI | *kan^R*, pJC1-$P_{cg2014}$-*venus* (#8) derivative with an inserted $P_{cg2014}$_sgRNA-1 sequence under control of the $P_{J23119}$ promoter and followed by the *rrnB* terminator behind the *venus*-*rrnB* (T) sequence | C: 554/R13<br>S: 554, R13 | This work |
| #71 | pJC1-$P_{cg2032}$-*venus*_T_$P_{J23119}$-sgRNA-1 | Gibson assembly 1341/1342 (#AQ) into #56 *SalI | *kan^R*, pJC1-$P_{cg2032}$-*venus* (#56) derivative with an inserted $P_{cg2032}$_sgRNA-1 sequence under control of the $P_{J23119}$ promoter and followed by the *rrnB* terminator behind the *venus*-*rrnB* (T) sequence | C: 554/R13<br>S: 554, R13 | This work |

| | | | | | |
|---|---|---|---|---|---|
| #72 | pJC1-$P_{cg2032}$-*venus*_T_$P_{J23119}$-sgRNA-2 | Gibson assembly 1341/1342 (#AR) into #56 *SalI | *kan*$^R$, pJC1-$P_{cg2032}$-*venus* (#56) derivative with an inserted $P_{cg2032}$_sgRNA-2 sequence under control of the $P_{J23119}$ promoter and followed by the *rrnB* terminator behind the *venus*-*rrnB* (T) sequence | C: 554/R13<br>S: 554, R13 | This work |
| #73 | pJC1-$P_{cg2065}$-*venus*_T_$P_{J23119}$-sgRNA-1 | Gibson assembly 1341/1342 (#AS) into #57 *SalI | *kan*$^R$, pJC1-$P_{cg2065}$-*venus* (#57) derivative with an inserted $P_{cg2065}$_sgRNA-1 sequence under control of the $P_{J23119}$ promoter and followed by the *rrnB* terminator behind the *venus*-*rrnB* (T) sequence | C: 554/R13<br>S: 554, R13 | This work |
| #74 | pJC1-*venus* | | *kan*$^R$, *E. coli*/*C. glutamicum* shuttle vector (pCG1 $ori_{Cg}$, pACYC177 p15A $ori_{Ec}$, promoter-less *venus* gene followed by terminator sequence from *Bacillus subtilis*) | | (17)<br>(pJC1-*venus*-term) |
| #75 | pJC1-$P_{LAC}$_tandem-$P_{cg1974}$-*venus* | 1494/116 (#3 *BamHI) | *kan*$^R$, pJC1-*venus* (#74) derivative with an inserted $P_{LAC}$ promoter | C: 766/114<br>S: 1491, R12, 555, R13 | This work |

| | | | | | |
|---|---|---|---|---|---|
| | | Into #74 *BamHI *SpeI | sequence in tandem orientation to the $P_{cg1974}$ promoter sequence that was fused to the reporter gene *venus* via a linker containing a stop codon and an artificial RBS | | |
| #76 | pJC1-$P_{J23150}$_tandem-$P_{cg1974}$-*venus* | 1493/116 (#3 *BamHI)<br>Into #74 *BamHI *SpeI | *kan*$^R$, pJC1-*venus* (#74) derivative with an inserted $P_{J23150}$ promoter sequence in tandem orientation to the $P_{cg1974}$ promoter sequence that was fused to the reporter gene *venus* via a linker containing a stop codon and an artificial RBS | C: 766/114<br>S: 1491, R12, 555, R13 | This work |
| #77 | pJC1-$P_{J23119}$_tandem-$P_{cg1974}$-*venus* | 1492/116 (#3 *BamHI)<br>Into #74 *BamHI *SpeI | *kan*$^R$, pJC1-*venus* (#74) derivative with an inserted $P_{J23119}$ promoter sequence in tandem orientation to the $P_{cg1974}$ promoter sequence that was fused to the reporter gene *venus* via a linker containing a stop codon and an artificial RBS | C: 766/114<br>S: 1491, R12, 555, R13 | This work |

| | | | | | |
|---|---|---|---|---|---|
| #78 | pJC1-$P_{LAC}$_divergent-$P_{cg1974}$-*venus* | 1497/851 (#A *SpeI)<br>Into #3 *BamHI | *kan*$^R$, pJC1-*venus* (#74) derivative with an inserted $P_{LAC}$ promoter sequence in divergent orientation to the $P_{cg1974}$ promoter sequence that was fused to the reporter gene *venus* via a linker containing a stop codon and an artificial RBS | C: R12/492<br>S: 1491, R12, 492 | This work |
| #79 | pJC1-$P_{J23150}$_divergent-$P_{cg1974}$-*venus* | 1496/851 (#A *SpeI)<br>Into #3 *BamHI | *kan*$^R$, pJC1-*venus* (#74) derivative with an inserted $P_{J23150}$ promoter sequence in divergent orientation to the $P_{cg1974}$ promoter sequence that was fused to the reporter gene *venus* via a linker containing a stop codon and an artificial RBS | C: R12/492<br>S: 1491, R12, 492 | This work |
| #80 | pJC1-$P_{J23119}$_divergent-$P_{cg1974}$-*venus* | 848/851 (#A *SpeI)<br>Into #3 *BamHI | *kan*$^R$, pJC1-*venus* (#74) derivative with an inserted $P_{J23119}$ promoter sequence in divergent orientation to the $P_{cg1974}$ promoter sequence that was fused to the | C: R12/492<br>S: 1491, R12, 492 | This work |

| | | | | | |
|---|---|---|---|---|---|
| | | | reporter gene *venus* via a linker containing a stop codon and an artificial RBS | | |
| #81 | pJC1-$P_{J23119}$_forward-*venus* | 1424/1422 (#74) and 1423/1342 (#A) into #74 *SalI | *kan*$^R$, pJC1-*venus* (#74) derivative with an inserted $P_{J23119}$ promoter sequence fused in forward orientation to the reporter gene *venus* via a linker containing an artificial RBS | C: R12/R13<br><br>S: R12, 492, R13 | This work |
| #82 | pJC1-$P_{J23119}$_reverse-*venus* | 1527/851 (#A) and 115/116 (#74) into #74 *BamHI *SpeI | *kan*$^R$, pJC1-*venus* (#74) derivative with an inserted $P_{J23119}$ promoter sequence combined with the *rrnB* terminator in reverse orientation to the reporter gene *venus* via a linker containing an artificial RBS | C: R12/R13<br><br>S: 1491, R12, 555, R13 | This work |
| #83 | pMC345 | Gibson assembly: three successive fragment additions into #AT: (i) dCas9-M13F/dCas9-TipAR (#AU) *StuI *BglII, (ii) CRISPR-pSET152-1/CRISPR- | *apr*$^R$, *thio*$^R$ pCRISPR-dCas9 (#AT) derivative with theophylline riboswitch-controlled *tipA,* integration apparatus (phage C31 *attP-int* locus), and a new, unique | S: whole plasmid sequencing by plasmidsaurus | This work |

| | | | | | |
|---|---|---|---|---|---|
| | | pSET152-2 (#AV) *ApaLI *NotI, (iii) FixNco-F3, FixNco-R/FixNco-R2 (#AT) *AseI *SnaBI | *Avr*II site for guide sequence cloning | | |
| #84 | pMC346 | Gibson assembly: guide sequence $P_{vnz04460}$_sgRNA into #83 *AvrII | *apr*$^R$, *thio*$^R$ pMC345 (#83) derivative with $P_{vnz04460}$_sgRNA | S: universal M13 Reverse | This work |
| #85 | pMC347 | Gibson assembly: guide sequence *vnz04465*_sgRNA into #83 *AvrII | *apr*$^R$, *thio*$^R$ pMC345 (#83) derivative with *vnz04465*_sgRNA | S: universal M13 Reverse | This work |

**Table S3A. Oligonucleotides used for plasmid constructions (see Table S2).**

| Oligonucleotide | Sequence (5´-3´) |
|---|---|
| B024 | AGTTTAATTTGTAGTATCCAACGAACATTAAACGGGTAAAGGTAA |
| B026 | TTTACCCGTTTAATGTTCGTTGGATACTACAAATTAAACTCTAGTA |
| OSM13 | AAGATCCCCAGCTTGTTGATACACCTGCAGGAGGAGAAAGGATCTATGG |
| OSM14 | CGGTACCCGGGGATCCTCTAGAGTCGACCCCTAGGTATAAACGCAGAAAG |
| OSM35 | CAGTTTAGCGGTCTGTTTAAGAGCTATGCTGGAAATAGCAAGTTAAAAT |
| OSM36 | ATTTTAACTTGCTATTTCCAGCATAGCTCTTAAACAGACCGCTAAACTG |
| OSM37 | TAAGAGCTATGCTGGAAACAGCATAGCAAGTTTAAATAAGGCTAGTCCGT |
| OSM38 | ACGGACTAGCCTTATTTAAACTTGCTATGCTGTTTCCAGCATAGCTCTTA |

| | |
|---|---|
| OSM39 | GCTTGTTGATACACCTGAGGAGGAGAAAGGATC |
| OSM40 | GATCCTTTCTCCTCCTCAGGTGTATCAACAAGC |
| OSM47 | GCTCAGTCCTAGGTATAATACTAGTGTTTAAGAGCTATGCTGGAAAC |
| OSM48 | GTTTCCAGCATAGCTCTTAAACACTAGTATTATACCTAGGACTGAGC |
| R65 | CATCTGTGCGGTATTTCACACCGCACCTGCAGGCATGCAAGCTTGGCTGT |
| R66 | ACAGCCAAGCTTGCATGCCTGCAGGTGCGGTGTGAAATACCGCACAGATG |
| R67 | GTGCGGTATTTCACACCGCAGTATAAACGCAGAAAGGC |
| R68 | TAGTGTTAACCCCGGGATGGATAAGAAATACTCAATAGG |
| R69 | TCTTATCCATCCCGGGGTTAACACTAGTGTTTAAGAGCTATGCTG |
| R70 | CAAGCTTGCATGCCTGCAGGGGATCCAGTTCACCGACAAAC |
| R87 | GCCTATTGAGTATTTCTTATCCATATCTATATCTCCTTGGATCCTCTAG |
| R88 | CTGTTTCCAGCATAGCTCTTAAACACTAGTGTCGACTCTAGAGGATCCTGTACAACCATAGTGGCCATGAGC |
| R91 | GCCTATTGAGTATTTCTTATCCATATCTATATCTCCTTGGATCCTCTAG |
| R92 | GATGGCTGCCTGAACCATAGTGGCCATGAGC |
| R93 | TGGCCACTATGGTTCAGGCAGCCATCGGAAG |
| R94 | CTGTTTCCAGCATAGCTCTTAAACACTAGTGTCGACTCTAGAGGATCCTGTACATTATACGAGCCGATGATTAATTGTCAAC |
| R172 | TAATCATCGGCTCGTATAATTTCTGTTACTATTAACATATGTTTAAGAGCTATGCTGGAA |
| R173 | TTCCAGCATAGCTCTTAAACATATGTTAATAGTAACAGAAATTATACGAGCCGATGATTA |
| R176 | TAATCATCGGCTCGTATAATGAGTTTAATTTGTAGTATCCGTTTAAGAGCTATGCTGGAA |
| R177 | TTCCAGCATAGCTCTTAAACGGATACTACAAATTAAACTCATTATACGAGCCGATGATTA |
| 115 | TGAACTTTAAGAAGGAGATATCATATGGTGAGCAAGGGCGAGGAG |

| | |
|---|---|
| 116 | AAAACGACGGCCAGTACTAGTTACTTGTACAGCTCGTCCATGCC |
| 117 | AGCGACGCCGCAGGGGGATCCGCTCAAGGAAGAGTTCTTCATTGGTC |
| 722 | TGATATCTCCTTCTTAAAGTTCAAAGACGACGAGAATTAAGCTTCAAAGAC |
| 723 | TGATATCTCCTTCTTAAAGTTCATGTTTGTTCCTTAAAACTACTACCAAACAT |
| 724 | TGATATCTCCTTCTTAAAGTTCACCGATGTTGTTGCGGTGCTCG |
| 725 | TGATATCTCCTTCTTAAAGTTCAGATCTCATTCTCCGGCGTTCCC |
| 726 | TGATATCTCCTTCTTAAAGTTCACATTGCTGCTGGATTACCCAACG |
| 789 | AGCGACGCCGCAGGGGGATCCGACGGTTCACAGATCACCACG |
| 790 | AGCGACGCCGCAGGGGGATCCCCCGTTAGAGTCGGCAGAAGG |
| 791 | AGCGACGCCGCAGGGGGATCCACTAGCACAAAACTACTTTGGTGGTG |
| 792 | AGCGACGCCGCAGGGGGATCCGAGCGTCCAATACTCCAAGTGTC |
| 793 | AGCGACGCCGCAGGGGGATCCTTTCGTGAAGCGGTAATGAGTTGTGC |
| 848 | CAATGAAGAACTCTTCCTTGAGCGGATCGGAATTCTAAAGATCTTTGACAGCTAGCTC |
| 851 | AGCGACGCCGCAGGGGGATCCGGATCCAGTTCACCGACAAACAAC |
| 852 | AATATTCCTTATCTATATAAGTTTAAGAGCTATGCTGGAAACAGC |
| 853 | TTATATAGATAAGGAATATTACTAGTATTATACCTAGGACTGAGCTAG |
| 854 | TTCTCACCTTTATATAGATAGTTTAAGAGCTATGCTGGAAACAGC |
| 855 | TATCTATATAAAGGTGAGAAACTAGTATTATACCTAGGACTGAGCTAG |
| 856 | TAAAGGTAGTCTTAAATGTTGTTTAAGAGCTATGCTGGAAACAGC |
| 857 | AACATTTAAGACTACCTTTAACTAGTATTATACCTAGGACTGAGCTAG |
| 858 | AAACGCGATAGGCGTGTATGGTTTAAGAGCTATGCTGGAAACAGC |
| 859 | CATACACGCCTATCGCGTTTACTAGTATTATACCTAGGACTGAGCTAG |
| 860 | TATAACCCATTCTGGATTGCGTTTAAGAGCTATGCTGGAAACAGC |
| 861 | GCAATCCAGAATGGGTTATAACTAGTATTATACCTAGGACTGAGCTAG |

| | |
|---|---|
| 864 | GAGTTTAATTTGTAGTATCCGTTTAAGAGCTATGCTGGAAACAGC |
| 865 | GGATACTACAAATTAAACTCACTAGTATTATACCTAGGACTGAGCTAG |
| 866 | GTTTATTTATGTGATTTGACGTTTAAGAGCTATGCTGGAAACAGC |
| 867 | GTCAAATCACATAAATAAACACTAGTATTATACCTAGGACTGAGCTAG |
| 868 | CTTAATTGAGTTTATTTTTAGTTTAAGAGCTATGCTGGAAACAGC |
| 869 | TAAAAATAAACTCAATTAAGACTAGTATTATACCTAGGACTGAGCTAG |
| 870 | TAATTACTTGATTTAATTGAGTTTAAGAGCTATGCTGGAAACAGC |
| 871 | TCAATTAAATCAAGTAATTAACTAGTATTATACCTAGGACTGAGCTAG |
| 872 | CTTTCAGCCTTGATAACAAGGTTTAAGAGCTATGCTGGAAACAGC |
| 873 | CTTGTTATCAAGGCTGAAAGACTAGTATTATACCTAGGACTGAGCTAG |
| 874 | AGCTTGTTTAATTAATTAAAGTTTAAGAGCTATGCTGGAAACAGC |
| 875 | TTTAATTAATTAAACAAGCTACTAGTATTATACCTAGGACTGAGCTAG |
| 876 | AAAGTACTAATATTACAGAGGTTTAAGAGCTATGCTGGAAACAGC |
| 877 | CTCTGTAATATTAGTACTTTACTAGTATTATACCTAGGACTGAGCTAG |
| 878 | AGTATTGCATTTGTTTGACAGTTTAAGAGCTATGCTGGAAACAGC |
| 879 | TGTCAAACAAATGCAATACTACTAGTATTATACCTAGGACTGAGCTAG |
| 880 | AATTCAAAGTTATTTTAGAAGTTTAAGAGCTATGCTGGAAACAGC |
| 881 | TTCTAAAATAACTTTGAATTACTAGTATTATACCTAGGACTGAGCTAG |
| 882 | ATTCTAAAATAACTTTGAATGTTTAAGAGCTATGCTGGAAACAGC |
| 883 | ATTCAAAGTTATTTTAGAATACTAGTATTATACCTAGGACTGAGCTAG |
| 884 | ATTAATATTTTATTAACTTAGTTTAAGAGCTATGCTGGAAACAGC |
| 885 | TAAGTTAATAAAATATTAATACTAGTATTATACCTAGGACTGAGCTAG |
| 1341 | GCGTTTCTACAAACTCTTTTGGGGAATTCTAAAGATCTTTGACAGCTAGCTC |
| 1342 | CGTTGTTGCCATTGCTGCAGGTCGAGGATCCAGTTCACCGACAAACAAC |
| 1422 | GTTTTTGTTCGGGCCCAAGCTTCTTACTTGTACAGCTCGTCCATGCC |

| | |
|---|---|
| 1423 | GGCATGGACGAGCTGTACAAGTAAGAAGCTTGGGCCCGAACAAAAAC |
| 1424 | GCGTTTCTACAAACTCTTTTGGGGAATTCTAAAGATCTTTGACAGCTAGCTCAGTCCTAG |
| 1492 | AGCGACGCCGCAGGGGGATCCTTGACAGCTAGCTCAGTCCTAGGTATAATACTAGTGCTCAAGGAAGAGTTCTTCATTGGTC |
| 1493 | AGCGACGCCGCAGGGGGATCCTTTACGGCTAGCTCAGTCCTAGGTATTATACTAGTGCTCAAGGAAGAGTTCTTCATTGGTC |
| 1494 | AGCGACGCCGCAGGGGGATCCATAAATGTGAGCGGATAACATTGACATTGTGAGCGGATAACAAGATACTGAGCACGCTCAAGGAAGAGTTCTTCATTGGTC |
| 1496 | CAATGAAGAACTCTTCCTTGAGCGGATCTTTACGGCTAGCTCAGTCCTAGGTATTATACTAGTGTTTAAGAGCTATGCTGGAAACAGC |
| 1497 | CAATGAAGAACTCTTCCTTGAGCGGATCATAAATGTGAGCGGATAACATTGACATTGTGAGCGGATAACAAGATACTGAGCACGTTTAAGAGCTATGCTGGAAACAGC |
| 1527 | ATGATATCTCCTTCTTAAAGTTCATTGACAGCTAGCTCAGTCCTAGG |
| 1659 | TTAAAAGTATATTTCAGCAAGTTTAAGAGCTATGCTGGAAACAGC |
| 1660 | TTGCTGAAATATACTTTTAAACTAGTATTATACCTAGGACTGAGCTAG |
| 1661 | ATACTTAATAAGAATTGTTCGTTTAAGAGCTATGCTGGAAACAGC |
| 1662 | GAACAATTCTTATTAAGTATACTAGTATTATACCTAGGACTGAGCTAG |
| 1663 | TACTTAATAAGAATTGTTCTGTTTAAGAGCTATGCTGGAAACAGC |
| 1664 | AGAACAATTCTTATTAAGTAACTAGTATTATACCTAGGACTGAGCTAG |
| 1665 | CCTTTCACACTAATTTGTTCGTTTAAGAGCTATGCTGGAAACAGC |
| 1666 | GAACAAATTAGTGTGAAAGGACTAGTATTATACCTAGGACTGAGCTAG |
| 1667 | CCCTTTCACACTAATTTGTTGTTTAAGAGCTATGCTGGAAACAGC |
| 1668 | AACAAATTAGTGTGAAAGGGACTAGTATTATACCTAGGACTGAGCTAG |
| 1669 | CCCGAACAAATTAGTGTGAAGTTTAAGAGCTATGCTGGAAACAGC |
| 1670 | TTCACACTAATTTGTTCGGGACTAGTATTATACCTAGGACTGAGCTAG |
| 1671 | CCGAACAAATTAGTGTGAAAGTTTAAGAGCTATGCTGGAAACAGC |

| | |
|---|---|
| 1672 | TTTCACACTAATTTGTTCGGACTAGTATTATACCTAGGACTGAGCTAG |
| 1673 | GTGTGAAAGGGTTTGTATCAGTTTAAGAGCTATGCTGGAAACAGC |
| 1674 | TGATACAAACCCTTTCACACACTAGTATTATACCTAGGACTGAGCTAG |
| 1675 | CATGGCTAATTTTCGAAGCAGTTTAAGAGCTATGCTGGAAACAGC |
| 1676 | TGCTTCGAAAATTAGCCATGACTAGTATTATACCTAGGACTGAGCTAG |
| 1711 | AGGTTGAGGGGTAACTGGGTGTTTAAGAGCTATGCTGGAAACAGC |
| 1712 | ACCCAGTTACCCCTCAACCTACTAGTATTATACCTAGGACTGAGCTAG |
| 1713 | CACTACACCCAGTTCTCGAAGTTTAAGAGCTATGCTGGAAACAGC |
| 1714 | TTCGAGAACTGGGTGTAGTGACTAGTATTATACCTAGGACTGAGCTAG |
| 1715 | GAAAAAGCCTTTCGAGAACTGTTTAAGAGCTATGCTGGAAACAGC |
| 1716 | AGTTCTCGAAAGGCTTTTTCACTAGTATTATACCTAGGACTGAGCTAG |
| 1717 | GGTGTAGTGATTTCTGTTGCGTTTAAGAGCTATGCTGGAAACAGC |
| 1718 | GCAACAGAAATCACTACACCACTAGTATTATACCTAGGACTGAGCTAG |
| 1719 | TTCTGTTACTATTAACATATGTTTAAGAGCTATGCTGGAAACAGC |
| 1720 | ATATGTTAATAGTAACAGAAACTAGTATTATACCTAGGACTGAGCTAG |
| 1721 | AGTTTAATTTGTAGTATCCAGTTTAAGAGCTATGCTGGAAACAGC |
| 1722 | TGGATACTACAAATTAAACTACTAGTATTATACCTAGGACTGAGCTAG |
| 1723 | AGTATCCAGGGAACATTAAAGTTTAAGAGCTATGCTGGAAACAGC |
| 1724 | TTTAATGTTCCCTGGATACTACTAGTATTATACCTAGGACTGAGCTAG |
| 1725 | GTATCCAGGGAACATTAAACGTTTAAGAGCTATGCTGGAAACAGC |
| 1726 | GTTTAATGTTCCCTGGATACACTAGTATTATACCTAGGACTGAGCTAG |
| 1727 | AGGGAACATTAAACGGGTAAGTTTAAGAGCTATGCTGGAAACAGC |
| 1728 | TTACCCGTTTAATGTTCCCTACTAGTATTATACCTAGGACTGAGCTAG |
| 1729 | CATTAAACGGGTAAAGGTAAGTTTAAGAGCTATGCTGGAAACAGC |
| 1730 | TTACCTTTACCCGTTTAATGACTAGTATTATACCTAGGACTGAGCTAG |

| | |
|---|---|
| 1731 | AGGTAAAGGACAAACGAACAGTTTAAGAGCTATGCTGGAAACAGC |
| 1732 | TGTTCGTTTGTCCTTTACCTACTAGTATTATACCTAGGACTGAGCTAG |
| 1733 | CAAACGAACATGGCGATTAAGTTTAAGAGCTATGCTGGAAACAGC |
| 1734 | TTAATCGCCATGTTCGTTTGACTAGTATTATACCTAGGACTGAGCTAG |
| 1735 | CTTCTTAAAGTTCAATTTTTGTTTAAGAGCTATGCTGGAAACAGC |
| 1736 | AAAAATTGAACTTTAAGAAGACTAGTATTATACCTAGGACTGAGCTAG |
| 1778 | GCGTTTCTACAAACTCTTTTGGTCGATGAGCTGTTTACAATTAATCATCGTGTGG |
| 1779 | CGTTGTTGCCATTGCTGCAGGTCGATCTAGACTCCATTTAAATAAAACGAAAGGCT C |
| 1780 | GGATCCGAATTTCTACTGTTGTAGATACTAGTAGATCTTTAGAATTCCAGAAATCAT CCTTAG |
| 1781 | ATCTACAACAGTAGAAATTCGGATCCATTATACCTAGGACTGAGCTAGCTGTCAAG ATCCTTACTCGAGTCTAGACTGCAGG |
| 1782 | CCATTGCTGCAGGTCGATCTAGTTGACAGCTAGCTCAGTCCTAGG |
| 1783 | CCATTGCTGCAGGTCGATCTAGTTGACAGCTAGCTCAGTCCTAGG |
| 1796 | CTGAGCCTTTCGTTTTATTTAAATGGAGTCTAGTGCAGCCAGCACCAGTCGCCATCT ACAACAGTAGAAATTCGGATC |
| 1797 | CTGAGCCTTTCGTTTTATTTAAATGGAGTCTAGTAACTGGGTAGGTCATGAGAATCT ACAACAGTAGAAATTCGGATC |
| 1798 | CTGAGCCTTTCGTTTTATTTAAATGGAGTCTAGAATCACTACACCCAGTTCTCATCTA CAACAGTAGAAATTCGGATC |
| 1799 | CTGAGCCTTTCGTTTTATTTAAATGGAGTCTAGGTCTACATAAACCTGCAACAATCT ACAACAGTAGAAATTCGGATC |
| 1800 | CTGAGCCTTTCGTTTTATTTAAATGGAGTCTAGCCTATATGTTAATAGTAACAATCTA CAACAGTAGAAATTCGGATC |
| 1801 | CTGAGCCTTTCGTTTTATTTAAATGGAGTCTAGTTAATGTTCCCTGGATACTAATCTA CAACAGTAGAAATTCGGATC |
| 1802 | CTGAGCCTTTCGTTTTATTTAAATGGAGTCTAGTTGTAGTATCCAGGGAACATATCT ACAACAGTAGAAATTCGGATC |

| | |
|---|---|
| 1803 | CTGAGCCTTTCGTTTTATTTAAATGGAGTCTAGGAACATTAAACGGGTAAAGGATCT ACAACAGTAGAAATTCGGATC |
| 1873 | GACTCCATTTAAATAAAACGAAAGGCTCAGTCGAAAGACT |
| 1874 | CTAGATCGACCTGCAGCAATGGTGCAGGTGCGGTGTGAAATACCG |
| 1875 | CTGAGCCTTTCGTTTTATTTAAATGGAGTCTAGGAAATATACTTTTAAATTAAATCTA CAACAGTAGAAATTCGGATC |
| 1876 | CTGAGCCTTTCGTTTTATTTAAATGGAGTCTAGCCATTGCTGAAATATACTTTATCTA CAACAGTAGAAATTCGGATC |
| 1877 | CTGAGCCTTTCGTTTTATTTAAATGGAGTCTAGAAAGTATATTTCAGCAATGGATCT ACAACAGTAGAAATTCGGATC |
| 1878 | CTGAGCCTTTCGTTTTATTTAAATGGAGTCTAGGAGATTCATTTACCATTGCTATCTA CAACAGTAGAAATTCGGATC |
| 1880 | CTGAGCCTTTCGTTTTATTTAAATGGAGTCTAGACAAATGCAATACTTTTCTTATCTA CAACAGTAGAAATTCGGATC |
| 1881 | CTGAGCCTTTCGTTTTATTTAAATGGAGTCTAGGTGAAGCATAATCACCTTGTATCT ACAACAGTAGAAATTCGGATC |
| dCas9-M13F | GGTCGATCCCCGCATATAGGTTGTAAAACGACGGCCAGTG |
| dCas9-TipAR | TACCGAGCTCGAATTCCCCATCTCACCAAGACGCTGGTCG |
| CRISPR-pSET152-1 | TCGCTCCAAGCTGGGCTGTGTGCACGA |
| CRISPR-pSET152-2 | CTCGACCTGGGCGACGCGGCAAATTCCCCAATGTCAAGCAC |
| FixNco-F3 | GCGCGTTGGCCGATTCATTAATG |
| FixNco-R | GATCGACGGCCTAGGGTTTTAGAGCTAGAAATAG |
| FixNco-R2 | ACATCGTAGCTGACGCCTAC |

### Table S3B. Oligonucleotides used for sequencing and colony PCR (see Table S2).

| **Oligonucleotide** | **Sequence (5´-3´)** |
|---|---|

| | |
|---|---|
| R12 | CAGGGACAAGCCACCCGCACA |
| R13 | GGAAGCTAGAGTAAGTAGTTCGC |
| R71 | CGAGTCAGTGAGCGAGGAAG |
| R72 | CAACGTTCAAATCCGCTCCC |
| R73 | ACACGCTTAGAAAATTCACTGATT |
| R74 | TCACCATAGACAAACTCCGATTCA |
| R75 | ACTTTGACCAATTCATCAACAACT |
| R76 | TTGTTTCTTCAGACTTCCGAGT |
| R77 | AGCATCTACTCCACTTGCGT |
| R78 | AGGCTAGTCCGTTATCAACTTGA |
| R95 | TTGTGCCGATAGCTAAGCCT |
| R96 | AAACTTGCTATGCTGTTTCCAG |
| R169 | CGACTCGGTGCCACTTTTTC |
| R171 | CAGCCACGTTTCTGCGAAAA |
| 114 | TGATATCTCCTTCTTAAAGTTCAATTTTTCGGCATTGCGCCTTTAATCGC |
| 492 | CTCGAACTTCACCTCGGCGC |
| 554 | GCGCACCATCTTCTTCAAGG |
| 555 | CGGCGGTGATATAGACGTTGTG |
| 766 | GACAGCTAGCTCAGTCCTAGGTATAATACTAGT |
| 890 | GGAATAAGGGCGACACGGAAATG |
| 891 | GAAAATCTTCTCTCATCCGCC |
| 1441 | CCCCTGATTCTGTGGATAACCG |
| 1442 | GGCAGTTTATGGCGGGCGTC |
| 1491 | GCCGTAGAGCGATTGAAGACC |
| 1782 | CCATTGCTGCAGGTCGATCTAGTTGACAGCTAGCTCAGTCCTAGG |

| | |
|---|---|
| 1787 | GAGGTGGCGTTCGCCGCGAGCGATGGACAGGATGTGCACGTC |
| 1788 | CTCGGCATGGACGAGCTGTAC |
| 1789 | CTGCATGGTGGTGACCACATC |
| 1790 | CTCTACTCCCAGCAGATCAACG |
| 1791 | GTTCGAGTACGACCTGATCAAGG |
| 1792 | GTGCTGCAATGATACCGCGAG |
| universal M13 Reverse | CAGGAAACAGCTATGAC |

### Table S3C. Oligonucleotides used for qPCR.

| Oligonucleotide | Sequence (5´-3´) | Target |
|---|---|---|
| B040 | CCCACGTTCACCCCACAAACG | End of CGP3 prophage element |
| B041 | CTAAAATGAAGCCATCGCGACC | End of CGP3 prophage element |
| B042 | ACGTGCTGTTCCTGTGCATGG | *ddh* (NCgl2528, cg2900) control gene |
| B043 | GCTCGGCTAAGACTGCCGCT | *ddh* (NCgl2528, cg2900) control gene |

### Table S4. Overview of guide RNA spacer sequences.

Spacer name, target region and the spacer sequence of the guide RNAs are listed. Additionally, the orientation of the PAM sequence (5´-spacer-NGG-3´ for sgRNAs, 5´-(T)TTN-spacer-3´ for crRNAs) to the spacer binding position is given.

| **Spacer** | **Gene locus of target promoter** | **Sequence (5´-3´)** | **Orientation spacer binding position–PAM sequence (5´-3´)** |
|---|---|---|---|
| $P_{cg1896}$_sgRNA-1 | cg1896 | AATATTCCTTATCTATATAA | 5´-spacer-NGG-3´ |
| $P_{cg1896}$_sgRNA-2 | cg1896 | TTCTCACCTTTATATAGATA | 5´-spacer-NGG-3´ |
| $P_{cg1929}$_sgRNA-1 | cg1929 | TAAAGGTAGTCTTAAATGTT | 5´-spacer-NGG-3´ |
| $P_{cg1940}$_sgRNA-1 | cg1940 | AAACGCGATAGGCGTGTATG | 5´-spacer-NGG-3´ |
| $P_{cg1940}$_sgRNA-2 | cg1940 | TATAACCCATTCTGGATTGC | 5´-spacer-NGG-3´ |
| $P_{cg1974}$_sgRNA-CS1 | cg1974 | TTCTGTTACTATTAACATAT | 5´-spacer-NGG-3´ |
| $P_{cg1974}$_sgRNA-CS3 | cg1974 | GAGTTTAATTTGTAGTATCC | 5´-spacer-NGG-3´ |
| $P_{cg1974}$_sgRNA-25 | cg1974 | AGGTTGAGGGGTAACTGGGT | 5´-spacer-NGG-3´ |
| $P_{cg1974}$_sgRNA-26 | cg1974 | CACTACACCCAGTTCTCGAA | 5´-spacer-NGG-3´ |
| $P_{cg1974}$_sgRNA-28 | cg1974 | GAAAAAGCCTTTCGAGAACT | 5´-spacer-NGG-3´ |
| $P_{cg1974}$_sgRNA-29 | cg1974 | GGTGTAGTGATTTCTGTTGC | 5´-spacer-NGG-3´ |
| $P_{cg1974}$_sgRNA-33 | cg1974 | AGTTTAATTTGTAGTATCCA | 5´-spacer-NGG-3´ |
| $P_{cg1974}$_sgRNA-34 | cg1974 | AGTATCCAGGGAACATTAAA | 5´-spacer-NGG-3´ |
| $P_{cg1974}$_sgRNA-35 | cg1974 | GTATCCAGGGAACATTAAAC | 5´-spacer-NGG-3´ |
| $P_{cg1974}$_sgRNA-36 | cg1974 | AGGGAACATTAAACGGGTAA | 5´-spacer-NGG-3´ |
| $P_{cg1974}$_sgRNA-37 | cg1974 | CATTAAACGGGTAAAGGTAA | 5´-spacer-NGG-3´ |
| $P_{cg1974}$_sgRNA-38 | cg1974 | AGGTAAAGGACAAACGAACA | 5´-spacer-NGG-3´ |
| $P_{cg1974}$_sgRNA-39 | cg1974 | CAAACGAACATGGCGATTAA | 5´-spacer-NGG-3´ |
| $P_{cg1974}$_sgRNA-40 | cg1974 | CTTCTTAAAGTTCAATTTTT | 5´-spacer-NGG-3´ |
| $P_{cg1974}$_crRNA-1 | cg1974 | ATTTGTAGTATCCAGGGAAC | 5´-TTTN-spacer-3´ |
| $P_{cg1974}$_crRNA-3 | cg1974 | GGCGACTGGTGCTGGCTGCA | 5´-TTTN-spacer-3´ |
| $P_{cg1974}$_crRNA-4 | cg1974 | TCTCATGACCTACCCAGTTA | 5´-TTTN-spacer-3´ |

| | | | |
|---|---|---|---|
| $P_{cg1974}$_crRNA-5 | cg1974 | GAGAACTGGGTGTAGTGATT | 5´-TTTN-spacer-3´ |
| $P_{cg1974}$_crRNA-6 | cg1974 | TGTTGCAGGTTTATGTAGAC | 5´-TTTN-spacer-3´ |
| $P_{cg1974}$_crRNA-7 | cg1974 | TGTTACTATTAACATATAGG | 5´-TTTN-spacer-3´ |
| $P_{cg1974}$_crRNA-8 | cg1974 | TAGTATCCAGGGAACATTAA | 5´-TTTN-spacer-3´ |
| $P_{cg1974}$_crRNA-9 | cg1974 | ATGTTCCCTGGATACTACAA | 5´-TTTN-spacer-3´ |
| $P_{cg1974}$_crRNA-10 | cg1974 | CCTTTACCCGTTTAATGTTC | 5´-TTTN-spacer-3´ |
| $P_{cg1977}$_sgRNA-1 | cg1977 | GTTTATTTATGTGATTTGAC | 5´-spacer-NGG-3´ |
| $P_{cg1999}$_sgRNA-1 | cg1999 | CTTAATTGAGTTTATTTTTA | 5´-spacer-NGG-3´ |
| $P_{cg1999}$_sgRNA-2 | cg1999 | TAATTACTTGATTTAATTGA | 5´-spacer-NGG-3´ |
| $P_{cg2004}$_sgRNA-1 | cg2004 | CTTTCAGCCTTGATAACAAG | 5´-spacer-NGG-3´ |
| $P_{cg2004}$_sgRNA-2 | cg2004 | AGCTTGTTTAATTAATTAAA | 5´-spacer-NGG-3´ |
| $P_{cg2011}$_sgRNA-1 | cg2011 | AAAGTACTAATATTACAGAG | 5´-spacer-NGG-3´ |
| $P_{cg2014}$_sgRNA-1 | cg2014 | AGTATTGCATTTGTTTGACA | 5´-spacer-NGG-3´ |
| $P_{cg2014}$_sgRNA-22 | cg2014 | TTAAAAGTATATTTCAGCAA | 5´-spacer-NGG-3´ |
| $P_{cg2014}$_sgRNA-24 | cg2014 | ATACTTAATAAGAATTGTTC | 5´-spacer-NGG-3´ |
| $P_{cg2014}$_sgRNA-25 | cg2014 | TACTTAATAAGAATTGTTCT | 5´-spacer-NGG-3´ |
| $P_{cg2014}$_sgRNA-26 | cg2014 | CCTTTCACACTAATTTGTTC | 5´-spacer-NGG-3´ |
| $P_{cg2014}$_sgRNA-27 | cg2014 | CCCTTTCACACTAATTTGTT | 5´-spacer-NGG-3´ |
| $P_{cg2014}$_sgRNA-28 | cg2014 | CCCGAACAAATTAGTGTGAA | 5´-spacer-NGG-3´ |
| $P_{cg2014}$_sgRNA-29 | cg2014 | CCGAACAAATTAGTGTGAAA | 5´-spacer-NGG-3´ |
| $P_{cg2014}$_sgRNA-30 | cg2014 | GTGTGAAAGGGTTTGTATCA | 5´-spacer-NGG-3´ |
| $P_{cg2014}$_sgRNA-31 | cg2014 | CATGGCTAATTTTCGAAGCA | 5´-spacer-NGG-3´ |
| $P_{cg2014}$_crRNA-1 | cg2014 | TTAATTTAAAAGTATATTTC | 5´-TTTN-spacer-3´ |
| $P_{cg2014}$_crRNA-2 | cg2014 | AAAGTATATTTCAGCAATGG | 5´-TTTN-spacer-3´ |
| $P_{cg2014}$_crRNA-3 | cg2014 | CCATTGCTGAAATATACTTT | 5´-TTTN-spacer-3´ |

| | | | |
|---|---|---|---|
| $P_{cg2014}$_crRNA-4 | cg2014 | AGCAATGGTAAATGAATCTC | 5´-TTTN-spacer-3´ |
| $P_{cg2014}$_crRNA-6 | cg2014 | AAGAAAAGTATTGCATTTGT | 5´-TTTN-spacer-3´ |
| $P_{cg2014}$_crRNA-7 | cg2014 | ACAAGGTGATTATGCTTCAC | 5´-TTTN-spacer-3´ |
| $P_{cg2032}$_sgRNA-1 | cg2032 | AATTCAAAGTTATTTTAGAA | 5´-spacer-NGG-3´ |
| $P_{cg2032}$_sgRNA-2 | cg2032 | ATTCTAAAATAACTTTGAAT | 5´-spacer-NGG-3´ |
| $P_{cg2065}$_sgRNA-1 | cg2065 | ATTAATATTTTATTAACTTA | 5´-spacer-NGG-3´ |
| $P_{vnz04460}$_sgRNA | *vnz_04460* | AAGGAACCGGACAGTTGAAC | 5´-spacer-NGG-3´ |
| *vnz04465*_sgRNA | *vnz_04465* | TTTCATGACGACCCCGGACG | 5´-spacer-NGG-3´ |

### FIGURES SUPPLEMENTARY MATERIAL

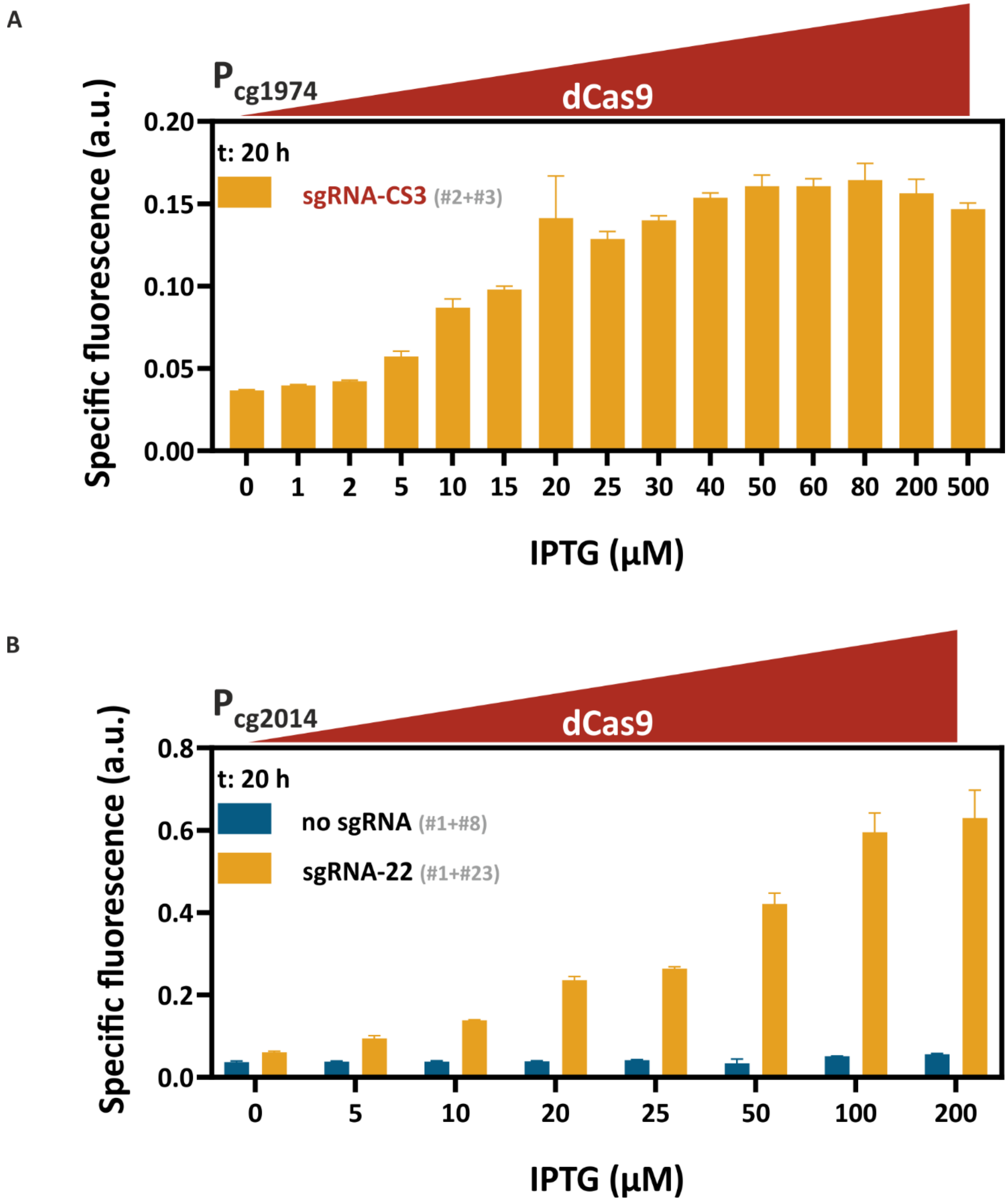


**Figure S1. Correlation of CRISPRcosi efficiencies at prophage promoter $P_{cg1974}$ (A) and $P_{cg2014}$ (B) and *dcas9* expression levels.** Systematic dose-response analysis with increasing amounts of IPTG to induce LacI/$P_{tac}$-controlled *dcas9* expression. Bar plots represent backscatter-normalized reporter outputs (t: 20 h; specific Venus fluorescence) of prophage-free, *cgpS* expressing *C. glutamicum* strains (Δphage::$P_{cgpS}$-*cgpS*, (5)) that were cultivated in CGXII minimal medium supplemented with 111 mM glucose, 25 µg/ml kanamycin, 10 µg/ml chloramphenicol and varying IPTG concentrations in the BioLector I® microtiter cultivation system (n=3). (A) Cells harboured plasmids #3 and #2, which provided the $P_{cg1974}$-*venus* reporter system and allowed for constitutive cg1974_sgRNA-CS3 as well as IPTG-inducible *dcas9* expression. (B) Cells harboured plasmid #1 allowing for IPTG-inducible *dcas9* expression and plasmid #23 or #8, which provided the $P_{cg2014}$-*venus* reporter system with or without cg2014_sgRNA-22, respectively. Plasmid IDs (#number) refer to Table S2.

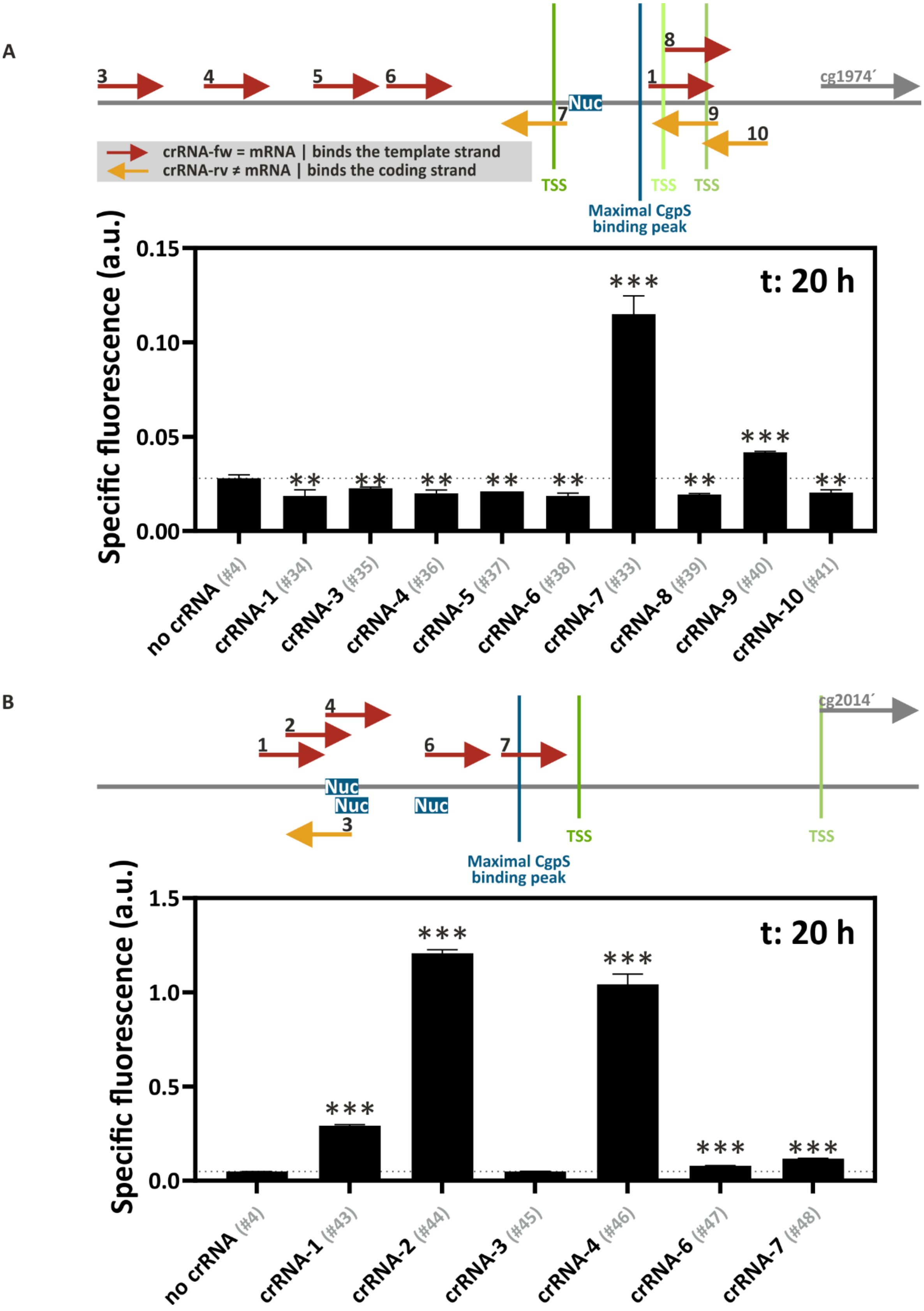
A
3
4
5
6
8
1
7
9
10
Nuc
cg1974´
crRNA-fw = mRNA | binds the template strand
crRNA-rv ≠ mRNA | binds the coding strand
TSS
TSS
TSS
Maximal CgpS binding peak
Specific fluorescence (a.u.)
t: 20 h
no crRNA (#4)
crRNA-1 (#34)
crRNA-3 (#35)
crRNA-4 (#36)
crRNA-5 (#37)
crRNA-6 (#38)
crRNA-7 (#33)
crRNA-8 (#39)
crRNA-9 (#40)
crRNA-10 (#41)
B
1
2
3
4
6
7
Nuc
Nuc
Nuc
cg2014´
TSS
TSS
Maximal CgpS binding peak
Specific fluorescence (a.u.)
t: 20 h
no crRNA (#4)
crRNA-1 (#43)
crRNA-2 (#44)
crRNA-3 (#45)
crRNA-4 (#46)
crRNA-6 (#47)
crRNA-7 (#48)

**Figure S2. The crRNA binding positions determine the dCas12a-mediated CRISPRcosi efficiency.** Arrows in the graphical representations show the target position and DNA strand of the different crRNAs designed for dCas12a guiding to $P_{cg1974}$ (A) and $P_{cg2014}$ (B). The first 30 nucleotides of the corresponding cg1974 and cg2014 genes are shown as grey arrows and the positions of maximal CgpS binding are given as blue vertical lines, while the vertical lines in shades of green indicate the positions of the previously identified TSS positions (TSS colors were mapped according to the ranking of their enrichment scores: dark green > light green) (5). Bar plots represent backscatter-normalized reporter outputs (t: 20 h; specific Venus fluorescence) of prophage-free, *cgpS* expressing *C. glutamicum* strains (Δphage::$P_{cgpS}$-*cgpS*, (5)) harbouring one of the different pEC plasmids containing no (#4) or one of the crRNA encoding sequences ((A): $P_{cg1974}$_crRNA-1 (#34), -3(#35), -4 (#36), -5, (#37), -6 (#38), -7 (#33), -8 (#39), -9 (#40), -10 (#41); (B): $P_{cg2014}$_crRNA-1 (#43),-2 (#44),-3 (#45),-4(#46),-6 (#47), -7 (#48)) in combination with the pJC1-based plasmids #32 (A) or #42 (B) that provided the phage promoter-*venus* reporter system and allowed for the constitutive *dcas12a* expression. The reporter levels of cells lacking any crRNA are given as dotted grey horizontal lines. Number of asterisks above bars indicate the significance levels of crRNA effects (unpaired t test, ***two-tailed *p*-value<0.005, **two-tailed *p*-value<0.05). Strains were cultivated in the BioLector I® microtiter cultivation system in CGXII minimal medium containing 111 mM glucose, 25 µg/ml kanamycin and 10 µg/ml chloramphenicol (n=3). Plasmid IDs (#number) refer to Table S2.

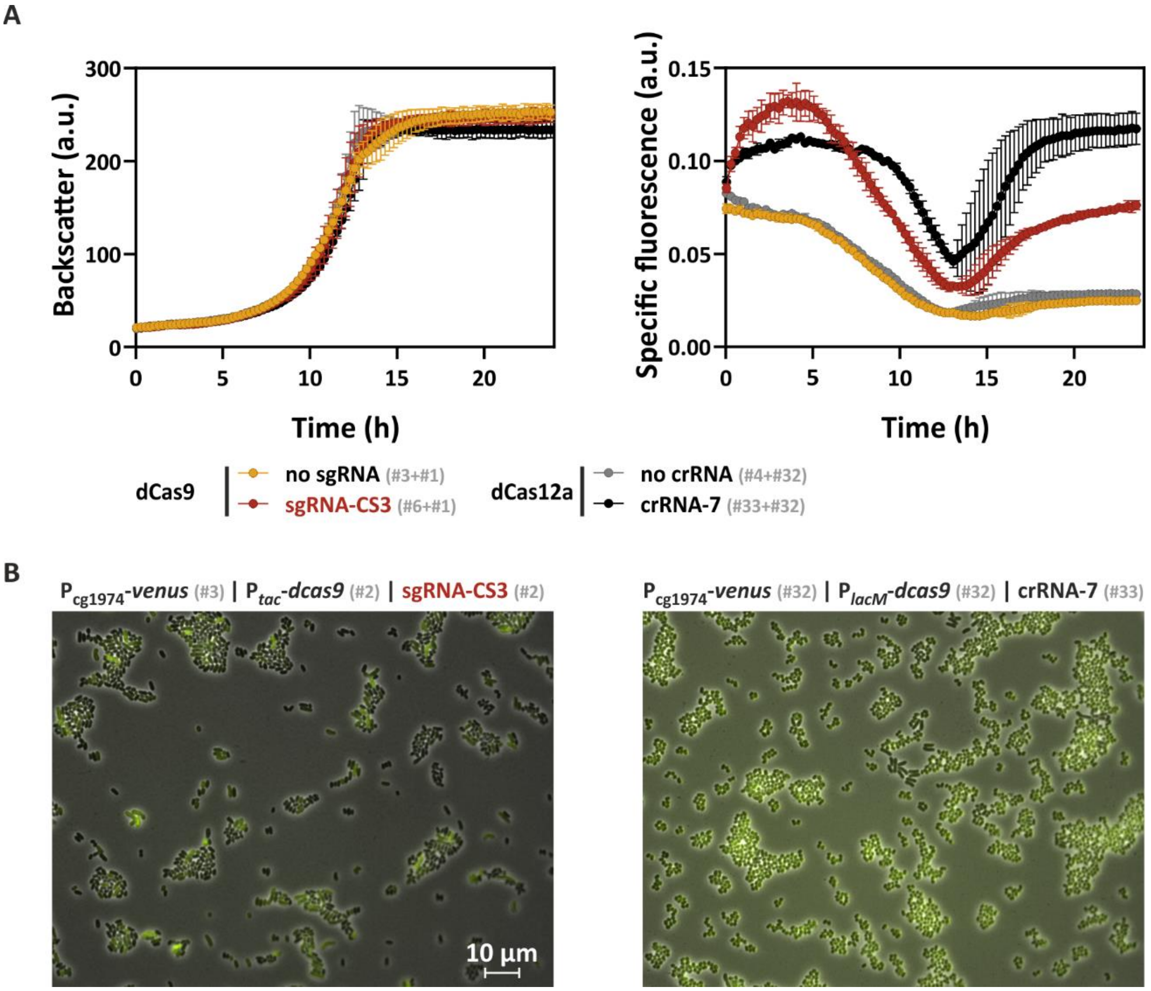


**Figure S3. Supplementary information on Figure 5: dCas9 and dCas12a both mediate efficient CRISPRcosi.** (A) The graphs represent the backscatter values and the time-resolved, backscatter-normalized reporter outputs (specific Venus fluorescence) of prophage-free, *cgpS* expressing *C. glutamicum* strains (Δphage::$P_{cgpS}$-*cgpS*, (5)) harbouring the $P_{cg1974}$-*venus* reporter construct and either the *dcas9* gene combined with the sgRNA-CS3-encoding sequence (#6+#1) or the *dcas12a* gene in concert with the crRNA-7 encoding sequence (#33+#32). Plasmid combinations lacking any guide RNAs encoding sequences (dCas9: #3+#1; dCas12a: #4+#32) served as controls. Strains were cultivated in the BioLector I® microtiter cultivation system in CGXII minimal medium containing 111 mM glucose, 25 µg/ml kanamycin, 10 µg/ml chloramphenicol and 200 µM IPTG (n=3). The drop in specific Venus fluorescence, which started after about five h of cultivation, is characteristic of fast-growing *C. glutamicum* cells and is caused by oxygen limitation leading to delayed oxygen-dependent maturation of the Venus protein. After entering the stationary phase, dissolved oxygen availability increased, and fluorescence signals normalized. (B) Representative fluorescence microscopy images of prophage-free, *cgpS* expressing *C. glutamicum* strains (Δphage::$P_{cgpS}$-*cgpS*, (5)) harbouring the $P_{cg1974}$-*venus* reporter construct and either the *dcas9* gene combined with the sgRNA-CS3-encoding sequence (#3+#2) or the *dcas12a* gene in concert with the crRNA-7 encoding sequence (#32+#33). The scale bar represents 10 µm. Plasmid IDs (#number) refer to Table S2.

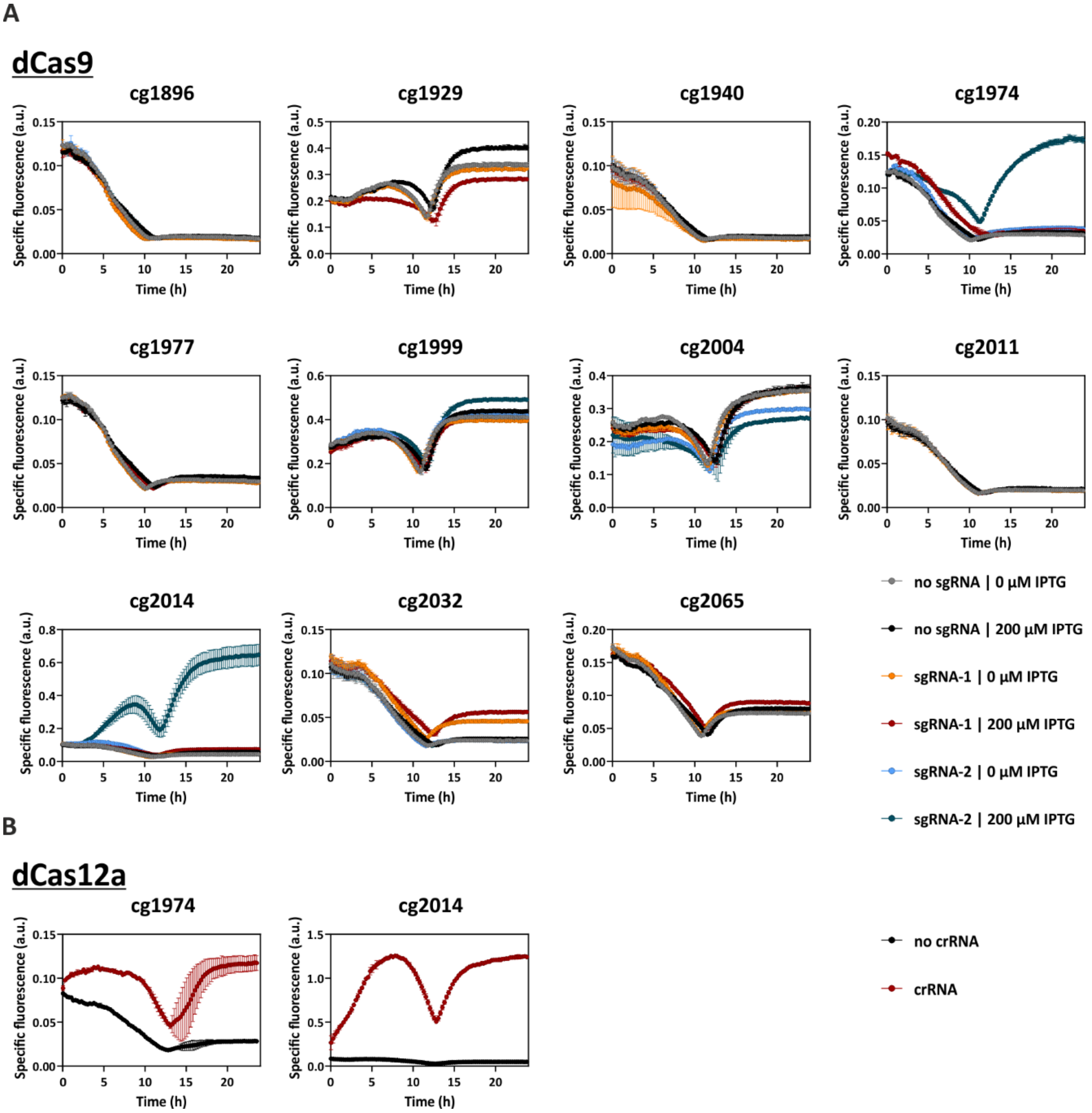


**Figure S4. Supplementary information on Figure 6: CRISPRcosi counteracts CgpS-mediated silencing at different prophage promoters.** Time-resolved representation of the backscatter-normalized specific Venus fluorescence values of all different strains shown in Figure 6. Graphs represent backscatter-normalized reporter outputs (specific Venus fluorescence) in the course of cultivation of cells with and without guide RNA encoding sequences. (A) Prophage-free, *cgpS* and *dcas9* co-expressing *C. glutamicum* cells (Δphage::$P_{cgpS}$-*cgpS* (5)) harbouring the *dcas9* encoding, sgRNA-free plasmid #1 served as platform strain. Guide RNA-free strains were equipped with pJC1-plasmids containing the sequences of the different prophage promoters fused to the reporter gene *venus* to monitor promoter activities ($P_{cg1896}$: #49; $P_{cg1929}$: #50; $P_{cg1940}$: #51; $P_{cg1974}$: #3; $P_{cg1977}$: #52; $P_{cg1999}$: #53; $P_{cg2004}$: #54; $P_{cg2011}$: #55; $P_{cg2014}$: #8; $P_{cg2032}$: #56; $P_{cg2065}$: #57). $P_{J23319}$-mediated expression of specific guide RNAs was achieved by inserting the sgRNA encoding sequences into the pJC1-plasmids ($P_{cg1896}$: sgRNA-1 (#58) and -2 (#59); $P_{cg1929}$: sgRNA-1 (#60); $P_{cg1940}$: sgRNA-1 (#61) and -2 (#62); $P_{cg1974}$: sgRNA-CS1 (#13) and -CS3 (#6); $P_{cg1977}$: sgRNA-1 (#64); $P_{cg1999}$: sgRNA-1 (#65) and -2 (#66); $P_{cg2004}$: sgRNA-1 (#67) and -2 (#68); $P_{cg2011}$: sgRNA-1 (#69); $P_{cg2014}$: sgRNA-1 (#70) and -22 (#23), $P_{cg2032}$: sgRNA-1 (#71) and -2 (#72); $P_{cg2065}$: sgRNA-1 (#73)). (B) Prophage-free, *cgpS* expressing *C. glutamicum* cells (Δphage::$P_{cgpS}$-*cgpS* (5)) harboured the pJC1-based plasmids containing the *dcas12a* gene and the phage promoter fused to the reporter gene *venus*. crRNA-free plasmid #4 served as control, while plasmids #33 ($P_{cg1974_}$crRNA-7) and #44 ($P_{cg2014_}$crRNA-2) were used for $P_{J23319}$-mediated expression of specific crRNAs. The drop in specific Venus fluorescence is caused by a delayed oxygen-dependent maturation of the Venus protein during exponential growth. Fluorescence signals normalize after cells have entered the stationary phase and the levels of dissolved oxygen increased. This effect does not affect the 20 h measurement provided in the main text, which allows for a fair comparison. Strains were cultivated in the BioLector I® microtiter cultivation system in CGXII minimal medium containing 111 mM glucose, 25 µg/ml kanamycin, 10 µg/ml chloramphenicol and 0 or 200 µM IPTG (n=3). Pplasmid IDs (#number) refer to Table S2.

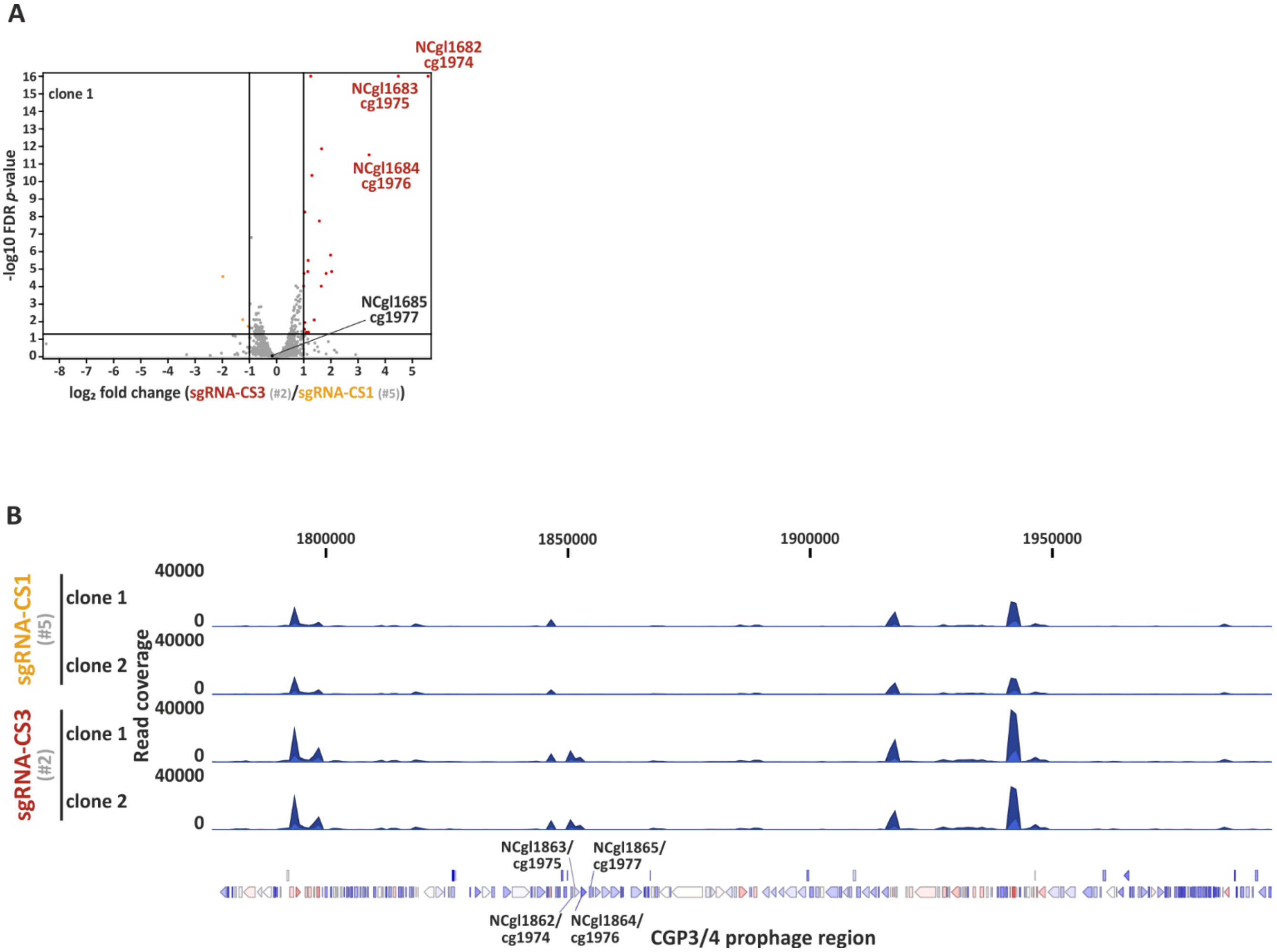


**Figure S5. Supplementary data on Figure 7: Investigation of potential off-target effects of CRISPRcosi.** Analysis of gene expression levels of *C. glutamicum* wild-type cells harbouring the pEC-based plasmids that allowed for IPTG-inducible *dcas9* expression and constitutive expression of sgRNA-CS3 (#2) in comparison to control cells expressing sgRNA-CS1 (#5). Both strains were cultivated in duplicates in CGXII minimal medium supplemented with 111 mM glucose, 10 µg/ml chloramphenicol and 200 µm IPTG for 7 h. (A) Red and orange dots in the volcano plot represent significantly up- and downregulated genes (FDR-*p*-values<0.05 and $|\log_2$ fold change$|>1$), respectively, in cells expressing sgRNA-CS3 (clone 1) in comparison to cells expressing sgRNA-CS-1 (clone 1+2). Non-significant hits are given as grey dots (FDR-*p*-values>0.05 or $|\log_2$ fold change$|<1$), while the corresponding significance thresholds are represented by black lines. (B) RNA-seq read coverage profiles plotted over the CGP3/4 prophage region of clones expressing sgRNA-CS1 and sgRNA-CS3. Coverage data aggregation was set to over 100 bp. Prophage genes are shown as arrows below the plots, highlighted are genes cg1974-77 (NCgl1682-85). Plasmid IDs (#number) refer to Table S2.

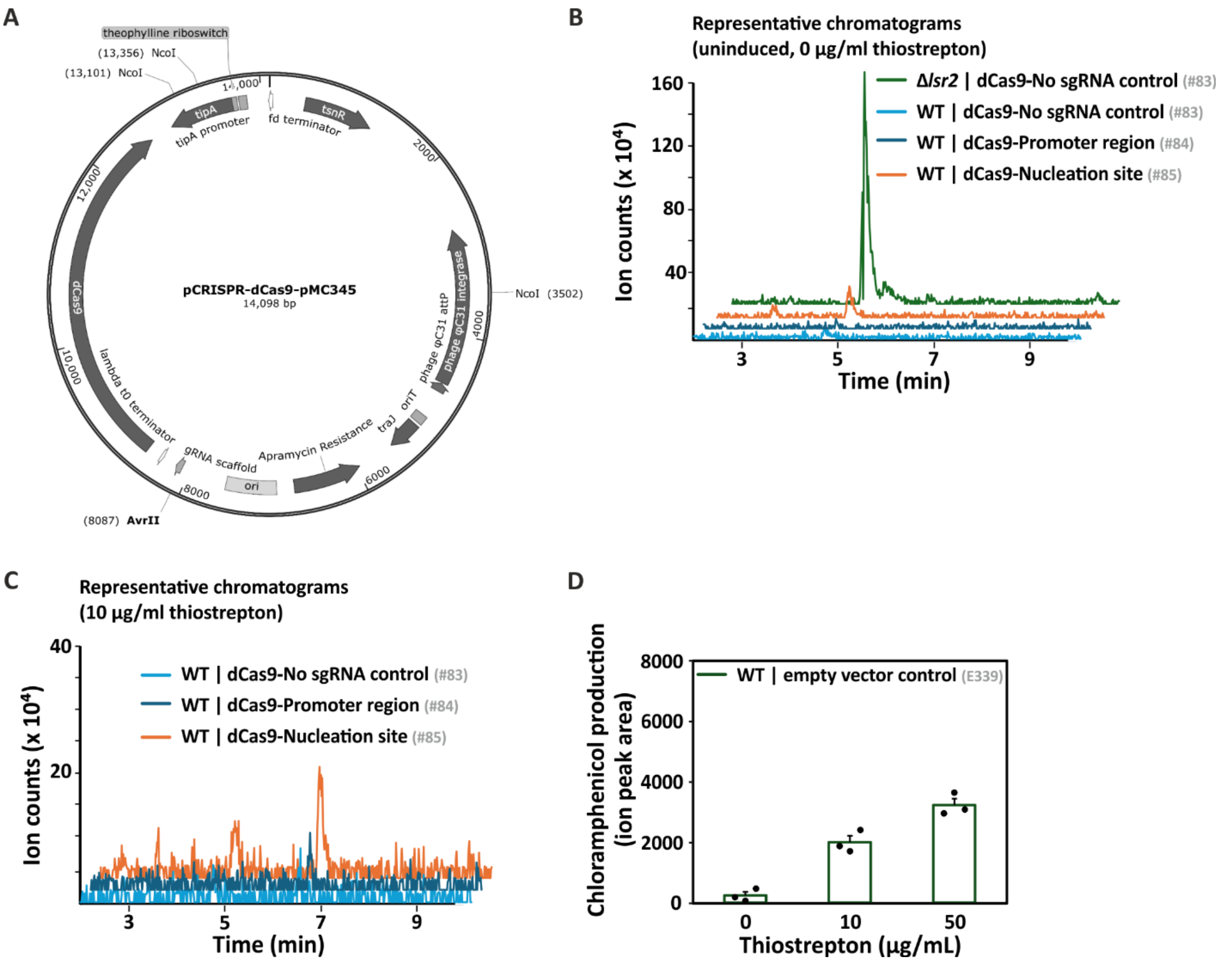


**Figure S6. Supplementary data on Figure 9: Application of CRISPRcosi to increase antibiotic production by *Streptomyces*.** (A) Map of an integrative and inducible dCas9 plasmid, derived from pCRISPR-dCas9 (Addgene plasmid #125687, #AT, (15)). The original NcoI sgRNA cloning site was altered to generate a unique AvrII site. (B) Representative ion chromatograms of chloramphenicol ($[M-H]^-$ at *m/z* 321.00). The input samples were metabolites extracted from non-induced (0 µg/ml thiostrepton) *S. venezuelae* wild-type (WT) carrying empty dCas9 plasmid control (no sgRNA, light blue, #83), sgRNA-vnz04460 (promoter-targeting, dark blue, #84), sgRNA-vnz04465 (nucleation site-targeting, orange,

#85), or a Δ*lsr2* mutant carrying the empty plasmid as a positive uninduced control (green, #83). (C) Similar to (B) except the samples were from the same dCas9 construct-carrying *S. venezuelae* wild-type strains grown under thiostrepton induction (10 µg/mL). Chloramphenicol was eluted at 4.75 min under the gradient used in the mobile phase, shown by major peaks close to the 5 min mark in the figures. (D) Bar plots represent chloramphenicol production by wild-type *S. venezuelae* carrying empty plasmid control E339 (pIJ6902-hyg: no dCas9, so sgRNA, thiostrepton resistant). Thiostrepton (10 or 50 µg/mL) was added to the culture, after which chloramphenicol production was assessed, as described for Fig. 9. Error bars indicate standard error of three biological replicates of MYM liquid cultures grown for 3 days. The value of each replicate is depicted as a black filled circle. These results showed that thiostrepton stimulated the chloramphenicol production in *S. venezuelae*, but not to the same extend as observed with CRISPRcosi (Figure 9B). Plasmid IDs (#number) refer to Table S2.
